# Near-atomistic simulations reveal the molecular principles that control chromatin structure and phase separation

**DOI:** 10.1101/2025.11.17.688899

**Authors:** Kieran Russell, Yifang Chen, Jorge R. Espinosa, David Farré-Gil, Huabin Zhou, Maria Julia Maristany, Jose Ignacio Perez-Lopez, Jan Huertas, Modesto Orozco, Michael K. Rosen, Rosana Collepardo-Guevara

## Abstract

Understanding how chromatin’s physicochemical properties shape its emergent organisation is central to deciphering genome function. To address this, we present OpenCGChromatin, a high-performance coarse-grained model that achieves near-atomistic simulations of chromatin systems an order of magnitude larger than previously possible, spanning biomolecular condensates and fibers tens of kilobases in length. OpenCGChromatin simulations independently predict, from physicochemical principles, the linker-DNA-dependent chromatin structures observed by cryo-ET and the relative thermodynamic stability of condensates inferred from biochemical assays. Crucially, OpenCGChromatin resolves histone-tail dynamics and interaction networks that remain inaccessible experimentally, explaining how linker-DNA length controls histone tail accessibility and the resulting multiscale structure of chromatin condensates. Extending simulations to 108-nucleosome fibers shows that acetylation disrupts chromatin compaction in a pattern-specific manner by weakening key tail-mediated interactions, with H4K16 and H3K9 emerging as the most energetically disruptive modifications. These results position OpenCGChromatin as a powerful framework for linking molecular detail to emergent chromatin organization.

## I. INTRODUCTION

Chromatin organization is fundamental to genome function. Its hierarchical structure both influences and responds to gene expression, DNA repair, and other cellular processes [1, 2]. The basic unit of chromatin is the nucleosome, consisting of approximately 147 bp of DNA wrapped around a histone octamer [3, 4]. Nucleosomes possess ten intrinsically disordered protein (IDP) regions (histone tails) stemming out from the histone core. These tails are positively charged and mediate interactions with neighboring nucleosomes, DNA, and other proteins [4]. Consecutive nucleosomes are connected by DNA linkers of variable length (typically *∼*20–80 bp) and sequence.

In cells, nucleosomes are chemically heterogeneous, carrying distinct post-translational modifications—particularly within the histone tails—and recruiting diverse regulatory proteins according to genomic context, cell type, and cell-cycle stage [5]. Linker DNA length [6–9], irregular nucleosome spacing [10, 11], nucleosome-free regions [11– 14], linker histones [15, 16], and the distributions of post-translational modifications [17, 18] can all profoundly alter the nanoscale structure of chromatin. Consistently, chromatin organization from the nanoscale to whole-nucleus scales is highly heterogeneous and dynamic [13, 19–25].

Over the past decade, transformative experiments have shown that chromatin [26, 27], together with a growing list of intranuclear proteins and RNAs, can undergo phase separation to form condensed-phase droplets termed biomolecular condensates. This has motivated a view of the nucleus as an emulsion of functionally diverse biomolecular condensates, each forming specific microenvironments (distinct sets of biomolecules, metabolites, and thermodynamic parameters) that compartmentalize chromatin, control chemical reactions, and contribute to genetic function and regulation. Examples include nucleoli, Cajal bodies, and nuclear speckles [28], as well as transcriptional condensates containing transcription factors, co-factors, chromatin regulators, non-coding RNAs, and RNA polymerase II [29–34].

In the nucleus, chromatin phase separation emerges within a crowded, multicomponent environment. Here, multidomain and disordered proteins, nucleic acids, and nucleosomes form transient multivalent interactions that, under certain conditions, give rise to condensates sustained by interconnected interaction networks [9, 35–40]. The molecular features that regulate binding among nucleosomes, and between nucleosomes and proteins (e.g., chemical composition, mechanical properties, and local environment), are therefore key determinants of this process. In particular, the structure of individual chromatin fibers tunes collective phase behavior [8, 9, 26]. Linker lengths close to integer multiples of the DNA helical repeat plus 5 bp (i.e., 10*N*+5 bp, where *N* is an integer) promote disordered chromatin configurations that enhance multivalent interactions and favor phase separation, whereas linkers at integer multiples (i.e., 10*N* bp) form more ordered structures that suppress it [8, 9, 26, 27]. The disordered organization of nucleosomes—facilitated by nucleosome breathing—further amplifies chromatin multivalency, increasing the density of molecular connections within the condensed chromatin liquid network and reinforcing the stability of its condensates [25].

Understanding how chromatin conformational dynamics, and the ensuing formation and breakage of intermolecular interactions, regulates the properties of chromatin condensates is crucial for uncovering the physical principles that organize the genome and regulate its function. Yet, capturing these processes across the relevant molecular and mesoscale regimes remains a major challenge. State-of-the-art cryo-ET can now visualize chromatin organization inside condensates at nucleosome resolution [9], but it cannot directly resolve the dynamical behavior of individual biomolecules and quantify the energetic determinants of phase separation. Conversely, existing computational models either neglect chemical specificity or lack the resolution and efficiency needed to connect residue-level interactions with mesoscale organization [24]. To bridge this gap, computational frameworks must simultaneously capture the chemical detail of histones and DNA and the emergent mesoscale behavior of chromatin across phase boundaries. Among available approaches, so-called near-atomistic coarse-grained models— where individual amino acids and DNA base pairs are explicitly represented—offer this potential [25, 41–43]. However, their high computational cost and the rugged energy landscapes of chromatin have so far restricted simulations to small arrays of 3–12 nucleosomes, precluding direct exploration of gene-scale chromatin organization and condensate structure.

Here, we introduce OpenCGChromatin, a high-performance, residue- and nucleotide-resolution coarse-grained model that achieves over an order-of-magnitude acceleration relative to our previous residue-level model [25] while improving physicochemical accuracy and experimental agreement (Fig. 2, Fig. S3). This advance extends the frontiers of near-atomistic simulations to dynamics of hundreds of nucleosomes and biomolecular condensates. The model recapitulates key experimental observations, such as the oscillatory dependence of chromatin compaction on linker DNA length [6, 7], the structural motifs resolved by cryo-EM [44], X-ray crystallography [45] and cryo-ET [9], and the modulation of chromatin condensate stability with linker DNA length [8, 9]. OpenCGChromatin provides access to the dynamical configurational ensembles of chromatin, and its histone tails, across phase boundaries at near-atomistic resolution, revealing the molecular and thermodynamic mechanisms by which phase separation emerges and is regulated. In doing so, it bridges chemical-specific molecular interactions with mesoscale structure and thermodynamics.

Using OpenCGChromatin, we find that chromatin phase separation arises from the interplay of linker DNA length and mechanics, anisotropic internucleosome interactions, histone-tail dynamics, and chemical specificity. Linkers close to 10*N*+5 bp introduce single-fiber structural frustration that prevents regular nucleosome stacking, limits fiber compaction, and increases chromatin multivalency, thereby promoting phase separation. In contrast, 10*N* bp linkers favor compact zigzag conformations that suppress inter-fiber connectivity and disfavor phase separation.

Our work reveals that chromatin can be conceptualised as a semi-flexible stickers-and-spacers polymer [46], where the stickers correspond to nucleosome faces and histone tails, and the spacers to the linker-DNA segments that separate them. Critically, because the spacers are double-helical DNA with defined twist and bending mechanics, chromatin phase behaviour depends not only on the stickers-and-spacers composition but also sensitively on the length, geometry, and mechanical properties of the linker-DNA spacers. These linker-DNA features dictate nucleosome orientations and histone-tail accessibility and thereby either promote or suppress chromatin phase separation. OpenCGChromatin thus provides a physically grounded, near-atomistic description of the mechanisms regulating chromatin structure and phase behaviour, establishing a molecular foundation for understanding how phase separation contributes to chromatin structure.

## II. RESULTS

### A. The OpenCGChromatin model

We introduce OpenCGChromatin, a high-performance coarse-grained chromatin model that bridges atomistic detail and mesoscale organization, enabling simulations of chromatin arrays spanning tens of kilobases and condensates containing hundreds of nucleosomes at near-atomistic resolution.

OpenCGChromatin constitutes a complete GPU-compatible reformulation of our previous CPU-only chemically specific chromatin coarse-grained model, which represented histones at residue resolution and DNA at the base-pair-step level using ellipsoids. The new framework uses a fully particle-based description of histones and DNA, with one bead per amino acid (centered at the C_*α*_) and one per nucleotide (centered at the C1^*′*^ position) plus three virtual sites per base pair, while treating solvent and ions implicitly [25, 47]. The entire interaction scheme was redesigned and reparameterized using a hybrid bottom-up and top-down approach. At the bottom-up level, bead sizes, charges, interaction terms, configurational potentials, and nucleosome configurations were derived from atomistic structures and simulations [4, 18, 25, 47] to preserve essential chemical and structural features of chromatin components. At the top-down level, the energetic balance between electrostatic and hydrophobic protein–protein, protein–DNA and DNA–DNA interactions was tuned to reproduce experimental trends in fiber compaction with respect to linker length [6, 7] (Fig. S4). This comprehensive redesign ensures that OpenCGChromatin reproduces experimental observables across molecular and mesoscale regimes.

Parameters were optimized to capture linker-length-dependent compaction trends from sedimentation experiments [6, 7] for regular, reconstituted arrays of linker lengths 25, 30 and 58 bp (Fig. S4). All other results reported in this work, including sedimentation coefficients for other linker lengths, cryo-ET structural comparisons, condensate properties, and acetylation effects, are predicted by the model.

A central advance of OpenCGChromatin is the replacement of the rigid base pair description of DNA [48, 49]—based on helical coordinates and an ellipsoidal representation—with the state-of-the-art CGeNArate coarse-grained model [47], which employs a fully particle-based Cartesian representation. CGeNArate is a bottom-up coarse-grained model for B-form DNA parametrized from large-scale all-atom molecular dynamics simulations at the tetranucleotide level, while also capturing emergent mechanical properties of long DNA oligomers. In our previous chromatin model [25], representing inter-basepair DNA deformations in helical space required nontrivial coordinate transformations and Jacobians to compute forces. In contrast, CGeNArate treats nucleotides as Cartesian particles whose local deformations are described by low-order, sequence-dependent polynomial functions, enabling seamless integration with OpenMM [50, 51]. In addition, unlike the symmetric rigid base pair model, CGeNArate explicitly resolves the major and minor grooves of B-form DNA, providing a more faithful description of DNA shape, geometry, and flexibility. As a result, CGeNArate accurately reproduces the equilibrium atomistic free-energy landscape of B-form DNA, including twist–bend coupling and sequence-dependent stiffness, and captures the dynamics of diverse DNA constructs, including circular duplexes, kilobase-length fragments, and even complete mitochondrial DNA [47]. Solvent and mobile ions are treated implicitly via Debye–Hückel electrostatic screening, which captures the mean-field effect of monovalent salt concentration on electrostatic interactions without the computational cost of explicit solvent. In OpenCGChromatin, the one-bead-per-nucleotide of CGeNArate is augmented with three additional virtual sites per base pair: two phosphate beads carrying negative charges for electrostatic interactions, and one excluded volume bead positioned at the helix center to prevent unphysical histone tail configurations such as penetration into the DNA helix.

The secondary structure of the globular domains of the histone proteins and the overall architecture of the histone octamer are maintained using elastic network models derived from atomistic simulations of nucleosomes [16], as in our previous work [25]. Histone tails are treated as fully flexible polymers. The geometry of the nucleosomal DNA around the histone protein core is taken from high-resolution crystal structures of nucleosomes [4]. The central 127 bp of nucleosomal DNA are included in the elastic network to maintain single-nucleosome stability, while allowing breathing motions in the terminal 10 bp on each flank [3]. More extensive nucleosome breathing, which may occur in native chromatin arrays [52], could be explored in future work by relaxing or reformulating these restraints. Detailed model architecture, Hamiltonian, parameterization strategy, and comparison to our previous chemical-specific coarse-grained model [25] are provided in Supplementary Information.

To assess the computational efficiency of this GPU-accelerated implementation, we benchmarked OpenCGChromatin on systems ranging from 12 to 108 nucleosomes. The implementation of OpenCGChromatin in OpenMM achieved performance gains exceeding an order of magnitude over previous CPU-based models (Fig. 1b). Benchmarking on both consumer (NVIDIA RTX 4080) and data-center (NVIDIA H100) GPUs revealed that for 108-nucleosome systems (*∼*190,000 particles), a single H100 GPU outperformed a 128-core AMD EPYC 7742 node by more than an order of magnitude (Fig. 1b). Even for smaller 12-nucleosome fiber simulations, we observed 9-fold speedups relative to a full 56-core Intel Cascade Lake CPU node (Fig. 1b). When single simulations do not saturate the GPU, NVIDIA’s Multi-Process Service can be used to run multiple independent simulations concurrently, further increasing total throughput (Fig. 1b). This is enabled by the relatively low VRAM utilisation of a single simulation (Table S3), which means large systems or many copies of a small system fit easily within the memory of modern GPUs. Importantly, these comparisons juxtapose one GPU against many tens or hundreds of CPU cores; the per-core acceleration, and associated reductions in computational cost and energy consumption, are therefore even more striking. Thus, the real-world gains in throughput and resource efficiency for large-scale OpenCGChromatin simulations are substantially greater than even the raw order-of-magnitude improvements suggest.

**FIG. 1.**
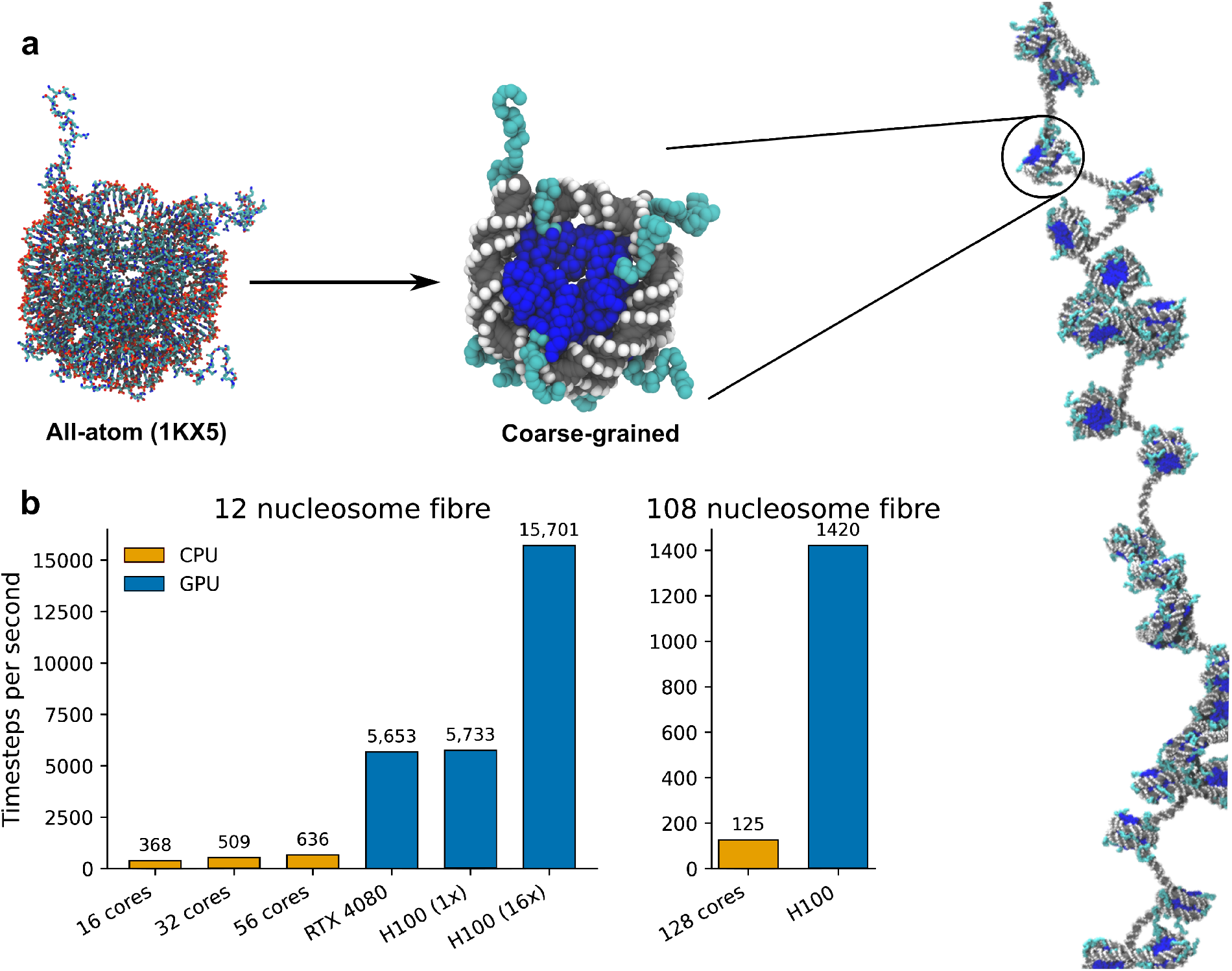
OpenCGChromatin enables large-scale high-resolution simulations of chromatin. (a) Atomistic (left) and coarse-grained (middle) nucleosome renderings, showing histone core (blue), histone tails (cyan), and DNA (grey). At right, a coarse-grained rendering of a chromatin fiber with inset highlighting a single nucleosome. (b) Simulation throughput (timesteps s^−1^). Left: 12-nucleosome fiber (18k particles) benchmarks on an Intel Cascade Lake CPU node; GPUs (RTX 4080, H100) achieve major performance gains, with the “H100 (16×)” case using NVIDIA Multi-Process Service to run 16 replicas simultaneously on a single GPU. Right: 108-nucleosome fiber (190k particles) benchmarks on an AMD EPYC 7742 CPU node; in both small and large systems a single H100 GPU outperforms an entire CPU node by more than an order of magnitude in terms of total throughput, enabling efficient chromatin simulations at unprecedented scale.

**FIG. 2.**
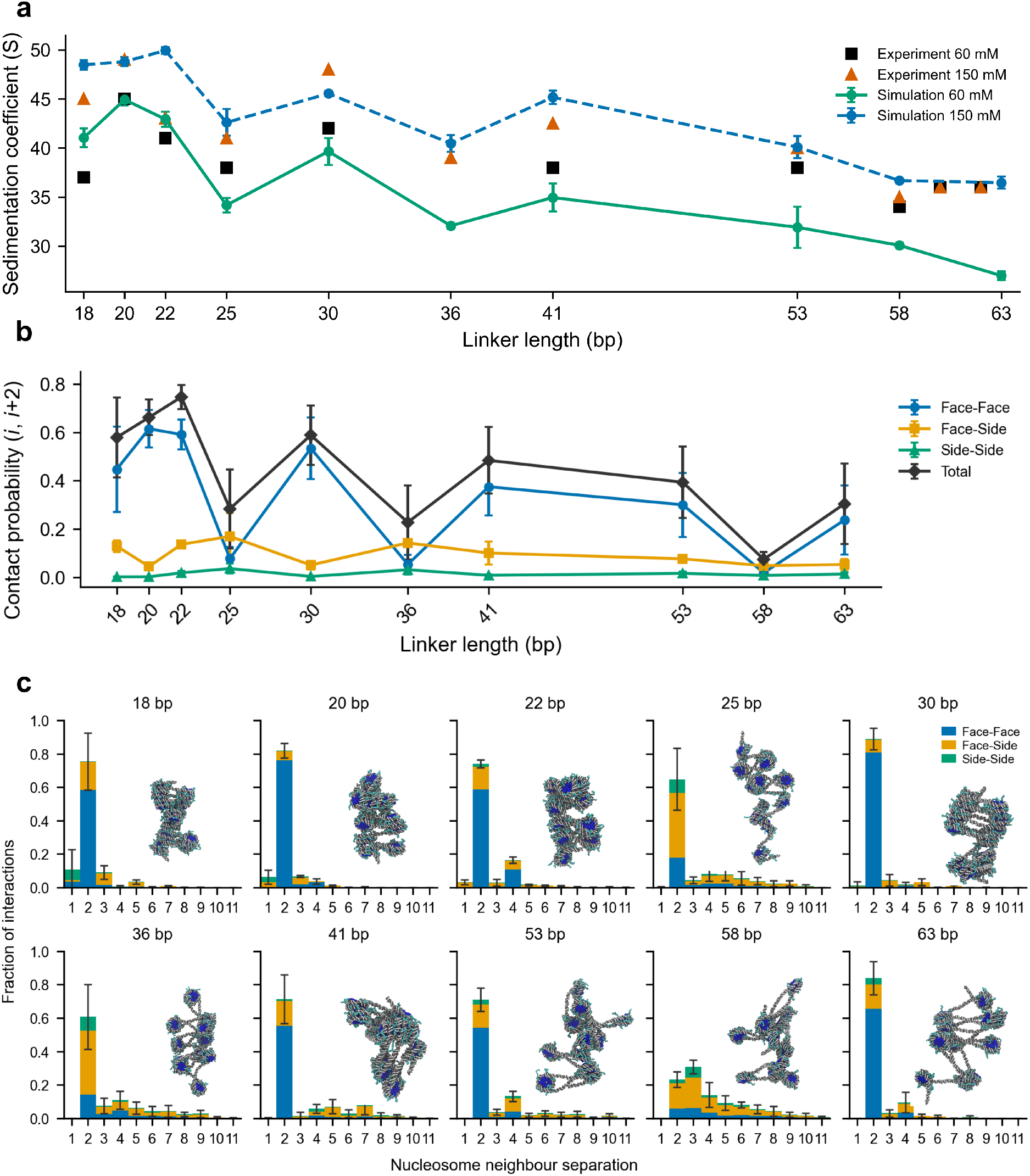
Trends in fiber compaction with respect to linker length. (a) Simulated sedimentation coefficients across linker lengths compared with experimental values [6, 7], revealing an oscillatory trend with enhanced compaction for linker lengths near 10*N* bp and reduced compaction for those near 10*N* + 5 bp. (b) Per-nucleosome contact frequencies for second-nearest neighbor nucleosome–nucleosome interactions across linker lengths. (c) Nucleosome interaction pattern summaries across linker lengths, highlighting the predominance of second-nearest neighbor face–face stacking near 10*N* and increasing heterogeneity at longer linkers. Plotted values represent means over five non-overlapping trajectory blocks; error bars show the standard deviation across block means.

This high-performance model makes it possible to simulate chromatin organization and bulk phase behavior at near-atomistic resolution and scales that were previously unattainable.

### B Model Validation

To validate OpenCGChromatin, we compare its structure and compaction predictions for 12-nucleosome arrays with different regular linker DNA lengths with *in vitro* experiments on reconstituted chromatin fibers, including sedimentation velocity analyses (Fig. 2a) [6, 7], and structures obtained by cryo-EM [44], X-ray crystallography [45] and cryo-ET (Fig. S7) [9].

We simulated ten distinct 12-nucleosome arrays with regular linker lengths equal to 18, 20, 22, 25, 30, 36, 41, 53, 58 and 63 bp. To ensure exhaustive sampling of the rugged free-energy landscape underlying chromatin at high resolution, we employed our Debye-length Hamiltonian replica-exchange (HREX) protocol [25]. In this method, replicas at constant temperature exchange configurations across different values of the Debye length (*λ*_*D*_) governing the screened Coulomb interactions. Fifteen HREX replicas were distributed between *λ*_*D*_ = 8.0 °A (corresponding to 150 mM NaCl) and *λ*_*D*_ = 13.0 °A (corresponding to 60 mM NaCl), achieving an average exchange acceptance of approximately 30 % (Supplementary Information, Fig. S8). This approach is particularly efficient for chromatin systems, requiring only about one-fifth as many replicas as conventional temperature replica exchange, as the dominant free-energy barriers arise from electrostatic interactions [25]. Moreover, because the Debye lengths correspond to experimentally relevant salt concentrations, HREX simultaneously enhances sampling and enables direct probing of the salt-dependent conformational behavior of chromatin.

The sedimentation coefficient (*S*) is widely used as a low-resolution, ensemble-average metric of chromatin compaction, with higher values indicating more compact conformations [6]. Results from the HREX simulations show that OpenCGChromatin achieves near-quantitative agreement with experimental sedimentation coefficients at physiological salt (150 mM). The model accurately captures the characteristic oscillatory dependence of chromatin compaction on linker DNA length [6], where arrays with linker lengths close to 10*N* bp are the most compact and those near 10*N*+5 bp are the least compact (Fig. 2a). This agreement is particularly significant as, to our knowledge, no previous near-atomistic chromatin model, including our original chemically specific model [25] (Supplementary Information, Fig. S3), has reproduced this fundamental experimental trend. In addition, comparing fiber compaction via the radius of gyration, as predicted by the simulations and measured by cryo-ET at 150 mM for 25 bp and 30 bp fibers, shows that all cryo-ET values fall within the simulated distributions (Fig. S7a).

Chromatin compaction emerges from a balance between DNA–DNA electrostatic repulsion, anisotropic nucleosome– nucleosome interactions, and DNA mechanics. This balance depends sensitively on linker length and ionic environment. As the linker DNA becomes longer, the total negative charge of chromatin increases. Therefore, as DNA lengthens, the electrostatic repulsion between adjacent DNA segments begins to dominate over attractive histone-mediated interactions and DNA mechanical constraints. This shift progressively drives chromatin towards decompaction at long linker lengths, regardless of the 10*N* versus 10*N*+5 periodicity. Consistent with this picture, at low ionic strength (60 mM NaCl), diminished electrostatic screening amplifies DNA–DNA repulsion, promoting less compact chromatin states. OpenCGChromatin accurately reproduces this salt-dependent decompaction for short to medium linker DNAs (Fig. 2a). For the longest linkers (*≥*50–63 bp), the experimental sedimentation coefficients decrease more weakly with reduced salt than predicted by our model. This discrepancy may reflect the limited sensitivity of sedimentation velocity assays to highly open, heterogeneous fibers, as well as their susceptibility to small sample variations such as partial nucleosome occupancy. The discrepancy may also arise from the Debye–Hückel treatment of counterion screening in our model, which can overestimate long-range electrostatic repulsion at reduced ionic strength [53] and inherently neglects ion correlation effects. Because the Debye–Hückel potential decays exponentially with a longer screening length at lower salt, the overestimation of repulsion grows as the ionic strength decreases, which is consistent with the discrepancy being most pronounced at 60 mM for the longest linkers. To evaluate the performance of the model in very low salt regimes, we also ran long unbiased MD simulations of the same set of chromatin fibres at 5 mM and compared against experimental sedimentation coefficients [6]. We find that, although the model reproduces the overall qualitative trends, the quantitative agreement with experiment deteriorates at low salt, with simulations progressively underestimating sedimentation coefficients as linker length increases (Fig. S10).

Because nucleosome–nucleosome interactions are intrinsically anisotropic, we characterized chromatin structures across linker lengths by quantifying inter-nucleosome contact frequencies as a function of neighbor order (*i*_*th*_ neighbors) and relative orientation, classifying them into face– face, face–side, and side–side categories. Here, the face denotes the flat nucleosome surface containing the H2A–H2B acidic patch, while the side refers to its cylindrical periphery, perpendicular to the face plane. We have previously shown that face–face stacking yields the strongest inter-nucleosome interactions, followed by face–side and side–side contacts [8, 25]. Hence, face–face contacts are energetically favoured, provided nucleosomes do not need to disturb the orientations imposed by their linker DNAs.

In Fig. 2b, we show the contact probability for different classes of second-nearest-neighbor nucleosome interactions as a function of linker length, highlighting the prevalence of face–face interactions for linker lengths close to 10*N* bp and face–side interactions for those close to 10*N*+5 bp. Face– face and total second-nearest-neighbor contacts display an oscillatory dependence on linker length, diminishing as DNA lengthens mirroring the compaction behaviour in Fig. 2a.

The qualitative linker-length dependence and rank or-dering of contact types in Fig. 2b are preserved across a 12–14 nm range of nucleosome–nucleosome distance cut-offs (Fig. S11). Fig. 2c displays fractions of the total nucleosome–nucleosome interactions in each system, allowing comparison of the relative importance of different contact types. Fig. S9 provides the underlying interaction counts normalized only by the number of frames, enabling comparison of absolute interaction levels across linker lengths.

Our analysis confirms that the oscillatory compaction trend in our simulations arises from a physically consistent mechanism: the helical twist of B-form DNA, which completes one full turn every *∼*10.5–10.7 bp [54], determines the fraction of second-nearest-neighbor face–face stacking interactions that drive compaction (Fig. 2b). Thus, in agreement with experiments [9, 44, 45], linker lengths close to integer multiples of the helical repeat (10*N* bp) position adjacent nucleosomes in phase—with their dyad axes rotated by roughly 0° relative to one another—allowing second-nearest neighbors to align for optimal face–face stacking, yielding the characteristic compact zigzag architecture (Fig. 2c). In contrast, as shown experimentally [9], linker lengths of 10*N*+5 bp correspond to a residual half turn of DNA (a 180° rotational offset between nucleosomes), placing them out of phase and thereby suppressing face–face contacts. When compared to the 10*N* structures, the 10*N*+5 fibers adopt more expanded heterogeneous structures with a greater proportion of face–side contacts (Fig. 2c) [8, 9]. The total number of inter-nucleosome interactions per fiber decreases progressively, consistent with the greater decompaction observed at longer linker lengths (Fig. S9).

To quantitatively compare the structures predicted by OpenCGChromatin with those visualized by cryo-ET for 12-nucleosome arrays with 25 or 30 bp linkers [9], we computed three geometric descriptors of consecutive nucleosome triplets and compared with the distributions reported for cryo-ET fibers in our previous work [9]. These include the center-of-mass distance between second-nearest-neighbors (*D*), and dihedral angles between nucleosome planes *α*_*i,i*+1_ and para_*i,i*+2_ (Fig. S7b). Here, *α*_*i,i*+1_ describes the relative orientation of sequential nucleosome planes and para_*i,i*+2_ reports the orientation of second-nearest-neighbor nucleosome planes (details of calculations in Supplementary Information). The simulated *D* and *α*_*i,i*+1_ closely match those from cryo-ET, while para_*i,i*+2_ shows the correct trend but with a slightly higher mean for 25 bp and a broader distribution for 30 bp linkers. The broader distribution likely reflects the more exhaustive sampling of inter-nucleosome configurations in the simulations compared with cryo-ET. Larger para_*i,i*+2_ values for 25 bp indicate that second-nearest-neighbor nucleosomes are slightly more out of phase in the simulations than in the experiment. This discrepancy could arise from modest over-rigidity in linker-DNA torsion and bending, or overestimation of DNA–DNA electrostatic repulsion under the symmetric Debye–Hückel treatment (as discussed for compaction and in previous work [53]). Because para_*i,i*+2_ depends on the relative orientation of nucleosomes connected by two consecutive linkers, it is more sensi-tive than *D* or *α*_*i,i*+1_ to small variations in linker-DNA twist and bending. Twist deviations accumulate over two linkers, amplifying minor differences in torsional rigidity, helical pitch, or electrostatic interactions. Nonetheless, the overall agreement is excellent. Indeed, consistently with the predictions of OpenCGChromatin, analysis of the cryo-ET internucleosome interactions [9] confirms the dominance of ordered face–face arrangements for 10*N*-like linkers and the prevalence of irregular configurations with face–side interactions for 10*N*+5-like arrays. Given that high-resolution structural data for 10*N*+5 chromatin remain scarce, this agreement provides rare structural insight into the less-characterized, irregular regime.

The 10*N* chromatin structures also align with the more widely available high-resolution molecular data [44, 45], with ordered zigzag configurations matching cryo-EM structures of reconstituted 12-nucleosome arrays (30 bp linker) and the tetranucleosome crystal structure (20 bp linker).

Regardless of the 10*N* vs 10*N*+5 periodicity or linker length, second-nearest neighbor (*i ±* 2) interactions consistently dominate, in agreement with in situ sequencing-based assays—including Micro-C [55, 56], RICC-seq [57], and Hi-CO [58]—which resolve single-nucleosome interactions across variable linkers and reveal preferential *i ±* 2 contacts.

OpenCGChromatin complements experiments by unveiling the molecular basis of the linker-length-dependent structural differences via analysis of residue-level DNA–protein and protein–protein inter-nucleosome contacts (Fig. 3c). Histone tail contact patterns show clear oscillatory trends with linker DNA length, with H4 and H2A N-terminal tails exhibiting the highest contact frequencies at linker lengths near 10*N* (Fig. 3a,b). This behavior is structurally consistent with the surface localization of both tails, which lie on the nucleosome face (Fig. 3d). Thus, in zigzag (10N-like) fibers, the H4 and H2A N tail are optimally positioned to mediate face–face contacts between nucleosomes.

**FIG. 3.**
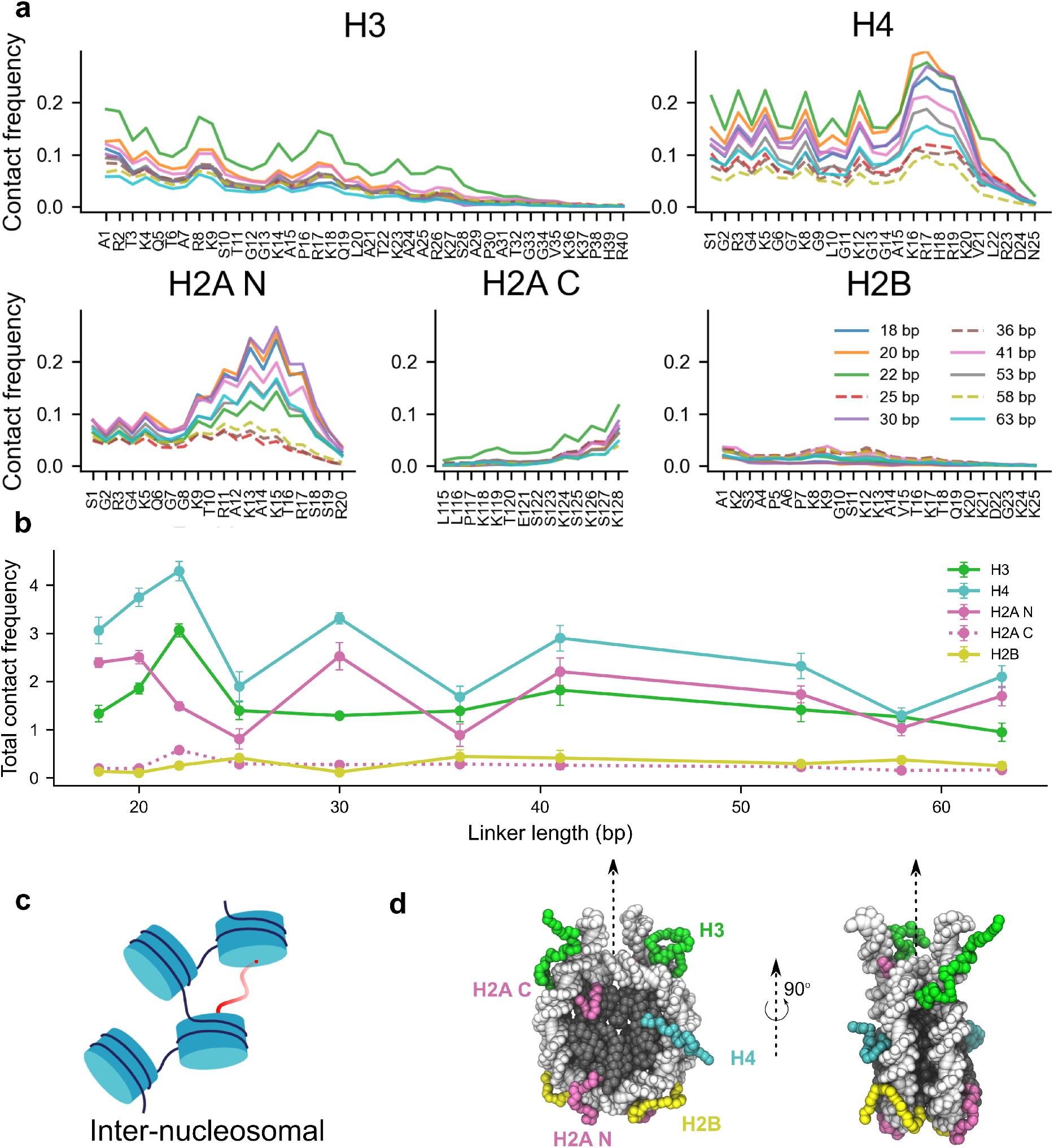
Histone tail contact patterns from short fibers. (a) Line plots showing the contact frequency of each histone tail with other nucleosomes across all simulated fibers, revealing enriched H4/H2A contacts for linker lengths close to 10*N* (solid lines) as compared to those close to 10*N*+5 (dashed lines). (b) Sum of contact frequencies for each of the five histone tails across linker lengths. Plotted values represent means over five non-overlapping trajectory blocks; error bars show the standard deviation across block means. (c) Schematic of an inter-nucleosomal, or non-parental, tail-mediated contact [59]. (d) Diagram showing the positions of each histone tail on the nucleosome.

At the residue level, the H4 basic patch (K16–K20) is the dominant interaction hotspot, followed by the basic patch of H2AN (R11–R17), which mediate frequent electrostatic interactions with both DNA and the H2A– H2B acidic patch of other nucleosomes across all 10*N* linker lengths. Consistent with these findings, electron microscopy and single-molecule force spectroscopy experiments on H4-tail cross-linked chromatin [60, 61] show that folding of zigzag fibers is stabilized by face–face interactions between the H4 tail of one nucleosome and the H2A– H2B surface of another. The central role of the H4 basic patch is further supported by the well-established decompaction triggered by H4K16 acetylation [18, 62], and by the observation that reversible acetylation at this residue— one of the most frequent post-translational modifications across eukaryotes—has widespread regulatory consequences for chromatin structure and function [63]. This mechanistic picture aligns with extensive biochemical and biophysical evidence demonstrating that the H4 tail, and specifically its basic patch, is essential for compaction of zigzag fibers [64, 65]. Finally, the enrichment of H2A tail–DNA contacts observed in our simulations mirrors cryo-EM evidence in 12-nucleosome arrays [44] as well as results from biochemical assays, which have identified the important role of the H2A tail in the compaction [66] and oligomerization [67] of chromatin. In particular, the cation-rich patch from R11–R17 has high contact frequencies, peaking around K13 and K15, particularly for linker lengths close to 10*N*. We find that by evolving structures under the OpenCGChromatin Hamiltonian alone, our unbiased simulations arrive at conclusions similar to those from simulations explicitly biased toward experimentally observed cryo-ET chromatin structures [9], indicating similar underlying nucleosome configurations in both cases. To distinguish whether histone tails preferentially interact with their own nucleosome or with neighbouring nucleosomes, we decomposed residue-level tail contacts into parental (intra-nucleosomal) and non-parental (inter-nucleosomal) contributions (Fig. S12). The parental contribution is largely independent of linker length, whereas the linker-length dependence described in this section arises predominantly from non-parental inter-nucleosomal interactions (Fig. S12). As a robustness check, we also confirmed that the relative ordering of tail contact frequency is unaffected by reasonable variation of the residue-level contact-distance cutoff (Fig. S13).

These comparisons demonstrate that OpenCGChromatin quantitatively reproduces the modulation of chromatin compaction and structure with linker length, validating it as a predictive model for probing chromatin structure across multiple scales.

## C. Physical and molecular mechanisms of chromatin phase separation

We investigated the phase behavior of chromatin by performing direct coexistence molecular dynamics simulations of chromatin solutions at residue and nucleotide resolution. We simulated two types of systems, each comprising 81 tetranucleosome fibers with either 25 or 30 bp linker lengths, at room temperature and 100 mM NaCl. Using the direct coexistence method, we simulated the condensed phase in equilibrium with the dilute phases in the same elongated simulation box, separated by an interface [68]. This setup allows the two phases to freely exchange molecules and energy until equilibrium is reached, enabling the equilibrium coexistence densities to be obtained directly without additional thermodynamic assumptions or biasing potentials. As a result, our unbiased simulations predict, from physicochemical principles, the equilibrium structures of chromatin within condensates, as well as the inter-nucleosome interactions, and the dynamical behavior of histone tails, which sustain these condensates at their equilibrium coexistence densities. We note that the direct coexistence approach employed here probes the effective equilibrium thermodynamic behaviour of chromatin condensates. As such, the present framework does not explicitly account for non-equilibrium ATP-driven processes, including chromatin remodeling, transcription, or loop extrusion.

OpenCGChromatin independently predicts that the 25 bp chromatin solution phase-separates into a chromatin-rich condensate with sharp interfaces coexisting with a chromatin-poor dilute phase, whereas the 30 bp system lies near the phase boundary with only weak phase separation (diffuse clusters and no stable interface; Fig. 4a,b). These predictions are in agreement with experiments [8, 9, 26, 27] and with our phase-separation assay at 100 mM salt (Fig. 4e,f).

**FIG. 4.**
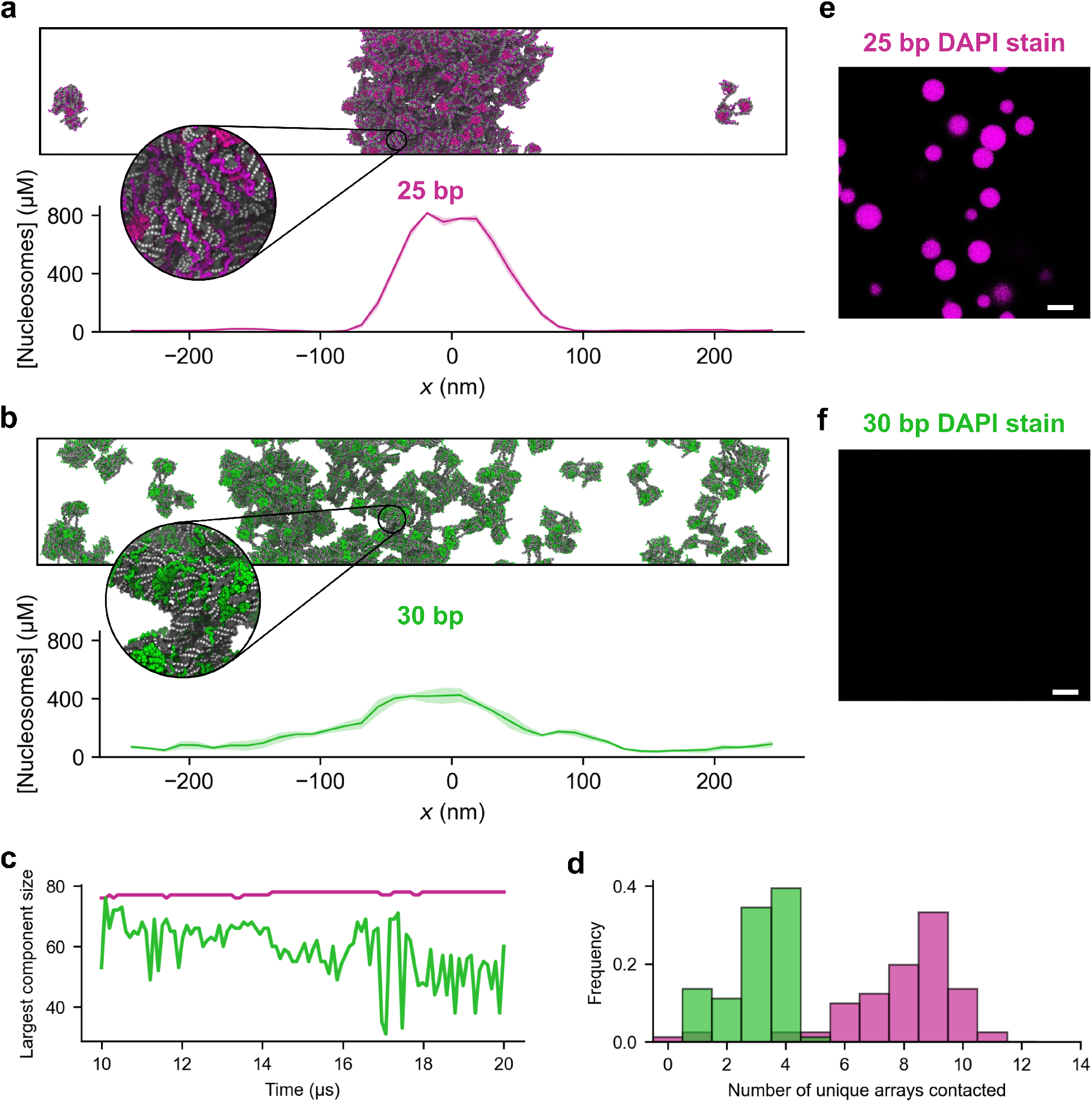
Linker length modulates phase separation of chromatin arrays. (a) Simulation snapshot and density profile of 25 bp chromatin, showing a dense condensate with sharp interfaces and few arrays in the dilute phase. Lines show means over five non-overlapping trajectory blocks; shaded bands show the standard deviation across block means. (b) Simulation snapshot and density profile of 30 bp chromatin, showing the lack of clear interfaces and reduced condensate density. Lines show means over five non-overlapping trajectory blocks; shaded bands show the standard deviation across block means. (c) Time progression of the largest connected component size in the coexistence simulations over the analysis period. The 25 bp simulation has a stable largest component containing 77–78/81 arrays over the analysis period. In contrast, for 30 bp the largest connected component is smaller on average (*≈*50/81) and fluctuates more widely (30–70/81). (d) Histograms showing the distribution of the number of unique arrays contacted by each tetranucleosome, from a total of 8,100 samples per linker (100 frames, 81 tetranucleosomes per frame). The 25 bp arrays are highly valent, with most arrays contacting 8–11 other partners. In contrast, the 30 bp arrays have significantly fewer interaction partners, with the majority contacting less than four other arrays. (e–f) Fluorescence microscopy images of 12-nucleosome arrays at 100 mM salt with 25 bp (e) and 30 bp linker DNA, stained with DAPI. Scale bar, 5 *µ*m.

To gain molecular and thermodynamic insight into these differences, we quantified the ‘largest connected component’ and ‘valency’ of chromatin arrays across simulations.

The largest connected component was defined as the largest set of chromatin arrays linked through pairwise contacts. Two arrays were considered connected if any pair of their nucleosomes lay within 13 nm. The valency of each chromatin array was quantified as the number of distinct arrays it contacted within this cutoff. Consistent with their phase behavior, 25 bp chromatin forms a stable, condensate-spanning percolated network encompassing nearly all arrays (77–78 out of 81), with individual arrays exhibiting high valency (frequently 8–11 partners per array). In contrast, the 30 bp chromatin displays low valency and transient inter-molecular connectivity, with most arrays interacting with fewer than four others and forming small clusters that continuously form and dissolve (Fig. 4c,d). Because direct-coexistence simulations exhibit sensitivity to finite-size effects, we repeated the simulations while systematically enlarging the cross-sectional dimensions of the simulation box. Supplementary Fig. S16 shows that both the density profiles and the resulting structural clustering remain qualitatively unchanged across these tests, indicating that our conclusions and qualitative comparisons are not dominated by finite size effects.

A key advantage of OpenCGChromatin is that it enables us to zoom in on chromatin within and outside condensates, connecting molecular structure with mesoscale phase behavior.

To probe how the conformational landscape of chromatin changes across the phase boundary, and how those changes, in turn, regulate condensation, we performed additional single-array HREX simulations of 25 and 30 bp tetranucleosome fibers (i.e., chromatin in infinite dilute conditions). We then analyzed over 30,000 tetranucleosome conformations from simulations probing both condensate-forming and dilute conditions (see Supplementary Information), representing the trajectories using symmetrized internucleosomal distance features [69]. In this representation, each tetranucleosome conformation is described by a set of six pairwise center-of-mass distances between nucleosomes (*d*_12_, *d*_13_, *d*_14_, *d*_23_, *d*_24_, *d*_34_). To account for the approximate symmetry of the tetranucleosome arrays (i.e., where nucleosomes 1 and 4 are equivalent at the edges of the fiber, and nucleosomes 2 and 3 are equivalent at the center), a symmetrized version of these features was used. For example, *d*_12_ and *d*_34_ are combined into *s*_1_ = *d*_12_ + *d*_34_ (see Supplementary Information). Although chromatin is not intrinsically symmetric along the 5’–3’ direction of the DNA, the coarse-grained representation is nearly symmetric with respect to fiber reversal, and the energetic differences between the two directions are negligible at the resolution considered here. Thus, symmetrization reduces noise and ensures that conformational clustering robustly captures structural variability. To characterize the conformational landscape, we applied *k*-means (*k* = 3) clustering to the combined ensemble, identifying three intuitive states: compact (C1, two nucleosome pairs in contact), semi-extended (C2, one pair in contact), and extended (C3, no nucleosome–nucleosome contacts) (Fig. 5a). Principal component analysis was then used to reduce dimensionality for visualization, with the first two components capturing over 77% of the overall variation (Fig. 5b).

**FIG. 5.**
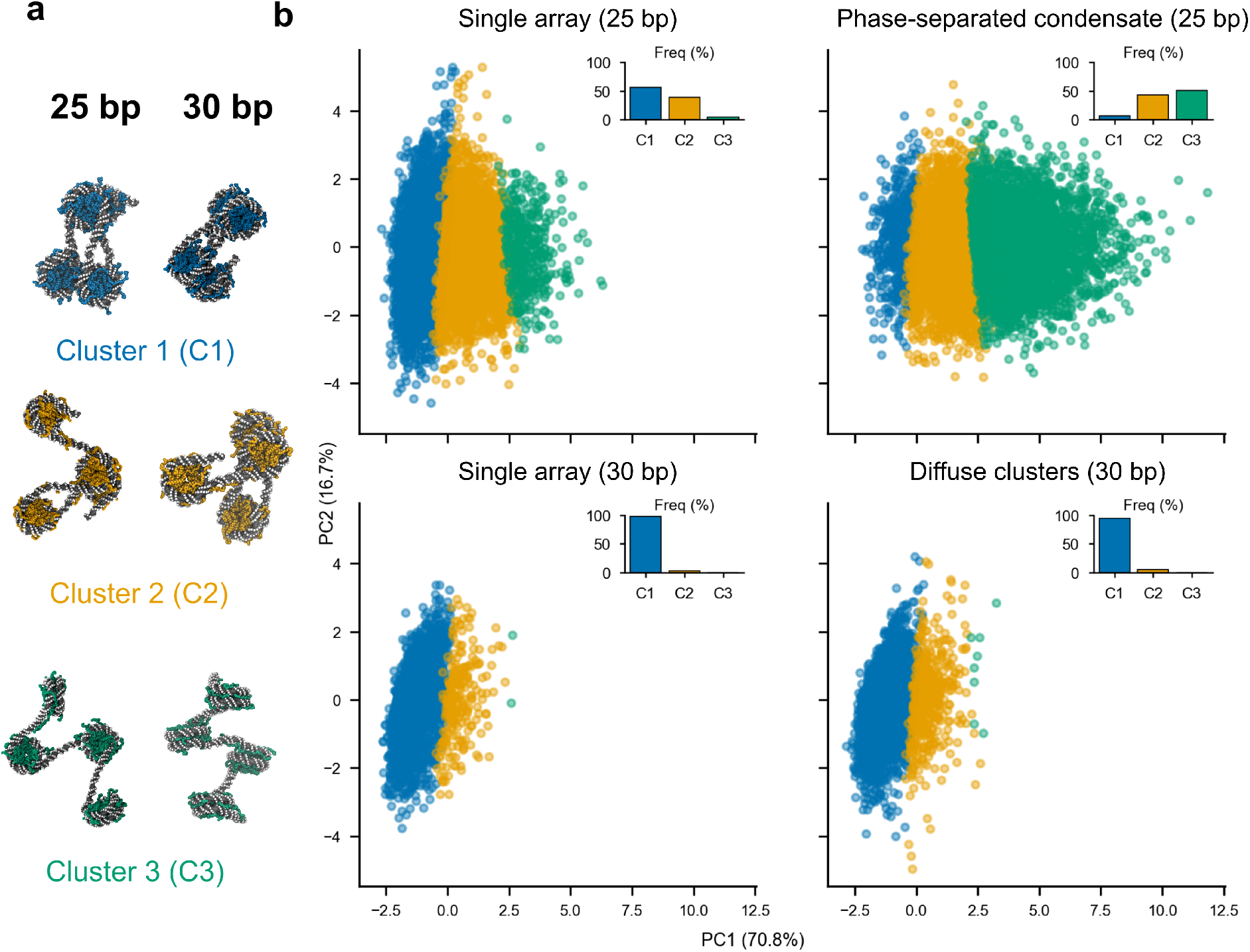
Structural features of chromatin phase separation. (a) Representative tetranucleosome configurations corresponding to each cluster, highlighting the intuitive interpretation of each cluster. Cluster 1 (C1) represents structures where two nucleosome–nucleosome contacts form, C2 represents structures with one nucleosome–nucleosome contact and C3 represents extended structures with no contacts. (b) Scatter plot showing configurations of tetranucleosome arrays in four conditions on the first two principal components, with inset bar charts showing the distribution of configurations between each cluster. The configurational distributions reveal that 25 bp chromatin has more heterogeneous structures than 30 bp chromatin in all conditions. The ensemble of 30 bp structures is similar in both the HREX and coexistence datasets, with both ensembles dominated by stacked C1 structures. In contrast, 25 bp chromatin undergoes extension in the coexistence dataset, with a clear shift in cluster distribution towards C2/C3.

Comparing structures across the phase boundary reveals that arrays with 25 bp and 30 bp linkers behave in fundamentally different ways. The 30 bp (10*N*) arrays remain conformationally homogeneous both in isolation and within condensates, dominated by compact zigzag conformations (C1) with narrow distributions along the first principal component (Fig. 5b). In contrast, the 25 bp (10*N*+5) arrays adopt more expanded conformations than 30 bp species in the dilute phase and undergo further expansion upon phase separation: their ensemble shifts from a mixture of compact and semi-extended states (C1 and C2) in dilute conditions to extended configurations (C3 *>* C2) inside condensates (Fig. 5b). This adaptive expansion and increased heterogeneity are consistent with a broader accessible conformational ensemble for 25 bp chromatin in the condensed phase, which may partially offset the translational entropy cost of condensation.

The expansion of 25 bp chromatin upon phase separation is physically consistent, as it increases tail accessibility and intermolecular valency, thus stabilizing the condensate. However, analysis of 25 bp chromatin structure via cryo-ET did not detect significant structural differences between arrays inside and outside condensates [9]. This may reflect the more limited sampling in the experiment versus the simulations, given the significant challenge of tracing individual fibres from the cryo-ET data, or that the fibres assigned to the dilute phase were captured close to the interface, where they may not yet be fully relaxed. In addition, selection bias toward well-resolved segments could under-represent the most expanded conformations in the experiment. Supporting this interpretation, the simulated tetramer *R*_*g*_ distributions inside and outside condensates show only a modest mean shift for 25 bp fibres (Fig. S15a,b), indicating that such subtle effects would be difficult to capture with a small set of experimental traces. A contribution from model limitations, such as modest deviations in DNA torsional or bending rigidity or overestimated DNA–DNA repulsion (as discussed earlier), could also explain the discrepancy, as these factors would favour more open states. We also verified that the clustering result is robust to the finite size of the direct-coexistence simulation box. Scanning the cross-section scale factor of the 25 bp system from 1.00× to 1.10× recovers the same three principal-component clusters with comparable relative populations (Fig. S16b).

As discussed earlier for individual chromatin fibers, the origin of the contrasting structural behaviour of 25 versus 30 bp linkers lies in the inter-nucleosome orientations imposed by the linker DNA. Linkers of 10*N*+5 bp introduce a half-helical turn rotational mismatch that frustrates regular face–face stacking between nucleosomes, preventing the formation of ordered zigzag fibers. This zigzag frustration yields structurally flexible, dynamic fibers that can open up in the dilute phase and reorganize within the condensate. Conversely, 10*N* linkers favor face–face stacking, which sequesters nucleosome surfaces and constrains conformational rearrangements. Linker-DNA-imposed zigzag frustration therefore acts to control phase behavior: 25 bp fibers can expand to maximise inter-molecular interactions, whereas 30 bp fibers remain locked in compact zigzag conformations.

The difference in intra-array nucleosome stacking for 25 versus 30 bp systems has important consequences for histone-tail accessibility and intermolecular connectivity. Energy-weighted contact analyses demonstrate that in 30 bp chromatin the H4 and H2A N-terminal tails—the strongest energetic contributors to nucleosome–nucleosome interactions—are buried in intra-array face–face interfaces, stabilizing folded fibers but suppressing inter-array binding (Fig. 6). In contrast, in the more expanded 25 bp fibers, all histone tails, including H4 and H2A N as well as H3 and H2A C, are exposed to the solvent and available to mediate a diverse set of inter-fiber interactions, increasing chromatin valency and the network connectivity of the condensate (Fig. 6). Thus, in 25 bp chromatin the interaction energy, that is concentrated in H4/H2A N intra-array contacts in 30 bp fibres, is redistributed across heterogeneous interactions involving all the histone tails.

**FIG. 6.**
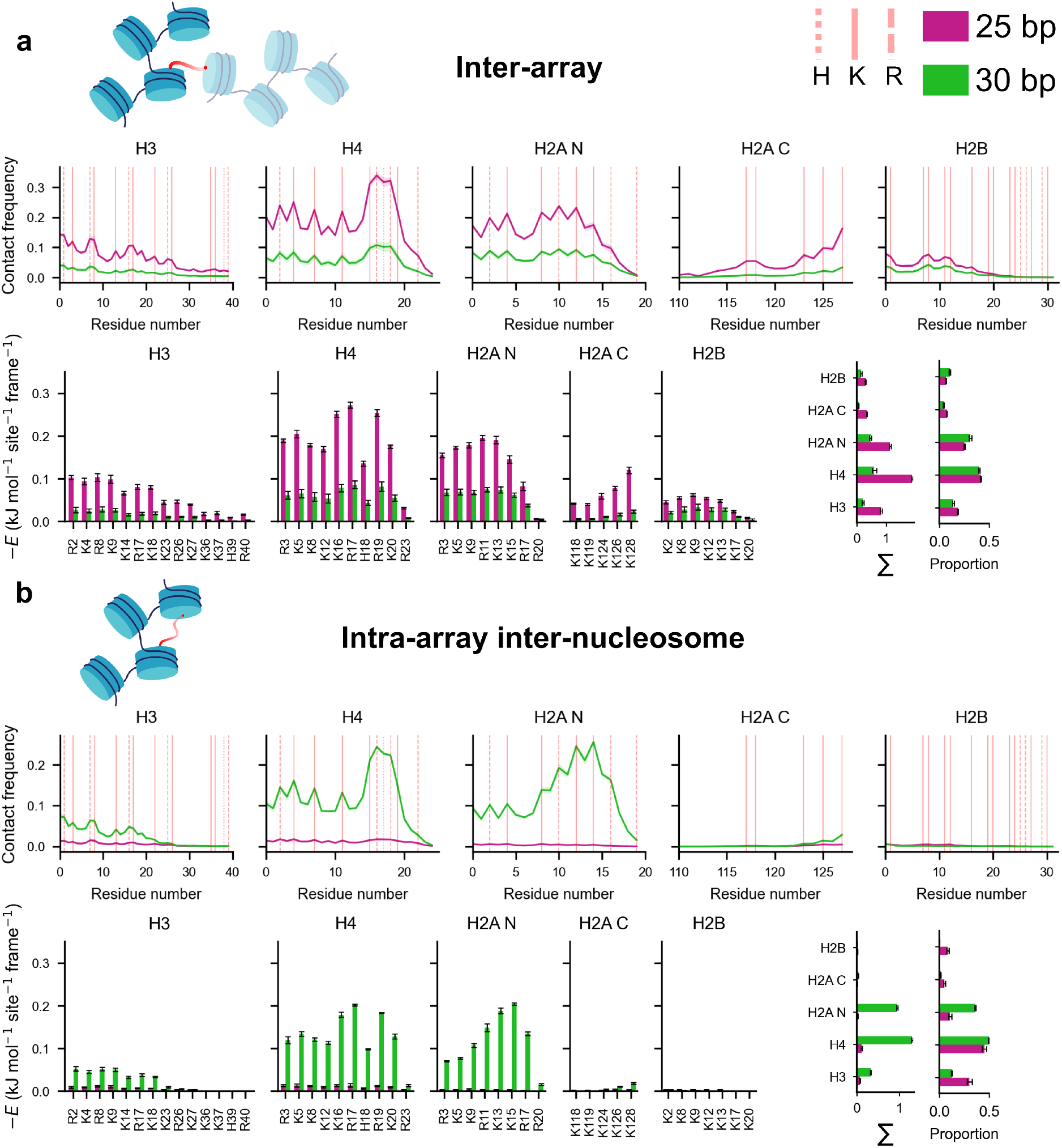
Residue-resolution contact analysis of condensed chromatin. For each histone tail (H3, H4, H2A N, H2A C, H2B), line plots show per-nucleosome, per-frame contact frequencies for tail residues, with vertical dashed lines marking cationic residues (H/K/R) along each tail. Bar plots beneath each line graph report the energy-weighted contact score for cationic residues in that tail and condition, highlighting residues that contribute the most overall interaction energy. Vertical bars on the right-hand side represent the total and proportional energetic contribution of each tail’s cationic residues. (a) Inter-array contacts (stabilize condensates, inter-molecular) [59]; (b) Intra-array inter-nucleosomal contacts (stabilize folded fibers, intra-molecular) are shown side-by-side for the two linker length conditions [59]. Plotted values are means computed over five non-overlapping trajectory blocks; error bars and shaded bands show the standard deviation across block means.

The differential roles of histone tails in 25 versus 30 bp condensates predicted from these unbiased OpenCGChromatin condensate simulations closely match our earlier results based on simulations also using OpenCGChromatin but in which nucleosome positions were restrained to those observed in the cryo-ET data [9]. That the same behavior emerges independently—now in simulations without any restraints and in the more demanding context of sampling equilibrium condensate densities—provides strong additional validation of the model.

Altogether, these results suggest that chromatin phase separation is governed not only by the balance of attractive sticker (histone tail-mediated) and repulsive spacer (DNA– DNA) interactions, but also by single-molecule conformational properties. Despite carrying a higher net negative charge—and hence greater electrostatic self-repulsion—the 30 bp arrays exhibit weak phase separation because they are locked in compact zigzag structures that prevent reorganization and multivalent engagement. In contrast, the simulations suggest that linker-induced frustration in the 25 bp arrays enables conformational expansion upon phase separation that exposes additional interaction sites, allowing fibers to reorganize and diversify their interaction modes. This structural adaptability increases both the number and variety of crosslinks that sustain the condensate, providing a direct coupling between single-fiber frustration, histone-tail accessibility, and network connectivity that defines how chromatin’s geometry and chemical makeup jointly control its phase behavior.

The distinct molecular interaction networks identified here—concentrated H4/H2A intra-array contacts in 30 bp versus diverse inter-array tail-mediated crosslinks in 25 bp condensates—suggest differences in macroscopic condensate material properties that are not directly accessible from our near-atomistic simulations. To probe these emergent physical properties, we carried out direct coexistence simulations of tetranucleosome chromatin arrays with 25 and 30 bp linker lengths using our minimal coarse-grained model for chromatin [25] (Fig. S14a,b), which approximates chromatin at lower resolution (1 bead per histone protein core and 1 ellipsoid per 5 base pairs) and enables extensive sampling for full binodals (Fig. S14c) and rheological calculations (Fig. S14d,e). Analysis of the shear stress relaxation modulus, *G* (*t*), reveals that behavior on short time scales (less than 10^−7^ s) is similar for both 25 and 30 bp chromatin, indicating that the fast, dynamic processes contributing to viscoelastic behavior on these timescales (primarily intra-molecular fluctuations such as bond and angle relaxations) are similar in both cases. In contrast, the longer time behaviour shows that *G* (*t*)_25 bp_ *> G* (*t*)_30 bp_. Since slower, collective mechanisms such as fiber rearrangement and diffusion dominate viscoelastic behavior in the longer time regime, we can infer that these processes are hindered in the more dense, strongly connected condensates formed by 25 bp chromatin (Fig. S14d). Furthermore, we find that the surface tension of 25 bp chromatin is higher than that of 30 bp chromatin, consistent with stronger intermolecular interactions (Fig. S14e). This concordance between the molecular interaction networks revealed by OpenCGChromatin and the macroscopic material properties calculated with the minimal model demonstrates that linker-DNA-controlled structural frustration has direct, measurable consequences for condensate rheology.

In summary, chromatin is an associative polymer whose structure and phase behavior depend not only on sticker-and-spacer composition but also on how spacers configure the stickers. In chromatin, linker-DNA length imposes mechanical constraints and nucleosome orientations that shift the fibre between two semi-flexible polymer regimes. Arrays with 10*N* linkers form rigid zigzag structures in which nucleosomes are locked into stable intra-array interactions, limiting their ability to form inter-array connectivity. In contrast, 10*N*+5 chromatin behaves as a more compliant semi-flexible polymer, where disrupted intra-array stacking restricts single-fibre compaction while enabling structural irregularity that favours multivalent, tail-mediated inter-array interactions. This structural adaptability enhances inter-array multivalency and explains the higher thermodynamic stability of 10*N*+5 chromatin condensates relative to 10*N* [8, 9, 26].

We note that the direct coexistence phase study employs short tetranucleosome arrays with uniform linker lengths. While the core physical mechanism—linker-DNA-imposed nucleosome orientations controlling tail accessibility and multivalent engagement—is expected to operate in longer fibers and heterogeneous linker sequences, the quantitative phase boundaries and material properties may differ for more complex chromatin substrates.

## D. Histone acetylation modulates structure and dynamics of long chromatin fibers

Finally, we applied OpenCGChromatin to investigate how histone acetylation modulates chromatin structure and dynamics on the scale of small genes (over 18 kb). We conducted 100 *µ*s simulations of 108-nucleosome fibers with regularly spaced 22 bp linkers and different distributions of partially acetylated nucleosomes. Because neither the stoichiometry nor the specific combinations of lysines acetylated per nucleosome within euchromatin are quantitatively resolved in vivo, we randomly acetylated 50% of lysine residues in the histone tails to explore the upper bound of plausible acetylation density. We compared three systems: (i) a 108-nucleosome control fiber in which all histones were unmodified (wild-type, 0 Ac); (ii) a partially acetylated fiber in which 12-nucleosome blocks of acetylated and unmodified nucleosomes alternated along the fiber (5 ×12 Ac), and (iii) a partially acetylated fiber in which the central 54-nucleosome region was acetylated, flanked by two unmodified regions of 27-nucleosomes each (54 Ac) (Fig. 7a–c). We find that acetylated systems consistently adopt more extended conformations with larger radius of gyration fluctuations, whilst the unmodified fiber collapses into a compact globule over around 40 *µ*s (Fig. 7d). This behaviour arises because the 108-nucleosome unmodified fiber constitutes a polymer with very high effective sticker (i.e. nucleosomes with histone tails) density. In the collapsed globule, nucleosomes can form many simultaneous tail-mediated contacts with other nucleosomes, thus maximising the overall valency of the system. Acetylated nucleosomes have fewer effective binding sites, reducing molecular valency and leading to the formation of elongated fiber-like structures in which they predominantly interact with their nearest neighbors.

**FIG. 7.**
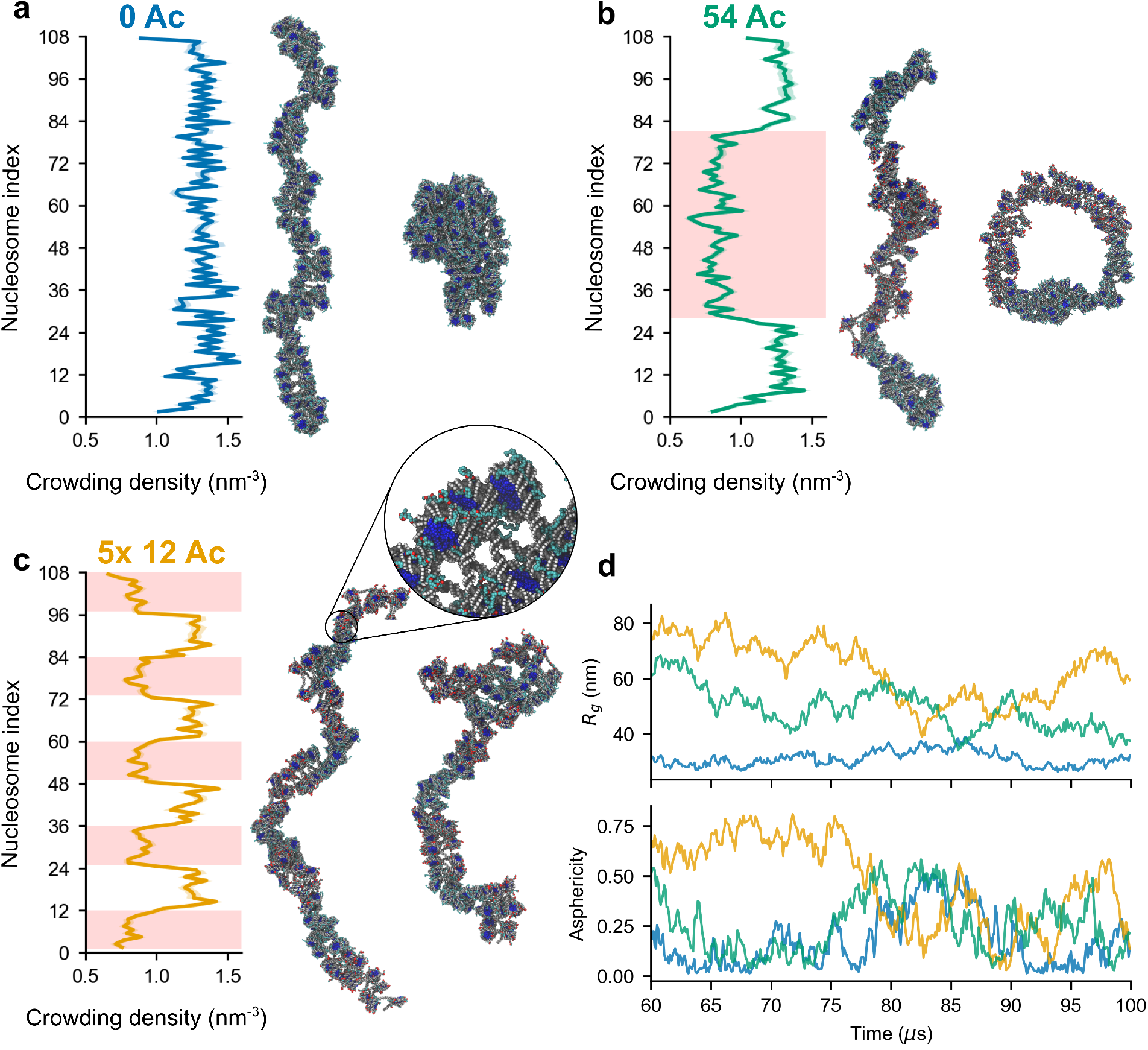
Acetylation promotes expanded structures and accessible DNA. (a–c) Crowding density profiles with acetylated region shaded in red and simulation snapshots at 10 (left) and *>*60 (right) *µ*s for 0 Ac (a), 54 Ac (b), and 5 × 12 Ac (c) fibers. Acetylated regions exhibit lower crowding densities, corresponding to more accessible DNA. Lines show means over five non-overlapping trajectory blocks; shaded bands show the standard deviation across block means. (d) Time progression of radius of gyration (*R*_*g*_) and asphericity for each fiber over the analysis window (60–100 *µ*s), where crowding density and contact calculations are performed.

Crowding density analysis reveals that linker regions flanked by acetylated nucleosomes have significantly lower local densities than those flanked by unmodified nucleosomes, corresponding to higher DNA accessibility (Fig. 7a– c). This increased accessibility provides a molecular mech-anism for the well-established role of histone acetylation in decompacting chromatin and promoting transcriptional activity [70, 71].

Analysis of nucleosome–nucleosome contact frequencies shows that acetylation produces a graded reduction in inter-nucleosome contacts (Fig. 8b). Because partially acetylated nucleosomes still form attractive interactions with other nucleosomes, but with reduced strength, their contacts break more readily and reform less frequently, leading to larger average inter-nucleosome separations between neighboring particles. However, by construction, the torsional and bending rigidity of the linker DNA constrain both the orientation and the spacing of consecutive nucleosomes, effectively limiting how far apart neighboring nucleosomes are relative to one another. This geometric coupling maintains frequent contacts between second-nearest neighbors (*j* = *i ±* 2), even though acetylation increases their mean separation. In contrast, for fourth neighbors (*j* = *i ±* 4), and even more so for longer-range pairs (|*i* − *j*| *>* 4), the cumulative increase in inter-nucleosome spacing over successive linkers cannot be compensated by DNA rigidity. Long-range contacts are then almost exclusively mediated by WT–WT pairs: the WT–WT contact fractions for |*i* − *j*| *>* 4 are similar in the 0 Ac and 54 Ac fibers, whereas the 5 × 12 Ac pattern shows a strong depletion of such bridges and very few Ac-containing long-range contacts. Thus, contiguous unacetylated stretches act as multivalent hubs that drive long-range compaction, while intervening acetylated segments behave as spacers that fragment these contact networks. Our results support the view that acetylation patterns are crucial determinants of higher-order chromatin folding, and the methodology we introduce offers a useful tool for probing exactly how such patterning differences drive fiber folding.

**FIG. 8.**
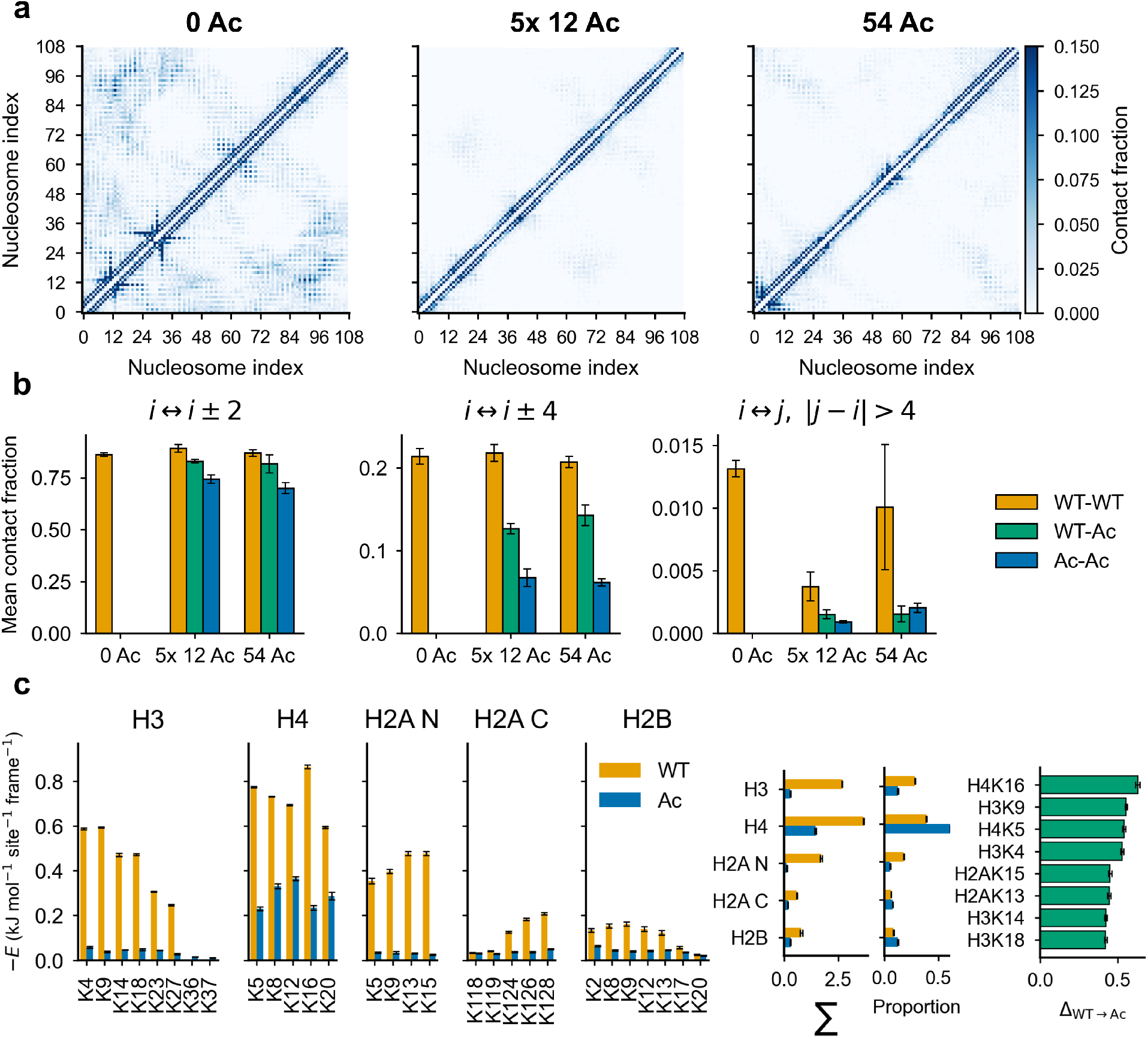
Nucleosome and residue-level contacts are modulated by acetylation. (a) Nucleosome–nucleosome contact frequency heatmaps for 0 Ac, 5 × 12 Ac and 54 Ac, normalized per frame. (b) Per-frame nucleosome–nucleosome contact frequencies separated by sequence distance and acetylation state. Acetylation produces a graded reduction in contacts: small for *i* ↔ *i* ± 2, moderate for *i* ↔ *i* ± 4, and large for longer-range pairs (|*i* − *j*| > 4). The frequency of long-range contacts scales with the length of uninterrupted unmodified stretches, indicating that extended unacetylated regions promote distant compaction. (c) Per-site, per-frame energy-weighted inter-nucleosomal lysine interaction-energy contributions, evaluated using a 1.6 nm Debye-range cutoff and plotted as *−E* so that larger positive values indicate stronger attractive contributions. In each frame in each trajectory, interactions involving each of 34 tail lysine sites (e.g., H3K9) are aggregated by lysine identity and acetylation state (WT/Ac), then weighted by their pairwise energy in the OpenCGChromatin Hamiltonian. Tail-level summary bars report the summed lysine contribution from each histone tail and the corresponding proportion of the total lysine-mediated signal for WT and acetylated states. The right panel highlights the 8 lysine sites exhibiting the largest drop in energy-weighted contribution upon acetylation. Bars show means over five non-overlapping trajectory blocks; error bars show the standard deviation across block means.

To identify which lysine sites are most affected by acetylation, we performed an energy-weighted residue-level contact analysis across all 108-nucleosome fibre simulations, using an extended Debye-range cutoff of 1.6 nm (2*λ*_*D*_). Unlike a binary contact analysis, which assigns a value of 1 when two beads are closer than a specified cutoff and 0 otherwise, this approach weights each lysine-containing interaction by its full instantaneous non-bonded pair energy at the observed bead–bead separation. We report this quantity as a positive stabilizing score, −*E* , such that larger values correspond to more favourable net interactions. This provides a site-resolved estimate of the stabilizing energetic contribution of lysine–DNA and lysine–protein associations to inter-nucleosome interactions.

For each lysine site (e.g., H4K16), we quantified its mean stabilizing interaction score and compared cases where the residue was acetylated with cases where it remained un-acetylated. Data from all three simulated systems (0 Ac, 5 × 12 Ac, and 54 Ac) were pooled and partitioned by lysine identity and acetylation state, allowing us to assess how acetylation reshapes the ensemble of site-specific interaction energies. To assess sensitivity to the interaction definition, we repeated the analysis using the shorter *σ*_*ij*_ +0.25 nm contact cutoff and compared both raw contact frequencies and energy-weighted scores; these controls are shown in Supplementary Fig. S17.

Our analyses highlight a subset of lysines whose acetylation most strongly weakens inter-nucleosome interactions (Fig. 8c). Remarkably, our simple coarse-grained model, using an unbiased approach of randomly acetylating 50% of tail lysines, successfully identifies H4K16 and H3K9—two of the best-characterized regulatory acetylation marks [9, 18, 64, 72, 73]—as having the largest energetic impact. This agreement suggests that much of these sites’ regulatory importance stems from basic physicochemical principles: acetylating a lysine within a charge block disrupts electrostatic interactions that cannot be compensated by non-electrostatic contacts, as neighboring cations still preferentially seek electrostatic partners. H4K16 (adjacent to R17) and H3K9 (adjacent to R8) are particularly sensitive because arginines cannot be acetylated, ensuring that acetylation of the neighboring lysine always creates incompatible interaction preferences within the charge block. Our analysis also reveals substantial reductions in contact energy upon acetylation of the closely spaced H2AK13 and H2AK15, which are ubiquitination targets involved in DNA repair [74] but not established transcriptional regulators. One possibility is that enzymatic acetylation of neighboring lysines may favor diacetylation over mono-acetylation, and while our random 50% acetylation frequently generates mono-acetylated states with conflicting interaction preferences, diacetylated tails may retain a greater capacity for cohesive tail–tail interactions. In contrast, lysines adjacent to non-acetylatable arginines consistently disrupt inter-nucleosome contacts regardless of local acetylation context, potentially explaining their more prominent regulatory roles in chromatin compaction and transcriptional control.

Aggregating the same 1.6 nm energy-weighted lysine contributions by histone tail provides a complementary view of this site-level analysis. In the wild-type state, H4 provides the largest absolute lysine-mediated inter-nucleosomal interaction energy, with H3 also making a substantial contribution. Acetylation strongly reduces the summed contributions from both tails, but the reduction is especially pronounced for H3, leaving the residual acetylated lysine-mediated signal disproportionately concentrated in H4. Thus, the proportional increase in H4 contribution after acetylation reflects redistribution within a depleted interaction network, not preservation or strengthening of H4-mediated contacts.

## III. DISCUSSION

This work presents OpenCGChromatin, a significant advance in chromatin simulation methodology that enables near-atomistic resolution investigations of chromatin structure, dynamics, and phase behavior at scales of hundreds of nucleosomes and tens of kb of DNA. The model expands the system sizes accessible to near-atomistic simulations by more than an order of magnitude compared to previous approaches [24, 25, 41], bridging the long-standing gap between molecular detail, chemical accuracy, and biologically relevant scales. This efficiency opens new avenues for predictive studies of chromatin regulation by post-translational modifications, chromatin remodeling, histone variant effects, and sequence-dependent DNA effects on chromatin-fibre structure. The present model treats solvent and ions implicitly via Debye–Hückel screening and does not include non-chromatin macromolecular crowders, chromatin-binding proteins (readers, writers, remodelers), or nuclear architectural constraints. Consequently, the results presented here isolate the intrinsic physicochemical behavior of chromatin and should be interpreted within this scope.

An important limitation of the Debye–Hückel framework is its inability to capture divalent-ion-specific effects. In physiological and experimental settings, Mg^2+^ ions can mediate ion bridging, correlation-driven attraction between like-charged surfaces, and specific binding to nucleosomal DNA [53]. These effects are not described by a mean-field monovalent screening model. The phase boundaries and material properties reported here should therefore be understood as conditional on an effective monovalent electrostatic environment. The experimental comparisons presented in this work employ reconstituted chromatin at defined monovalent salt concentrations (100–150 mM NaCl/KOAc).

OpenCGChromatin provides a predictive and mechanistically interpretable framework for exploring chromatin behavior, capable of both anticipating experimental trends and enriching their interpretation through molecular- and thermodynamic-level insight. Its predictive power is supported by the close agreement between the model results and available experimental observations. For instance, at the 12-nucleosome scale, the model captures the well-established oscillatory dependence of compaction on linker length, and explains that it arises from the mechanics of linker DNA and the B-DNA helical twist, which control the orientations of adjacent nucleosomes, the balance of face– face, face–side, and side–side intra-array contacts, and the contributions of the different histone tails. Linkers closer to 10*N* strongly involve the H4 basic patch, followed by H2AN, while linkers closer to 10*N*+5 are sustained by diverse tail-mediated interactions.

Histone acetylation modulates these interactions in a dosage- and pattern-dependent manner, weakening key electrostatic contacts, promoting dynamic, extended fibers, and modulating DNA accessibility. The model identifies H4K16 and H3K9 as sites whose acetylation most effectively disrupts chromatin compaction by neutralizing critical electro-static interactions. Remarkably, these two sites are known experimentally as sites of key epigenetic modifications, suggesting that biology has used these sites for regulation because of their energetic importance to chromatin structure.

Most importantly, OpenCGChromatin elucidates the molecular mechanism by which linker DNA length regulates chromatin phase separation, in close agreement with biochemical and cryo-ET observations [8, 9, 26]. Linker lengths close to integer helical repeats (10*N*) favor regular zigzag geometries stabilized by intra-array face–face stacking, consistent with cryo-EM, X-ray structures, and cryo-ET of 20–30 bp arrays [44, 45]. The compact zigzag sequesters key histone tails—particularly H4, which contributes most strongly to the inter-nucleosome interaction energy—into intra-array contacts, thereby limiting inter-array multivalent connectivity. In contrast, 10*N*+5 linker lengths frustrate regular zigzag folding by imposing a half-helical-turn rotational offset between nucleosomes. This frustration, also observed experimentally in 25 bp arrays inside and outside condensates [9], destabilizes intra-array stacking and drives molecular expansion within the condensed phase. Residue-level analysis reveals that the frustration of canonical stacking releases all histone tails for inter-array interactions, enabling the formation of a chemically diverse network of inter-array contacts that stabilizes the dense phase.

Our results position chromatin as a semi-flexible polymer whose thermodynamic behavior is governed by the degree of structural frustration encoded by its linker-DNA geometry. The stickers-and-spacers model [46] captures the nature of chromatin as an associative polymer, where nucleosome faces and histone tails can be classed as stickers and linker DNA as the spacers. In chromatin, however, the mechanical properties and geometry of these spacers, in addition to their solubility, have a profound impact on structure and phase separation. Linker-DNA spacers determine which stickers are sequestered or exposed, and therefore how multivalency is realised: 10*N* linkers stabilise compact, self-saturated fibres in which key H4 tails are buried in intra-array contacts, whereas 10*N*+5 linkers introduce structural frustration that liberates all histone tails and enable the formation of a dense, heterogeneous inter-array network that sustains condensate stability.

The coexistence of compact and frustrated chromatin states may represent a fundamental physical mechanism for balancing genome stability with regulatory plasticity. By tuning nucleosome spacing, cells can switch between structural regimes that either sequester or release histone tails, thereby suppressing or promoting multivalent interactions. This provides a geometric and thermodynamic mechanism for regulating interactions between chromatin fibers and, consequently, DNA accessibility. Subtle variations in linker length, post-translational modification, or chromatin-remodeler activity could thus serve as molecular toggles between compact, transcriptionally silent heterochromatin and frustrated, phase-separated euchromatin [8]. Compact fibers with 10*N* linkers provide rigid states, self-insulating structures that may help preserve epigenetic integrity and resist stochastic fluctuations. In contrast, frustrated 10*N*+5 condensate-forming arrays endow chromatin with adaptive flexibility, allowing reorganization in response to chemical and structural cues. Together, these complementary physical states highlight how energetic trade-offs between stability and adaptability shape chromatin organization and its capacity for regulation.

## IV. METHODS

### A. OpenCGChromatin

#### 1. Model overview

The chromatin model represents proteins with one bead per residue at C_*α*_ positions and DNA with one bead per nucleotide at C1’ positions. Protein–protein interactions employ Kim–Hummer Ashbaugh–Hatch parameters [75]. Histone cores are stabilized using elastic network models with the central 127 bp of nucleosomal DNA included to maintain stability whilst allowing terminal 10 bp to remain flexible for DNA “breathing”. DNA mechanics are captured through the CGeNArate model [47], employing sequence-dependent polynomial potentials. Virtual sites modify the DNA representation with phosphate (carrying −1*e* charge) and excluded volume beads (providing steric protection). Electrostatic interactions use Debye–Hückel screening. All simulations used the OpenCGChromatin package implemented in OpenMM [50, 51]. Detailed descriptions of the model Hamiltonian, virtual site implementation, force field parameterization strategy, and validation are provided in Supplementary Information.

#### 2. Simulation protocols

All simulations employed the LangevinMiddleIntegrator [76] with timestep 10 fs, damping coefficient 0.01 ps^−1^ and temperature 300 K.

##### Short fiber validation

Twelve-nucleosome arrays with linker lengths 18–63 bp were simulated using Hamiltonian replica exchange (HREX) implemented using the openmmtools package [77] with Debye length scaling (15 replicas, 0.8–1.3 nm, corresponding to approximately 150–60 mM salt). Nucleosomal DNA employed the 147 bp Widom-601 core sequence [78]. Simulations ran for 1.5 *µ*s, with the final 0.5 *µ*s used for analysis.

##### Phase separation

Tetranucleosome arrays with 25 bp and 30 bp linkers were first equilibrated using HREX (3 *µ*s). Direct coexistence simulations placed 81 arrays in an elongated periodic box (500 × 83 ×83 nm^3^), where the condensed phase spontaneously forms a slab-like domain spanning the short cross-section and producing two planar interfaces along the long axis under periodic boundary conditions. Simulations evolved for 20 *µ*s at *λ*_*D*_ = 1.0 nm (approximately 100 mM salt) with mass reduction factor 0.25. The final 10 *µ*s was used for analysis.

Full simulation protocols, including integrator parameters, enhanced sampling details, and system preparation workflows, are provided in Supplementary Information.

##### Long fiber dynamics

Three 108-nucleosome fibers with 22 bp linkers and different acetylation patterns (0 Ac,5 ×12 Ac, 54 Ac) were simulated for 100 *µ*s at *λ*_*D*_ = 0.8 nm (approximately 150 mM salt). In acetylated fibers, 50% of tail lysine residues per nucleosome were randomly modified. Mass reduction factor 0.5 was applied to accelerate sampling. The last 40 *µ*s of each simulation was used for contact and crowding density analysis.

Because coarse-graining and mass scaling alter the effective friction and inertia, all reported times correspond to simulation time and should not be directly mapped to experimental kinetics without independent calibration; however, equilibrium (time-averaged) properties, which are the focus of this work, are unaffected by mass scaling because particle masses cancel in the Boltzmann factor.

#### 3. Analysis methods

Sedimentation coefficients were calculated using the Kirkwood approximation applied to nucleosome centre-of-geometry positions (Supplementary Information Section 6.1). Radius of gyration (*R*_*g*_) was computed from nucleosome centre-of-geometry positions (Supplementary Information Section 6.3). Residue–residue contacts were defined using a distance cutoff of 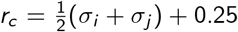 nm for each bead pair; non-parental or inter-nucleosomal contacts are those between two residues belonging to different nucleosomes, where the nucleosomal DNA and both 3^*′*^ and 5^*′*^ DNA linkers are considered part of each nucleosome. Energy-weighted residue-residue contacts weighted each contact by the pairwise interaction energy (Ashbaugh– Hatch plus Debye–Hückel) evaluated at the instantaneous separation using the OpenCGChromatin Hamiltonian. For the acetylated versus wild-type comparison of lysine sites in Fig. 8c, a longer contact cutoff of 1.6 nm (2*λ*_*D*_) was used before energy weighting in order to better capture the electrostatic energy beyond the close-contact shell. Nucleosome–nucleosome contacts used a 13 nm cutoff between centres of geometry and were classified by relative orientation into face–face, face–side, and side–side categories. Full mathematical definitions of all contact metrics, orientation-based classification, and the energy-weighting scheme are provided in Supplementary Information Section 6.2. Crowding density scores quantify the density of other protein/DNA beads around each linker DNA segment (Supplementary Information Section 6.4). Connected components were identified from nucleosome–nucleosome contact graphs (Supplementary Information Section 6.6). Structural clustering employed *k*-means (*k*=3) on symmetrized inter-nucleosomal distance features, with principal component analysis used to reduce dimensionality for visualization (Supplementary Information Section 6.5). Density profiles along the slab long axis were computed from binned particle coordinates averaged over the analysis window (Supplementary Information Section 6.7). Unless stated otherwise, reported means were calculated by dividing the relevant analysis window into five non-overlapping blocks; error bars and shaded bands denote the standard deviation across the five block means.

### B. Chromatin phase separation assay

Nucleosome arrays containing 12 repeats of the Widom 601 sequence with either 25 bp or 30 bp linker DNA were assembled as previously described [9, 26]. To induce phase separation, 2 *µ*M of nucleosome arrays were mixed with an equal volume of 2× phase separation buffer (40 mM Tris– OAc, pH 7.5, 200 mM KOAc, 10% glycerol, 1 *µ*M DAPI, and 0.2 mM EGTA). A total of 20 *µ*L of the mixture was transferred into a 384-well glass-bottom plate (Corning) and incubated for 30 minutes before imaging. The glass surface was passivated by siliconization (Sigmacote, Sigma-Aldrich), followed by incubation with 10 mg/mL bovine serum albumin (BSA) in double-distilled water for 10 min at room temperature. Fluorescence imaging was performed using a Leica SP8 confocal microscope equipped with a 405 nm laser for DAPI excitation.

### C. Minimal coarse-grained simulations

To complement the residue-resolution OpenCGChromatin simulations and probe macroscopic condensate properties such as surface tension and viscosity, we performed minimal coarse-grained simulations following the approach described previously [25]. In this minimal representation, each nucleosome is treated as a single particle with anisotropic interactions that capture the essential physics of face–face and face–side contacts (Fig. S14a). These simulations enable extensive sampling required for computing full phase diagrams (Fig. S14b,c) and rheological properties (Fig. S14d,e) that are not directly accessible from the higher-resolution simulations. Further details of the minimal model and the analysis methods relevant to this work are described in the Supplementary Information.

## V. DATA AVAILABILITY

Simulation data are available from the corresponding author upon reasonable request.

## VI. CODE AVAILABILITY

The OpenCGChromatin simulation package is available at https://github.com/CollepardoLab/OpenCGChromatin. Paper-specific simulation setup scripts and selected analysis workflows supporting the main quantitative results, including contact-analysis workflows, will be made available in a companion repository at https://github.com/CollepardoLab/Russell_2026 upon publication. Analysis scripts use standard Python packages including NumPy, SciPy, and MDAnalysis.

## SUPPLEMENTARY INFORMATION

Supplementary Information provides detailed descriptions of model architecture and Hamiltonian (Section 1, Figs. S1–S3), force field parameterization strategy (Section 2, Fig. S4), model implementation and protocol for building chromatin arrays (Section 3, Fig. S5), model validation (Section 4, Figs. S6–S7), simulation protocols including Hamiltonian replica exchange and direct coexistence setup (Section 5, Fig. S8), analysis methods for sedimentation coefficients, contacts, and structural clustering (Section 6), performance benchmarking specifications (Section 7), software implementation details (Section 8), and minimal model simulations and rheological analysis methods (Section 9, Fig. S14). Supplementary Figs. S9–S13 support the 12-nucleosome fiber results in main text Figs. 2 and 3, reporting per-frame nucleosome interaction counts, the new low-salt (5 mM) sedimentation runs, the cutoff-robustness of the nucleosome-level second-nearest-neighbour contact decomposition, the linker-length-independent intra-nucleosomal (parental) residue-level tail contacts, and the cutoff-robustness of the inter-nucleosomal residue-level histone-tail contact frequencies. Supplementary Figs. S15 and S16 support the direct-coexistence and tetramer clustering results in main text Figs. 4 and 5 by documenting the tetramer *R*_*g*_ distributions inside versus outside condensates and the cross-section-scaling robustness of both the slab density profile and the downstream structural clustering. Supplementary Fig. S17 reports close-contact energy-weighted lysine controls and raw close-contact and Debye-range contact frequencies supporting the Debye-range energy-weighted analysis in main text Fig. 8c. Supplementary Tables S1–S6 summarize force-field parameters, system details, performance and memory benchmarks, partial charges, validation sequences, and block-averaged direct-coexistence dense/dilute plateau densities and interface widths.

## AUTHOR CONTRIBUTIONS

Conceptualisation: KR, RCG; Model design: KR, YFC, JRE, DFG, MJM, JIPL, JH, MO, RCG; Data acquisition, analysis, or interpretation: KR, HZ, JRE, RCG; Performed simulations: KR, YFC, JRE; Model validation: KR, YFC, HZ; Creation of new software: KR, YFC, DFG; Visualization: KR, MJM; Writing original draft: KR, RCG; Editing: all authors; Supervision: RCG, MO, MR; Funding acquisition: RCG, MO, MR.

## ACKNOWLEDGMENTS

Research in the Collepardo-Guevara lab is supported by the UK Research Innovation (UKRI) Engineering and Physical Sciences Research Council (EPSRC) [EP/Z002028/1], following funding from the European Research Council (ERC) Consolidator Grant “ChromatinDroplets” under the European Union’s Horizon Europe research and innovation programme. We acknowledge EuroHPC Joint Undertaking for awarding access to MareNostrum5 at Barcelona Supercomputing Center (BSC), Spain [EHPC-REG-2025R01-166]. We also gratefully acknowledge the computer resources at MareNostrum and the technical support provided by BSC via the RES grant [RES-BCV-2025-1-0023]. The Orozco group is supported by the Center of Excellence for HPC H2020 European Commission; BioExcel-3: Centre of Excellence for Computational Biomolecular Research [European Union: 101093290; Ministerio de Ciencia e Innovación: PCI2022-134976-2]; the MDDB project [101094561]; Spanish Ministry of Science [PDI2024-155247NB-100]; and European Regional Development Fund, ERFD Operative Programme for Catalunya, the Catalan Government AGAUR [SGR2021 00863]. The IRB Barcelona is the recipient of a Severo Ochoa Award of Excellence from MINECO. Modesto Orozco is an ICREA Academia Fellow. Research in the Rosen lab was supported by the Howard Hughes Medical Institute, a Paul G. Allen Frontiers Distinguished Investigator Award, grants from the Welch Foundation [I-1544] and the National Institutes of Health [R35GM141736]. We thank Daniel and David Isenberg for their generous support of the MBL Chromatin Collaborative in honor of their father, Irvin Isenberg.

