## Supplementary Information for "Near-atomistic simulations reveal the molecular principles that control chromatin structure and phase separation"

(Dated: May 18, 2026)

#### SUPPLEMENTARY METHODS

This document provides supporting information for the manuscript "Near-atomistic simulations reveal the molecular principles that control chromatin structure and phase separation".

Section 1 describes the model architecture and Hamiltonian including bead mappings, bonded and non-bonded interactions, and virtual site implementations. Section 2 details force field parametrization utilizing experimental sedimentation data and comparison to previous models. Section 3 covers system building protocols for chromatin arrays. Section 4 presents model validation including DNA mechanics and cryo-ET fiber compaction analysis. Section 5 describes simulation protocols including Hamiltonian replica exchange and direct coexistence methods. Section 6 details analysis methods for structural and energetic characterization. Section 7 describes performance benchmarking on CPU and GPU hardware. Section 8 lists software and computational resources. Section 9 describes our lab's minimal model for chromatin and the rheological analyses deployed in this work.

Supplementary Figs. S1–S17 are at the end of this document. Supplementary Tables S1–S6 summarize key parameters, system details, validation datasets, and block-averaged numerical values for the direct-coexistence analysis.

#### I. MODEL ARCHITECTURE AND HAMILTONIAN

##### A. Bead mapping and structural representation

The OpenCGChromatin model represents proteins with one bead per residue positioned at the $C_\alpha$ atom. Histone core residues are stabilized using an elastic network model (ENM), while tail residues remain fully flexible. The designation of core versus tail residues follows previous work [1], originally determined from bias-exchange metadynamics simulations using an all-atom forcefield [2].

DNA is represented using one bead per nucleotide at the C1' position, following the CGeNArate model [3].

To ensure physically realistic protein–DNA interactions (i.e., preventing histone tails entering the center of the DNA helix), we modified this representation with two types of virtual sites:

- **Phosphate sites**: Positioned geometrically to match crystallographic phosphate locations [4], carrying DNA negative charges
- **Excluded volume sites**: Placed in the DNA major groove to provide steric protection, preventing unphysical histone tail penetration into the helix

* Corresponding authors

Fig. S1 illustrates the complete coarse-grained representations of the nucleosome and a chromatin fiber, including a cut-away view showing bead and bond representations for DNA and histone core. Fig. S2a schematically illustrates the virtual site definitions and placements.

#### B. Model Hamiltonian

The total potential energy is given by:

$$U_{\text{tot}} = U_{\text{prot-bonded}} + U_{\text{DNA-bonded}} + U_{\text{non}-\text{ele}} + U_{\text{ele}} \tag{1}$$

where $U_{\text{prot-bonded}}$ is the potential of protein–protein/protein–DNA bonded terms, $U_{\text{DNA-bonded}}$ is the sum of all DNA–DNA bonded terms, $U_{\text{non}-\text{ele}}$ is the sum of all non-electrostatic non-bonded interactions, and $U_{\text{ele}}$ is the sum of all electrostatic non-bonded interactions between charged atom pairs.

*a. Protein bonded terms* All protein bonded interactions use harmonic potentials:

$$U_{\text{harm}}(r) = k_{\text{harm}}(r - r_0)^2 \tag{2}$$

with $k_{\text{harm}} = 8031$ kJ mol$^{-1}$ nm$^{-2}$ for backbone bonds.

Histone tails are modelled as fully flexible chains with bonds of equilibrium length $r_0 = 0.381\,\text{nm}$ connecting consecutive residues. No angle or dihedral terms are included for tail residues, following our previous work [1].

The histone core structure is maintained via an elastic network model constructed by adding a harmonic bond between any pair of residues with an initial separation of less than $0.75\,\text{nm}$. The equilibrium length $r_0$ for each ENM bond is set to the $C_\alpha$–$C_\alpha$ separation calculated from the atomistic PDB structure [4]. The force constant is set to $k_{\text{ENM}} = 8031$ kJ mol$^{-1}$ nm$^{-2}$.

To maintain the positional stability of individual nucleosomes while retaining limited flexibility at the DNA entry and exit sites, we included the phosphate beads of the central 127 bp of nucleosomal DNA, centred on the dyad axis, in the core elastic network model (ENM).

This choice is motivated by the structural organisation of the nucleosome core particle: in the crystal structure of Luger et al., the 10-bp segment at each DNA terminus is described as essentially straight, while the remaining nucleosomal DNA follows the wrapped superhelical path around the histone octamer [5]. The flexible terminal segments correspond to the entry/exit regions, where nucleosomal DNA accessibility and spontaneous site exposure are most pronounced [6]. We emphasise that this treatment is a pragmatic stabilisation scheme rather than a complete model of nucleosome unwrapping: it preserves the structural register of the nucleosome core particle and prevents passive repositioning or disassembly during chromatin-fibre simulations, while still allowing local "breathing" motions at the entry and exit sites.

*b. DNA bonded terms* DNA bonds and angles employ sequence-dependent polynomial potentials from CGeNArate [3]. For bonds:

$$U_{\text{DNA-bond}}(r) = \sum_{n=2}^{4} k_n (r - r_0)^n \tag{3}$$

For angles:

$$U_{\text{DNA-angle}}(\theta) = \sum_{n=2}^{4} k_n (\theta - \theta_0)^n \tag{4}$$

Parameters ($k_n$, $r_0$, $\theta_0$) are sequence-dependent and loaded from CGeNArate data tables. These potentials are applied to overlapping sets of base pair tetramers, where each tetramer comprises eight C1′ beads. Within each tetramer, 16 unique bonds and four unique angles are defined, each with distinct equilibrium values and force constants for every possible base pair tetramer sequence. Fig. S2b illustrates the pattern of sequence-dependent bonds and angles present in each base-pair tetramer. In addition to these, sequence-independent long-range bonds act between cross-strand 4th and 5th nearest neighbors (i.e., w0–c5/c6 and c0–w5/w6 using the notation of Fig. S2b).

To ensure numerical stability, we add a weak repulsive force between consecutive excluded volume virtual sites:

$$U_{\text{rep}}(r) = \frac{k_{\text{rep}}}{r} \tag{5}$$

with repulsion constant $k_{\text{rep}} = 0.025\,\text{kJ}\,\text{nm}\,\text{mol}^{-1}$. This addition prevents simulation instabilities caused by overlap of two sites, while having negligible effect on DNA mechanical properties (Fig. S6a).

*c. Non-electrostatic interactions* Non-electrostatic interactions use the Ashbaugh–Hatch potential:

$$U_{\mathrm{AH}}(r_{ij}) = \begin{cases} 4\epsilon_{ij}\lambda_{ij}\left[\left(\frac{\sigma_{ij}}{r_{ij}}\right)^{12} - \left(\frac{\sigma_{ij}}{r_{ij}}\right)^{6} - \delta\right], & r_{ij} < 2^{1/6}\sigma_{ij} \\ 4\epsilon_{ij}\left[\left(\frac{\sigma_{ij}}{r_{ij}}\right)^{12} - \left(\frac{\sigma_{ij}}{r_{ij}}\right)^{6} - \lambda_{ij}\delta\right] + \epsilon_{ij}(1-\lambda_{ij}), & r_{ij} \geq 2^{1/6}\sigma_{ij} \end{cases} \tag{6}$$

where $\lambda_{ij} \in [0, 1]$ controls hydrophobicity (0 = fully hydrophobic, 1 = fully hydrophilic), $\epsilon_{ij}$ sets the energy scale, $\sigma_{ij}$ is the particle diameter, and the potential is shifted by $\delta = (\sigma_{ij}/r_c)^{12} - (\sigma_{ij}/r_c)^{6}$ and truncated at cutoff distance $r_c = 3\sigma_{ij}$.

The Ashbaugh–Hatch form was chosen over a conventional Lennard–Jones potential because its harder repulsive core provides a more effective steric barrier, helping prevent unphysical interpenetration when the attractive well is shallow.

*d. Electrostatic interactions* Electrostatic interactions are screened using the Debye–Hückel potential:

$$U_{\mathrm{ele}}(r_{ij}) = \frac{q_i q_j}{4\pi\epsilon_0\epsilon_r r_{ij}} \exp\left(-\frac{r_{ij}}{\lambda_D}\right) \tag{7}$$

where $q_i$ and $q_j$ are the charges of the interacting beads, $\epsilon_0$ is the vacuum permittivity, $\epsilon_r$ is the relative permittivity of the solvent (set to 80), and $\lambda_D$ is the Debye screening length.

The Debye length is related to ionic strength by:

$$\lambda_D = \sqrt{\frac{\varepsilon_r\varepsilon_0 k_B T}{2N_A e^2 I}} \tag{8}$$

where $N_A$ is Avogadro's constant, $e$ is the fundamental charge, $k_B$ is Boltzmann's constant, $T$ is temperature, and $I$ is the ionic strength.

Typical values of $\lambda_D$ used in the model range from 0.8–2.0 nm, corresponding to implicit salt concentrations of approximately 150–25 mM. Behaviour at 150 mM ($\lambda_D \approx 0.8$ nm) is a particularly important target for the model as this corresponds to physiological concentration, and the model's agreement with experimental compaction trends decreases at extremely low salt concentrations (Fig. S10).

#### C. Virtual site implementation

Virtual site positions are defined using OpenMM's `LocalCoordinatesSite` class, which constructs a local, orthogonal coordinate system from three reference C1' beads and places the virtual site at a fixed position within that frame.

**Excluded volume sites:** The coordinate system is defined by a C1' bead on each strand of a base pair (w0, c0) and the next C1' bead on the w-strand (w1). The site is placed at (0.0, 0.0, -0.3) in the local frame, positioning it near the base pair's midpoint to provide steric bulk in the major groove.

**Phosphate sites:** The coordinate system is defined by three consecutive C1' beads along one strand. The site is placed at (0.15, -0.30, 0.13) in this frame, a position chosen to match the crystallographic location of the phosphate group in PDB 1KX5 [4].

These precise definitions ensure that the virtual sites are positioned in a chemically meaningful way, allowing the DNA model to present a physically realistic steric and electrostatic profile to the protein components (Fig. S2a).

#### D. Force field parameters

The model inherits several parameter sets from established models. Protein–protein interactions use the Kim–Hummer Ashbaugh–Hatch parameters [7], while DNA bonded terms employ CGeNArate sequence-dependent polynomials [3]. Partial charges are assigned following our previous residue-resolution chromatin model [1].

Acetylated lysine parameters involve neutralisation of charge ($+1e \rightarrow 0$) and increased $\sigma$ to account for acetyl group volume.

Following previous work [1], DNA–DNA Ashbaugh–Hatch interactions are set to zero. DNA–DNA interactions are mediated only by electrostatic repulsion between phosphate groups, which is sufficient to prevent overlap over the range of $\lambda_D$ values used.

#### E. Bead charges

Charges assigned to each bead type are listed in Table S4.

##### F. Non-bonded exclusions

To prevent double-counting of interactions already captured in bonded terms and to improve computational efficiency, we apply the following non-bonded exclusions. For protein, non-bonded interactions between directly bonded bead pairs are excluded, while elastic network bonds do not create exclusions. For DNA phosphate virtual sites, we exclude $\pm 2$ phosphates on the same strand and $\pm 5$ base pairs cross-strand. This rationale avoids double-counting phosphate–phosphate repulsion already present in CGeNArate bonded potentials while minimising the total number of exclusions to improve computational performance.

#### II. FORCE FIELD PARAMETRIZATION

##### A. Comparison to previous LAMMPS model

Our previous chemically specific chromatin model [1], implemented in LAMMPS, used stronger non-bonded protein–DNA interactions than the present OpenCGChromatin model. These interactions were sufficient to stabilize individual nucleosomes in simulations where nucleosomal DNA was allowed to move relative to the histone core, and enabled studies of nucleosome plasticity and DNA unwrapping. However, in chromatin-fiber simulations at physiological salt, this fully flexible treatment did not robustly preserve nucleosome registration, leading to passive nucleosome repositioning on simulation timescales. This rapid loss of nucleosome register is inconsistent with reconstituted chromatin arrays, where linker-length-dependent structural and phase-separation properties persist *in vitro*, implying that nucleosome positions remain well defined on timescales far exceeding the microsecond–millisecond regime accessible to simulations [8, 9].

The previous model could alternatively be run with all 147 bp of nucleosomal DNA attached to the histone core, thereby preserving nucleosome registration during fiber simulations. However, the resulting high-resolution fiber model did not reproduce the experimentally observed linker-length dependence of chromatin compaction. In particular, it did not capture the oscillatory dependence of sedimentation coefficient on linker length at 150 mM salt, including the characteristic difference between $10N$ and $10N{+}5$ bp linker lengths (Fig. S3) [10, 11].

It is important to distinguish this chemically specific LAMMPS model from the minimal chromatin model introduced in the same study [1]. In the minimal model, nucleosomal DNA was permanently attached to the histone core in both the "breathing" and "non-breathing" variants, with the two modes differing only in the amount of DNA attached to the core [1]. Because nucleosome registration was therefore enforced by construction, the minimal model avoided passive repositioning during fiber simulations and was better able to reproduce the oscillatory dependence of fiber compaction on linker length [8]. However, this was achieved using a lower-resolution, more coarse-grained representation of the nucleosome than the chemically specific model considered here.

Together, these results highlight a key trade-off in coarse-grained chromatin modeling between nucleosome-level structural detail, enforced nucleosome registration, and fiber-level compaction behavior. Parameterizations with sufficiently strong non-bonded protein–DNA interactions to stabilize individual nucleosomes may over-stabilize histone–DNA contacts relative to the DNA mechanical penalties that shape fiber-level structure. Conversely, weakening these interactions can improve fiber-level compaction trends, but leaves individual nucleosomes insufficiently stable unless additional constraints are introduced to preserve their register.

The present OpenCGChromatin model addresses this trade-off using a different stabilization strategy. Rather than relying primarily on strong non-bonded protein–DNA interactions to maintain nucleosome integrity, we reduce the non-specific protein–DNA interaction strength and preserve nucleosome registration through the core ENM. As described above, only the central 127 bp of nucleosomal DNA are included in the core ENM, while the terminal 10 bp on each flank remain flexible. This provides a pragmatic compromise: the central wrapped register of the nucleosome is maintained during chromatin fiber simulations, while limited flexibility is retained at the entry and exit sites.

##### B. Protein–DNA interaction parameterization

We optimized protein–DNA non-bonded interactions to reproduce experimental trends in compaction with respect to linker length.

*a. Design principles* Following previous work [1], we use a simplified scheme in which protein–phosphate interactions provide the dominant non-electrostatic attraction, with the $\lambda$ value for protein–phosphate set $10\times$ larger than for protein–C1' or protein–excluded volume interactions.

*b. Parameter optimization strategy* We employed a two-stage optimization:

**Stage 1—Steric parameters:** We established $\sigma$ values for C1' and excluded volume beads through iterative testing, with the goals of preventing histone tail penetration through the DNA helix and eliminating numerical instabilities. Selected values are $\sigma = 0.575$ nm for the C1' bead and $\sigma = 0.900$ nm for the excluded volume bead (which provides major groove protection).

**Stage 2—Energetic parameters:** With steric parameters fixed, we optimized protein–phosphate $\sigma$, protein–phosphate $\lambda$ (relative to other DNA beads, maintaining the 10:1 ratio), and electrostatic cutoff $r_c$. This optimization aimed to reproduce experimental sedimentation coefficient trends [10, 11] across multiple linker lengths.

The first stage was a broad grid search using $2\,\mu s$ regular MD simulations of 12-nucleosome arrays with 25 and 30 bp linkers at two $\lambda_D$ values—$0.8\,\mathrm{nm}$ (150 mM) and $1.3\,\mathrm{nm}$ (60 mM). This tested all combinations of three phosphate $\sigma$ values (0.40, 0.45, and 0.50 nm), two $\lambda$ values (0.10 and 0.025), and three electrostatic cutoffs (3.5, 4.0, and $4.5\times\lambda_D$). Sedimentation coefficients were compared against experimental values to identify promising candidates (Fig. S4a).

For selected candidates, $1.5\,\mu s$ Debye-length Hamiltonian replica exchange (HREX) simulations were performed for 25, 30, and 58 bp linkers (Fig. S4b). The final parameters were then selected based off these HREX simulations.

*c. Final parameters* Optimized protein–DNA Ashbaugh–Hatch parameters are summarized in Table S1. Energy scale: $\epsilon = 4.184$ kJ mol$^{-1}$ for all protein–DNA interactions. Electrostatic cutoff: $r_c = 4.5\lambda_D$, where $\lambda_D$ is the Debye length of the simulation.

The final OpenCGChromatin parameterization combines weakened generic protein–DNA Ashbaugh–Hatch attractions with a longer Debye-length-scaled electrostatic cutoff. Nucleosome registration is maintained through the core ENM, which includes the central 127 bp of nucleosomal DNA while leaving the terminal 10 bp on each flank flexible, as described above. This combination preserves stable nucleosome core particles during chromatin-fiber simulations while allowing the protein–DNA interaction strength to be tuned for fiber-level behavior.

With these parameters, the model reproduces the experimentally observed linker-length dependence of chromatin compaction, including the oscillatory sedimentation-coefficient trends discussed in the main text (Fig. 2).

#### III. MODEL IMPLEMENTATION AND SYSTEM BUILDING

##### A. Software implementation

The model is implemented as a Python package built on OpenMM's Python application programming interface (API). The package, called OpenCGChromatin, integrates directly with the OpenMM Python API, allowing seamless integration with Python-based enhanced sampling and analysis schemes.

The `system_building` module prepares OpenMM `System` objects by iterating over the `Topology` and adding all Hamiltonian terms using OpenMM force objects. Harmonic bonds are implemented using the built-in `HarmonicBondForce` class, while all other force field terms are implemented as various `CustomForce` objects.

The model loads Ashbaugh–Hatch parameters from parameter files, with a mapping dictionary assigning masses, indices, and charges to particles. Equilibrium distances and force constants for DNA bonds and angles are derived from tables using canonical tetramer sequence indices, loaded from data files provided with the CGeNArate model [3].

##### B. Chromatin array construction

Chromatin fibers are constructed using the `NucleosomeArray.build_array()` method, which loads nucleosome templates, calculates necessary isometries for linker DNA segments, builds linkers, and stitches together the final array structure.

**Nucleosome templates** Three nucleosome templates with different entry/exit DNA angles are available. The **atomistic template**, derived directly from PDB 1KX5 [4], has a closed entry/exit angle (∼40°) that produces steric clashes when building arrays, making it suitable only for mononucleosome simulations. The closed coarse-grained template is generated by applying a harmonic pulling force to the terminal DNA beads of a mononucleosome while keeping the central 127 bp bonded to the histone core via the elastic network, producing an entry/exit angle of ∼90° suitable for most linker lengths. The open coarse-grained template employs a stronger pulling protocol to yield an entry/exit angle of ∼120°, which is required for linker lengths close to $10N$+5 (e.g., 25 bp, 36 bp) to avoid steric clashes when consecutive nucleosomes are positioned out-of-phase.

**Geometric construction** The geometry of the fiber depends on the entry/exit angle of the nucleosome template, the helical twist inherent in B-form DNA (∼10.5 bp per turn, or ∼36° per base pair), and the linker DNA length.

For a given linker length, the position and orientation of the next nucleosome is determined by constructing linker DNA with appropriate length and sequence, calculating the rigid body transformation (rotation + translation) that connects the exit of nucleosome $i$ to the entry of nucleosome $i+1$, applying this transformation to position the next nucleosome template, and bonding the linker DNA to both nucleosomes.

The DNA twist means that linker lengths near $10N$ bp (∼360° total twist) position consecutive nucleosomes in-phase for face–face stacking, while $10N$+5 bp (∼540° twist) rotates them out-of-phase by ∼180°.

Fig. S5 shows a schematic illustrating how initial configurations of chromatin arrays are constructed, including the three nucleosome templates and their compatibility with different linker lengths.

All fibers simulated in this work were built using the coarse-grained templates. All fibers with a linker length of 25 or 36 bp were built using the "open" template, all other fibers used the "closed" template (Fig. S5).

**Post-translational modifications** Histone acetylation is implemented via the `NucleosomeArray.acetylate_random()` method. For each nucleosome designated as acetylated, 50% of tail lysine residues are randomly selected and modified. Acetylation neutralises the positive charge ($q = +1e \to 0$) and increases $\sigma$ to account for acetyl group volume.

#### IV. MODEL VALIDATION

##### A. DNA model validation

To verify that our modifications to CGeNArate (virtual sites, non-bonded interactions, and $U_{\text{rep}}$) preserve accurate DNA mechanics, we simulated the 13 sequences of the miniABC dataset [12] and compared end-to-end distance distributions between four models: the original CGeNArate model [3], all-atom miniABC simulations [12], the OpenCGChromatin DNA model with $U_{\text{rep}}$, and the OpenCGChromatin DNA model without $U_{\text{rep}}$. The end-to-end distance is calculated between the first and last C1' atoms of the 3'→5' strand. The miniABC dataset contains 13 sequences with 18 nucleotides each, covering all possible base pair tetramer combinations (Table S5).

Fig. S6 shows the comparison of end-to-end distance distributions for the four models. The OpenCGChromatin DNA model closely matches the original CGeNArate model and all-atom reference data, confirming that our modifications preserve sequence-dependent mechanical properties while enabling stable protein–DNA interactions. The addition of $U_{\text{rep}}$ has negligible impact on this metric, justifying the functional form adopted.

##### B. Cryo-ET fiber validation

To further validate the model's ability to reproduce chromatin fiber compaction, we first performed fragment-wise radius of gyration ($R_g$) analysis comparing simulated 25 bp and 30 bp chromatin arrays against cryo-electron tomography (cryo-ET) data [9] (Fig. S7a). For each fragment length $k \in \{7, \ldots, 12\}$ nucleosomes, we calculated $R_g$ from contiguous $k$-nucleosome windows within our simulated 12-nucleosome arrays. Histograms show the distributions of $R_g$ values for 25 bp (magenta) and 30 bp (green) linker arrays, with vertical lines marking individual cryo-ET fragments [9]. Across all fragment lengths, 25 bp chromatin consistently exhibits larger $R_g$ values than 30 bp chromatin, in quantitative agreement with experimental observations. Importantly, the cryo-ET fragment $R_g$ values fall within or close to the simulated distribution ranges, validating the model's accuracy in reproducing not only local DNA mechanics but also chromatin compaction and the linker-length-dependent structural differences observed experimentally.

To further compare our simulated structures with the cryo-ET data, we compared three structural metrics calculated from consecutive triplets of nucleosomes; $D$, the distance between second-nearest-neighbor nucleosomes; $\alpha$, the dihedral angle between the planes of nearest-neighbor nucleosomes; and para, the dihedral angle between the planes of second-nearest-neighbor nucleosomes. The distance $D$ was calculated as the Euclidean distance between nucleosome centers of mass. The orientation of each nucleosome was defined by a unit vector $\mathbf{z}$ perpendicular to its disc-shaped plane, calculated from specific atom pairs (atoms 465 and 952) within each nucleosome chain:

$$\mathbf{z} = \frac{\mathbf{r}_{952} - \mathbf{r}_{465}}{\|\mathbf{r}_{952} - \mathbf{r}_{465}\|} \tag{9}$$

Following reference [9], dihedral angles were obtained from dot products of these orientation vectors. For 25 bp linker arrays (2.5 turns of DNA between nucleosomes), adjacent nucleosomes adopt opposite orientations, so one vector was flipped and the resulting angle was mapped to the range 180°–270°, where 180°corresponds to parallel planes and 270° to perpendicular planes:

$$\alpha_{25\text{bp}} = 180^\circ + \frac{1}{2}\arccos(\mathbf{z}_{\text{N}} \cdot (-\mathbf{z}_{\text{N+1}})) \times \frac{180^\circ}{\pi}. \tag{10}$$

For 30 bp linker arrays (3 DNA turns), adjacent nucleosomes are aligned and the angle was restricted to 0°–90°:

$$\alpha_{30\text{bp}} = \min(\theta, 180^\circ - \theta), \quad \text{where } \theta = \arccos(\mathbf{z}_{\text{N}} \cdot \mathbf{z}_{\text{N+1}}) \times \frac{180^\circ}{\pi}. \tag{11}$$

The alternating (para) angle between nucleosomes N and N+2, which are aligned for both linker lengths (5 or 6 DNA turns), was defined analogously and scaled to 0°–90°:

$$\text{para} = \min\left(\theta, 180^\circ - \theta\right), \quad \text{where } \theta = \arccos(\mathbf{z}_\mathrm{N} \cdot \mathbf{z}_{\mathrm{N+2}}) \times \frac{180^\circ}{\pi}. \tag{12}$$

All metrics were evaluated for every nucleosome triplet in each analyzed frame, and the mean values and standard deviations reported in the figures were obtained by averaging over the full trajectories (Fig. S7b).

#### V. SIMULATION PROTOCOLS

##### A. Molecular dynamics integration

All simulations employed the `LangevinMiddleIntegrator` from OpenMM [13] with timestep $\Delta t = 10$ fs, friction coefficient $\gamma = 0.01$ ps$^{-1}$, and temperature 300 K.

The LangevinMiddle discretization splits the dynamics into deterministic and stochastic substeps, leading to improved configurational sampling compared with other discretizations of Langevin dynamics [13].

##### B. Hamiltonian replica exchange

HREX simulations employed Debye length ($\lambda_D$) as the scaling parameter to enhance configurational sampling across different effective salt concentrations.

**Replica ladders** Debye lengths are sampled using an exponential schedule:

$$\lambda_D(i) = \lambda_\mathrm{low} \left(\frac{\lambda_\mathrm{high}}{\lambda_\mathrm{low}}\right)^{\left(\frac{i}{n-1}\right)^p} \tag{13}$$

where $\lambda_D(i)$ is the Debye length for replica $i$, $\lambda_\mathrm{low} = 0.8$ nm and $\lambda_\mathrm{high} = 1.3$ nm are the lowest and highest Debye lengths, $n = 15$ is the total number of replicas, $i$ is the replica index from 0 to $n-1$, and $p = 1.10$ controls the spacing distribution.

This corresponds to implicit salt concentrations ranging from approximately 150 mM (high salt, low $\lambda_D$) to 60 mM (low salt, high $\lambda_D$).

**Exchange protocol:** Exchange attempts are made every 1,000 MD steps (10 ps) between all adjacent and non-adjacent replica pairs. The acceptance criterion follows the Metropolis algorithm with energy difference $\Delta U$ evaluated from electrostatic term differences, targeting 20–40% acceptance between neighboring replicas.

The implementation uses the openmmtools `ReplicaExchangeSampler` with `LangevinDynamicsMove` [14]. Each move consists of 1,000 integration steps (10 ps) before attempting exchanges.

The algorithm calculates the full matrix containing the energy of each set of coordinates under each Hamiltonian before using Markov Chain Monte Carlo (MCMC) steps to propose swaps. This allows swaps between non-neighboring Hamiltonians, improving diffusion in Hamiltonian space.

Fig. S8 shows the replica exchange acceptance matrices for all 12-nucleosome HREX simulations, confirming efficient sampling across the Debye length ladder.

##### C. Reduced mass molecular dynamics

To accelerate equilibration while preserving equilibrium distributions, we employed reduced mass factors for larger systems: 12-nucleosome arrays use no mass reduction, 108-nucleosome fibers use a mass factor of 0.5, and phase separation systems (324 nucleosomes) use a mass factor of 0.25.

This approach scales down particle masses uniformly, allowing faster dynamics while maintaining the same equilibrium configurational distribution. Equilibrium distributions are preserved because masses cancel in the Boltzmann factor.

According to the canonical distribution, the equilibrium configurational distribution depends only on the potential energy function $U(\mathbf{R})$ and is invariant to particle masses. By reducing masses, inertial timescales decrease and sampling efficiency is enhanced, while the Langevin thermostat ensures correct canonical statistics.

##### D. Direct coexistence setup for phase separation

Phase separation simulations were initialised by selecting 81 uncorrelated configurations from the HREX high-salt replica ($\lambda_D = 0.8$ nm) and placing them in a large cubic grid with 40 nm spacing. The system was then compressed to a dense slab using a Monte Carlo barostat over 30 ns, after which 10 random arrays were displaced to the dilute phase to accelerate equilibration. Production simulations were performed with fixed box dimensions.

The slab geometry ensures that, if phase separation is thermodynamically favorable, the system spontaneously separates into a dense phase and a dilute phase, separated by two planar interfaces orthogonal to the long axis of the box.

#### VI. ANALYSIS METHODS

##### A. Sedimentation coefficient calculation

Hydrodynamic radii $R_h$ were calculated using the Kirkwood approximation applied to nucleosome center-of-geometry positions:

$$R_h = \left( \sum_{i<j} \frac{1}{r_{ij}} \right)^{-1} \tag{14}$$

where the sum runs over all pairs of nucleosome centers.

Sedimentation coefficients were estimated using:

$$S = S_1 \left[ 1 + \frac{2R_1}{N} \sum_{i<j} \frac{1}{r_{ij}} \right] \tag{15}$$

where $N$ is the number of nucleosomes, $S_1 = 11.1$ S is the base sedimentation coefficient for a single nucleosome, and $R_1 = 5.1$ nm is the effective nucleosome radius.

##### B. Contact analysis

*a. Residue–residue contacts* Two residues $i$ and $j$ are in contact if:

$$r_{ij} < r_c = \tfrac{1}{2}(\sigma_i + \sigma_j) + 0.25 \text{ nm} \tag{16}$$

Inter-nucleosomal or non-parental contacts are defined as occurring when a residue from one nucleosome contacts protein or DNA from a different nucleosome. For this purpose, the 147 bp or nucleosomal DNA as well as the full linkers before and after the nucleosomal DNA are considered as part of the parental nucleosome.

*b. Nucleosome–nucleosome contacts* Two nucleosomes are in contact if their centers of geometry are closer than the cutoff distance:

$$d_{ij} = \|\mathbf{r}_i - \mathbf{r}_j\| < 13 \text{ nm} \tag{17}$$

where $\mathbf{r}_i$ and $\mathbf{r}_j$ are the center-of-geometry positions calculated from histone core beads only (excluding flexible tails).

Nucleosome centers of geometry were calculated as the mean positions of core beads in each nucleosome, with tail beads excluded to avoid noise from highly flexible histone tails.

For each nucleosome $k$, the local $\mathbf{z}$ axis was defined using two reference atoms (indices 465 and 952, yields a vector pointing out of the face of the nucleosome):

$$\mathbf{z}_k = \frac{\mathbf{r}_{952}^{(k)} - \mathbf{r}_{465}^{(k)}}{\|\mathbf{r}_{952}^{(k)} - \mathbf{r}_{465}^{(k)}\|} \tag{18}$$

Relative orientations were then quantified by:

$$\alpha = \cos^{-1}\big(\mathbf{z}_i \cdot \mathbf{z}_j\big) \tag{19}$$

$$\beta_i = \cos^{-1}\left(\frac{\mathbf{r}_{ij}\cdot\mathbf{z}_i}{\|\mathbf{r}_{ij}\|}\right)$$
$$\beta_j = \cos^{-1}\left(\frac{\mathbf{r}_{ij}\cdot\mathbf{z}_j}{\|\mathbf{r}_{ij}\|}\right) \tag{20}$$

Contacts were classified as:

- **Face–face**, if ($\alpha < 45°$ or $\alpha > 135°$) and at least one of ($\beta_i, \beta_j$) lies outside $[45°, 135°]$, or if $d_{ij} < 11$ nm
- **Side–side**, if ($\alpha < 45°$ or $\alpha > 135°$) and both $\beta_i, \beta_j \in [45°, 135°]$, with $d_{ij} \geq 11$ nm
- **Face–side**, otherwise

These categories capture the most common relative orientations observed in fibers and mirror terminology used in structural studies. The algorithm used here is adapted from our previous work [1].

*c. Energy weighting* Energy-weighted contacts sum the full pairwise interaction energy for each contact using the complete Hamiltonian energy terms (both attractive and repulsive contributions):

$$E_{\text{contact}} = U_{\text{AH}}(d_{ij}) + U_{\text{ele}}(d_{ij}) \tag{21}$$

where $d_{ij}$ is the contact distance, $U_{\text{AH}}$ is the Ashbaugh–Hatch potential defined in Section 1.2, $U_{\text{ele}}$ is the Debye–Hückel electrostatic potential defined in Section 1.2, and atom type parameters ($\epsilon_{ij}$, $\sigma_{ij}$, $\lambda_{ij}$, $q_i$, $q_j$) are assigned according to the bead types involved in the contact.

We use two different cutoff distances for energy weighting: the standard close-contact cutoff, $r_c = \frac{1}{2}(\sigma_i + \sigma_j) + 0.25$ nm, and the longer Debye-range cutoff of 1.6 nm ($2\lambda_D$) for the comparison of WT and acetylated lysine sites. The Debye-range calculation used a cutoff distance twice the value of the simulated $\lambda_D$, aiming to more fully capture the energetic impact of specific site acetylations. Fig. S17 explicitly compares close-contact energy-weighted contact frequencies and raw, unweighted contact frequencies at both the close-contact and Debye-range cutoffs to supplement the Debye-range energy-weighted lysine interaction-energy contributions displayed in Fig. 8c.

##### C. Radius of gyration

The gyration radius $R_g$ of chromatin fibers/fragments was calculated from nucleosome center-of-geometry positions $\{\mathbf{r}_i\}$ as:

$$R_g = \sqrt{\frac{1}{N}\sum_{i=1}^{N}\|\mathbf{r}_i - \mathbf{r}_{\text{CoG}}\|^2} \tag{22}$$

where $N$ is the number of nucleosomes and $\mathbf{r}_{\text{CoG}}$ is the overall center of geometry of the fiber. Defining $R_g$ in terms of nucleosome centers reflects the global folding of the fiber, rather than local flexibility of individual residues.

##### D. DNA accessibility quantification

For each linker DNA segment, we count particles within a 5 nm shell around the linker's excluded volume virtual sites. Let $Q_\ell = \{\mathbf{q}_k\}$ be the set of excluded volume bead positions for linker $\ell$ and let $H$ be the set of atoms considered as neighbors (any protein bead or excluded volume bead not part of the same linker).

An atom $i \in H$ is counted if its minimum distance to the linker's excluded volume set lies within the shell:

$$\min_{\mathbf{q}\in Q_\ell} \|\mathbf{x}_i - \mathbf{q}\| \in [0,\ 5\ \text{nm}] \tag{23}$$

Let $N_{\ell,t}$ be the number of unique atoms satisfying this condition at frame $t$. The shell volume is:

$$V_{\text{shell}} = \tfrac{4}{3}\pi(5\ \text{nm})^3 \tag{24}$$

The instantaneous density is:

$$\rho_{\ell,t} = \frac{N_{\ell,t}}{V_{\text{shell}}} \tag{25}$$

The time-averaged exposure density per linker is:

$$\bar{\rho}_\ell = \frac{1}{T}\sum_{t\in\mathcal{T}} \rho_{\ell,t} \tag{26}$$

where $\mathcal{T}$ is the set of sampled frames and $T = |\mathcal{T}|$.

This quantity should be interpreted as a relative measure of accessibility rather than a strict physical density, as overlapping volumes around multiple beads are not explicitly accounted for.

##### E. Structural clustering of tetramers

For each four-nucleosome structure, we compute six pairwise inter-nucleosomal distances $\{d_{12}, d_{13}, d_{14}, d_{23}, d_{24}, d_{34}\}$ using center-of-geometry positions for each nucleosome.

These are transformed into symmetrised features invariant to label permutation $(1, 2, 3, 4) \leftrightarrow (4, 3, 2, 1)$:

$$s_1 = d_{12} + d_{34} \tag{27}$$
$$s_2 = d_{13} + d_{24} \tag{28}$$
$$s_3 = d_{14} \tag{29}$$
$$s_4 = d_{23} \tag{30}$$
$$s_5 = d_{12}d_{13} + d_{24}d_{34} \tag{31}$$
$$s_6 = d_{12}d_{13}^2 + d_{24}^2 d_{34} \tag{32}$$

This transformation follows the approach of Ding and Zhang [15].

**Clustering procedure:** K-means clustering (k=3) was applied to the six-dimensional symmetrized distance features using the SciPy implementation. The three clusters correspond to physically interpretable configurations: compact stacked structures with two nucleosome–nucleosome contacts (Cluster 1), intermediate structures with one contact (Cluster 2), and extended structures with no contacts (Cluster 3).

Principal component analysis (PCA) was performed on the same feature set for visualization, with the first two components capturing major structural variations.

##### F. Connected component analysis

To quantify chromatin array clustering, we performed graph-based connected component analysis. Two arrays are considered connected if any pair of nucleosomes (one from each array) have centers within 13 nm of each other. The largest connected component was identified as the maximum set of arrays that are transitively connected through direct or bridging interactions in each frame.

##### G. Density profile analysis

For direct coexistence simulations, density profiles along the slab axis were calculated as follows. Coordinates were first mapped so that the center of each molecule lay within the same periodic box. Each frame was then centered so that the overall center of geometry of all particles was placed at the origin, and $x$-coordinates were re-wrapped to ensure all particles fell within the analysis box. Particles were binned into a histogram along the $x$ axis, and density profiles from each frame were averaged to yield the final mean density profile. Interfacial properties were extracted by fitting each of the two slab boundaries locally to a hyperbolic tangent profile,

$$\rho(x) = \frac{1}{2}(\rho_{\text{dilute}} + \rho_{\text{dense}}) + \frac{1}{2}(\rho_{\text{dense}} - \rho_{\text{dilute}}) \tanh\left(\frac{x - x_0}{w}\right), \tag{33}$$

where $\rho_{\text{dilute}}$ and $\rho_{\text{dense}}$ are the fitted dilute- and dense-phase plateau densities, $x_0$ is the interface position, and $w$ is the interfacial width parameter. The mean density profile was smoothed with a Gaussian filter ($\sigma = 1$ bin) only to locate the two interfaces from the largest absolute gradients. Around each gradient peak, the unsmoothed density profile was fitted in a local window of $\pm 5$ bins. The reported interface width is the average of $|w|$ from the two local interface fits. Plateau densities were then calculated from the dense region between the two interfaces and the dilute region outside them, excluding $1.5|w|$ around each fitted interface.

### VII. PERFORMANCE BENCHMARKING

#### A. Hardware specifications

Benchmarks were performed on CPU nodes (Intel Cascade Lake with 56 cores for 12-nucleosome systems and AMD EPYC 7742 with 128 cores for 108-nucleosome systems) and GPUs (NVIDIA RTX 4080 consumer GPU with 16GB VRAM and NVIDIA H100 data center GPU).

#### B. Benchmark systems

Two system sizes were benchmarked: a small system comprising a 12-nucleosome fiber ($\sim$18,000 particles) and a large system comprising a 108-nucleosome fiber ($\sim$190,000 particles).

#### C. Performance metrics

Performance was measured as timesteps per second, averaged over 1,000,000 integration steps after equilibration. Performance comparisons for 12-nucleosome and 108-nucleosome systems across different hardware configurations are shown in main text Fig. 1. A single H100 GPU outperforms a 56-core CPU node by $\sim 9\times$ for 12-nucleosome systems and outperforms a 128-core CPU node by $>10\times$ for 108-nucleosome systems. For smaller systems, NVIDIA's Multi-Process Service can be used to run multiple simulation instances in parallel, increasing overall throughput.

The dramatic performance improvement enables simulation of systems that were previously computationally prohibitive. The OpenMM implementation can outperform CPU-only LAMMPS by at least an order of magnitude, with the gap widening as system size increases. Part of this difference arises from spatial domain decomposition in LAMMPS, which is highly efficient for dense bulk systems but can be wasteful for implicit-solvent coarse-grained simulations where large regions of the simulation box are empty [16].

Regarding GPU memory requirements, the benchmarked OpenCGChromatin simulations used negligible VRAM relative to modern accelerator capacity. Single-replica 12-nucleosome simulations used approximately 0.25 GiB on the RTX 4080 workstation and 0.52 GiB on an H100, while the 16-replica H100 MPS benchmark used 8.34 GiB in total including the MPS server, and the 108-nucleosome H100 benchmark used approximately 0.60 GiB (Table S3). These values include CUDA/OpenMM context overhead measured by `nvidia-smi`, which dominates the memory footprint for these benchmark systems. Thus, the Fig. 1 benchmark simulations are not limited by GPU memory; performance is instead governed by compute throughput.

### VIII. SOFTWARE AND COMPUTATIONAL RESOURCES

#### A. Simulation software

All simulations used OpenMM 8.0 [17, 18] with the custom OpenCGChromatin Python package. OpenCGChromatin is built on OpenMM's Python API and provides high-level classes for chromatin system construction, automated force field setup, integration with enhanced sampling methods (HREX via openmmtools), and utilities for post-translational modification.

Production simulations ran on NVIDIA H100 GPUs using the CUDA platform.

OpenMM allows bonded and non-bonded energy terms to be specified as algebraic expressions, which are symbolically differentiated to yield force expressions before just-in-time compilation converts them into efficient machine code. This enables arbitrary force field terms to be quickly prototyped and implemented with high performance.

#### B. Analysis software

Trajectory analysis employed:

- MDTraj for trajectory manipulation
- NumPy and SciPy for numerical computations

- scikit-learn for k-means clustering and PCA
- Matplotlib for data visualization

##### C. Visualization

Molecular graphics were rendered using VMD 1.9.4 with GPU-accelerated Tachyon ray tracer [19].

#### IX. MINIMAL MODEL SIMULATIONS

The histone core is represented by a single ellipsoidal bead with semi-axes $28 \times 28 \times 20\,\text{Å}$. Linker and nucleosomal DNA are modeled as finite-size, orientable spheres (ellipsoids with semi-axes $12 \times 12 \times 12\,\text{Å}$), with one DNA bead per 5 base pairs (Fig. S14a). Nucleosome–nucleosome and nucleosome–DNA interactions are described by orientation-dependent pair potentials fitted to reproduce internucleosome potentials of mean force obtained from the chemically specific model introduced by Farr et al. [1]. All model parameters and energy functions follow those used previously [1]. Direct coexistence simulations containing 200 copies each of four-nucleosome chromatin arrays were prepared using the method described in our previous work [1]. Surface tension was measured from minimal model direct coexistence simulations using the following expressions:

$$\gamma \;=\; \frac{1}{2}\int_{-L_x/2}^{L_x/2}\big[\,P_N(x) - P_T(x)\,\big]\,\mathrm{d}x, \qquad P_N \equiv P_{xx}, \;\; P_T \equiv \tfrac{1}{2}\big(P_{yy} + P_{zz}\big). \tag{34}$$

$$\gamma \;=\; \frac{L_x}{2N}\Big\langle P_N - P_T \Big\rangle. \tag{35}$$

where $L_x$ is the length of the long side of the simulation box, $N$ is the number of droplets in the simulation, and $P_N$ and $P_T$ are the normal and tangential components of the pressure tensor with respect to the interface. The tangential component of the pressure tensor is averaged over both tangential directions.

To characterize viscoelasticity in the minimal-model chromatin condensates, we evaluate the shear stress relaxation modulus $G(t)$ in the linear-response (zero-strain) limit using the expression:

$$G(t) \;=\; \frac{V}{k_\mathrm{B} T}\,\big\langle \sigma_{\alpha\beta}(t)\,\sigma_{\alpha\beta}(0)\big\rangle, \qquad \alpha \neq \beta \in \{x, y, z\}, \tag{36}$$

where $V$ is the system volume, $T$ the temperature, and $k_\mathrm{B}$ the Boltzmann constant. The average $\langle\cdot\rangle$ is taken over equilibrium time origins; in practice we improve statistics by averaging Eq. (36) over $(xy, yz, zx)$, using the following full expression:

$$\begin{aligned} G(t) = \frac{V}{5k_\mathrm{B}T}\Big[&\langle\sigma_{xy}(0)\,\sigma_{xy}(t)\rangle + \langle\sigma_{xz}(0)\,\sigma_{xz}(t)\rangle \\ &+ \langle\sigma_{yz}(0)\,\sigma_{yz}(t)\rangle + \frac{1}{6}\big(\langle N_{xy}(0)\,N_{xy}(t)\rangle + \langle N_{xz}(0)\,N_{xz}(t)\rangle + \langle N_{yz}(0)\,N_{yz}(t)\rangle\big)\Big]. \end{aligned} \tag{37}$$

where $N_{\alpha\beta} = \sigma_{\alpha\alpha} - \sigma_{\beta\beta}$ is the normal stress difference [20].

The correlation functions in Eq. (37) are accumulated on the fly during the simulation, introducing negligible computational overhead and avoiding postprocessing. Once $G(t)$ is known, the zero-shear viscosity follows directly from the Green–Kubo relation,

$$\eta_0 \;=\; \int_0^\infty G(t)\,\mathrm{d}t. \tag{38}$$

In practice, we evaluated this integral by direct numerical integration for short times ($t < 10^{-10}$ s). From $10^{-10}$ s on, we first fit $G(t)$ to the first three Maxwell modes and integrated the fitted expression.

#### SUPPLEMENTARY TABLES

TABLE S1. **Optimized protein–DNA and DNA–DNA interaction parameters.** Ashbaugh–Hatch parameters for protein interactions with DNA beads and virtual sites. Protein–protein parameters follow the Kim–Hummer parameterization [7]. Energy scale $\epsilon = 4.184$ kJ mol$^{-1}$ for all protein–DNA interactions. Electrostatic cutoff $r_c = 4.5\lambda_D$.

| Interaction | Description | $\sigma$ (nm) | $\epsilon$ (kJ mol$^{-1}$) | $\lambda$ |
|---|---|---|---|---|
| Protein–C1' | DNA backbone bead | 0.575 | 4.184 | 0.0025 |
| Protein–Phosphate | Virtual site, charged | 0.500 | 4.184 | 0.0250 |
| Protein–Excluded Volume | Major groove protection | 0.900 | 4.184 | 0.0025 |
| DNA–DNA (all) | No non-bonded interactions | — | 0 | — |

TABLE S2. **Simulation system details.** Summary of all simulated systems including size, composition, simulation time, and purpose. HREX = Hamiltonian replica exchange with Debye length scaling.

| System | $N_{\text{nuc}}$ | $N_{\text{particles}}$ | Linker (bp) | Box (nm$^3$) | Time ($\mu$s) | Salt (mM) | Purpose |
|---|---|---|---|---|---|---|---|
| 12-nuc arrays | 12 | ~18,000 | 18–63 | — | 5.0 | 5 | Low-salt sedimentation validation |
| 12-nuc arrays | 12 | ~18,000 | 18–63 | — | 1.5 | 60, 150 | Sedimentation validation (HREX) |
| 108-nuc fiber (WT) | 108 | ~190,000 | 22 | — | 100 | 150 | Control |
| 108-nuc fiber (5×12 Ac) | 108 | ~190,000 | 22 | — | 100 | 150 | Patterned acetylation |
| 108-nuc fiber (54 Ac) | 108 | ~190,000 | 22 | — | 100 | 150 | Block acetylation |
| Tetramer (25 bp) | 4 | ~6,000 | 25 | — | 3 | 60–150 | HREX ensemble |
| Tetramer (30 bp) | 4 | ~6,000 | 30 | — | 3 | 60–150 | HREX ensemble |
| Phase sep. (25 bp) | 324 | ~600,000 | 25 | 500×83×83 | 20 | 100 | Direct coexistence |
| Phase sep. (30 bp) | 324 | ~600,000 | 30 | 500×83×83 | 20 | 100 | Direct coexistence |

TABLE S3. **Performance benchmarks across hardware configurations.** Simulation throughput measured as timesteps per second for different system sizes and hardware, with GPU memory footprints where applicable. MPS = NVIDIA Multi-Process Service running 16 independent 12-nucleosome simulations concurrently on a single H100 GPU. Speedup calculated relative to CPU baseline for each system size. Performance data from main text Fig. 1; reported GPU memory is approximate process-resident VRAM measured with `nvidia-smi` during production.

| System | Hardware | $N_{\text{particles}}$ | Performance | Speedup | GPU memory (GiB) |
|---|---|---|---|---|---|
| 12-nucleosome | Intel Cascade Lake (56 cores) | ~18,000 | baseline | 1.0× | — |
| 12-nucleosome | NVIDIA RTX 4080 | ~18,000 | — | ~9× | 0.25 |
| 12-nucleosome | NVIDIA H100 | ~18,000 | — | ~9× | 0.52 |
| 12-nucleosome | NVIDIA H100 (16× MPS) | ~18,000 | — | ~27× total throughput | 8.34 total; 0.52 per replica |
| 108-nucleosome | AMD EPYC 7742 (128 cores) | ~190,000 | baseline | 1.0× | — |
| 108-nucleosome | NVIDIA H100 | ~190,000 | — | >10× | 0.60 |

TABLE S4. **Partial charges for all bead types.** Charges assigned to protein residues, modified residues, and DNA components in the OpenCGChromatin model.

| Bead Type | Charge ($e$) |
|---|---|
| *Neutral amino acids* | |
| Ala, Cys, Gly, Ile, Leu, Met, Phe, Pro, Ser, Thr, Trp, Tyr, Val | 0 |
| Asn, Gln | 0 |
| *Charged amino acids* | |
| Arginine (Arg) | +1 |
| Lysine (Lys) | +1 |
| Aspartic acid (Asp) | −1 |
| Glutamic acid (Glu) | −1 |
| Histidine (His) | +0.5 |
| *Modified residues* | |
| Acetylated lysine (Ac-Lys) | 0 |
| *DNA components* | |
| C1' bead | 0 |
| Phosphate virtual site | −1 |
| Excluded volume virtual site | 0 |

TABLE S5. **Sequences of the miniABC dataset.** The 13 DNA sequences (18 nucleotides each) used for DNA model validation, covering all possible base pair tetramer combinations [12].

| Sequence | DNA Sequence |
|---|---|
| 1 | GCAACGTGCTATGGAAGC |
| 2 | GCAATAAGTACCAGGAGC |
| 3 | GCAGAAACAGCTCTGCGC |
| 4 | GCAGGCGCAAGACTGAGC |
| 5 | GCATTGGGGACACTACGC |
| 6 | GCGAACTCAAAGGTTGGC |
| 7 | GCGACCGAATGTAATTGC |
| 8 | GCGGAGGGCCGGGTGGGC |
| 9 | GCGTTAGATTAAAATTGC |
| 10 | GCTACGCGGATCGAGAGC |
| 11 | GCTGATATACGATGCAGC |
| 12 | GCTGGCATGAAGCGACGC |
| 13 | GCTTGTGACGGCTAGGGC |

TABLE S6. **Block-averaged dense-phase, dilute-phase, and interfacial properties of the 25 and 30 bp tetranucleosome direct-coexistence simulations.** For each of the two systems, the analysis window of main text Fig. 4 is split into five non-overlapping blocks; each row reports a single block, and the bold summary row gives the across-block mean and sample standard deviation (mean ± SD, $N = 5$). $\rho_{\text{dense}}$ and $\rho_{\text{dilute}}$ are the plateau nucleosome concentrations on the dense and dilute sides of the slab respectively, and $w$ is the hyperbolic-tangent interfacial width parameter fitted independently on each block.

| Linker | Block (frames) | $\rho_{\text{dense}}$ ($\mu$M) | $\rho_{\text{dilute}}$ ($\mu$M) | $w$ (nm) |
|---|---|---|---|---|
| 25 bp | 1 | 781.6 | 8.74 | 16.81 |
| | 2 | 832.5 | 9.63 | 16.59 |
| | 3 | 768.7 | 13.19 | 16.45 |
| | 4 | 758.5 | 8.78 | 22.05 |
| | 5 | 757.2 | 9.36 | 18.64 |
| | **mean ± SD** | 779.7 ± 31.1 | 9.94 ± 1.85 | 18.11 ± 2.38 |
| 30 bp | 1 | 472.4 | 70.72 | 24.98 |
| | 2 | 312.0 | 82.06 | 32.96 |
| | 3 | 378.5 | 97.67 | 3.80 |
| | 4 | 569.5 | 102.43 | 14.55 |
| | 5 | 378.0 | 89.26 | 32.72 |
| | **mean ± SD** | 422.1 ± 100.3 | 88.43 ± 12.61 | 21.80 ± 12.56 |

#### SUPPLEMENTARY FIGURES

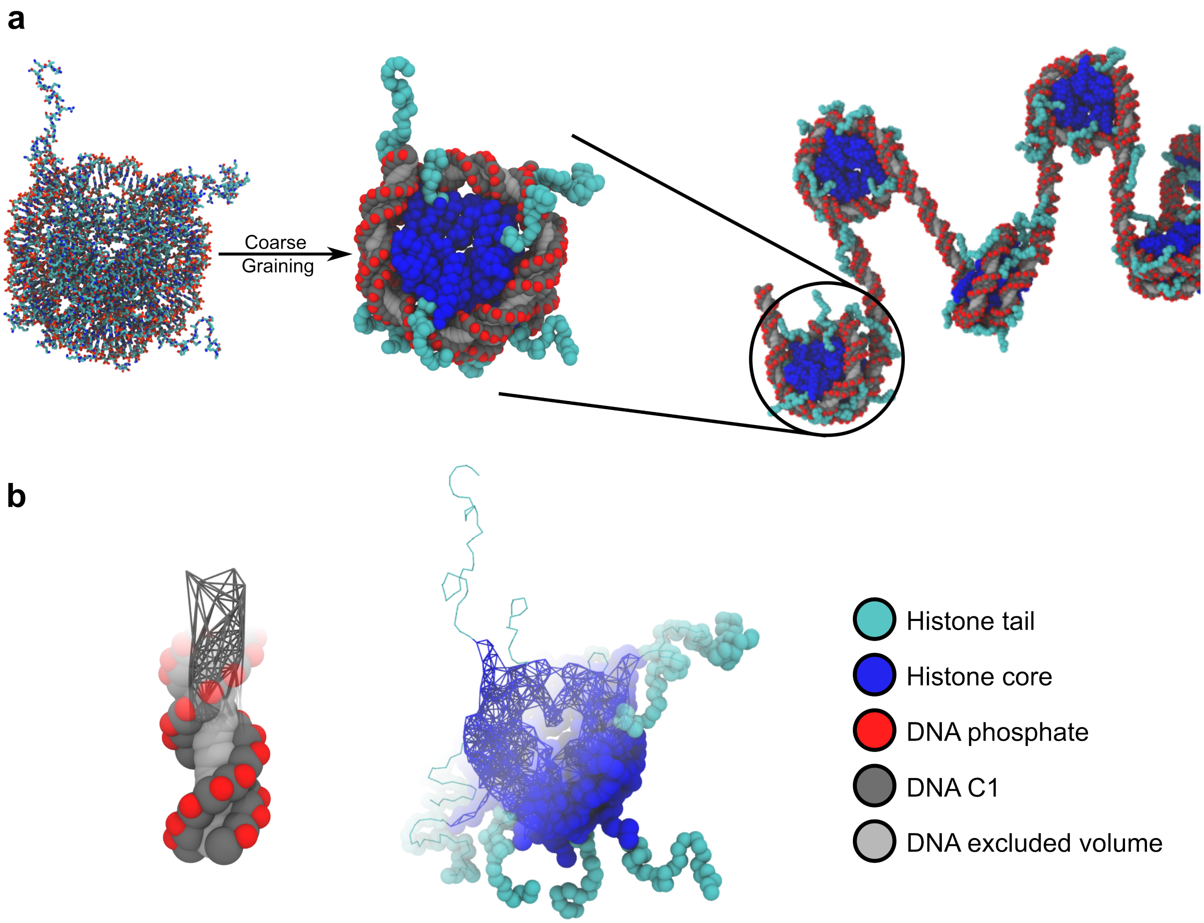


FIG. S1. **Coarse-grained representation of the nucleosome and chromatin fiber.** (a) Side-by-side visualization of all-atom and coarse-grained nucleosomes, together with a coarse-grained chromatin fiber. (b) Cut-away bead-and-bond views of the histone core and a short DNA segment, illustrating the coarse-grained sites and harmonic-bond connectivity.

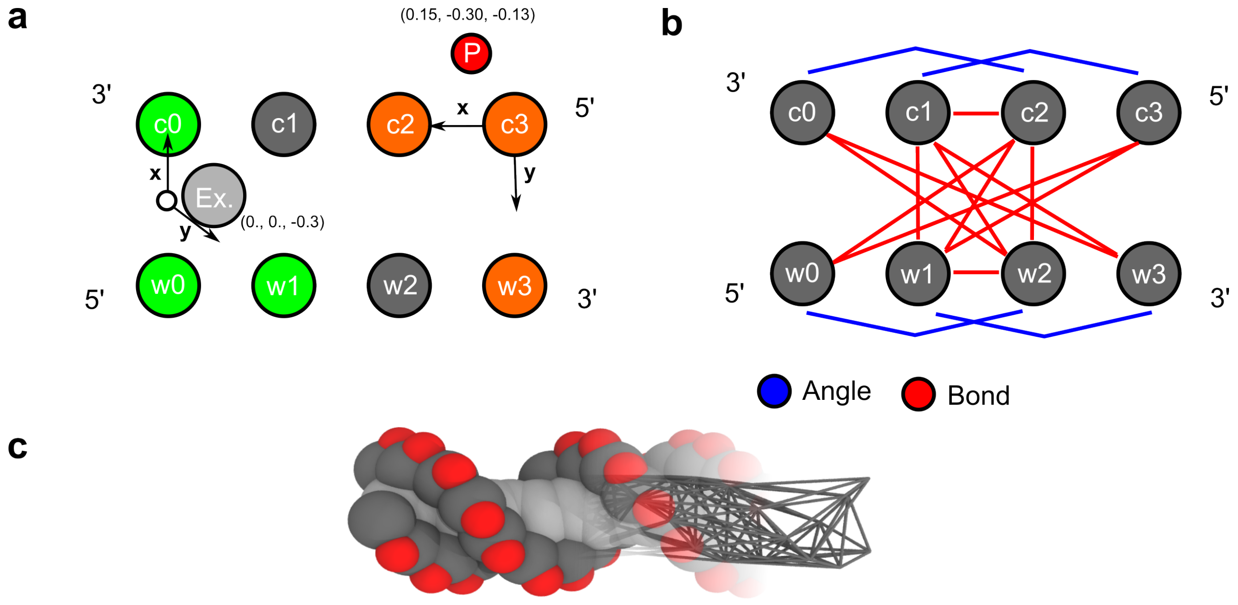


FIG. S2. **Virtual site definitions and tetramer bonded terms in the CGeNArate DNA model.** (a) Diagrams showing the geometric placement of phosphate and excluded volume virtual sites relative to C1′ reference beads, defined using local coordinate systems. Phosphate sites carry negative charges for electrostatic interactions, while excluded volume sites provide steric protection in the major groove. (b) Schematic of sequence-dependent bonds and angles within a base pair tetramer, illustrating the sequence-specific bonds and angles that capture DNA flexibility and helical geometry.

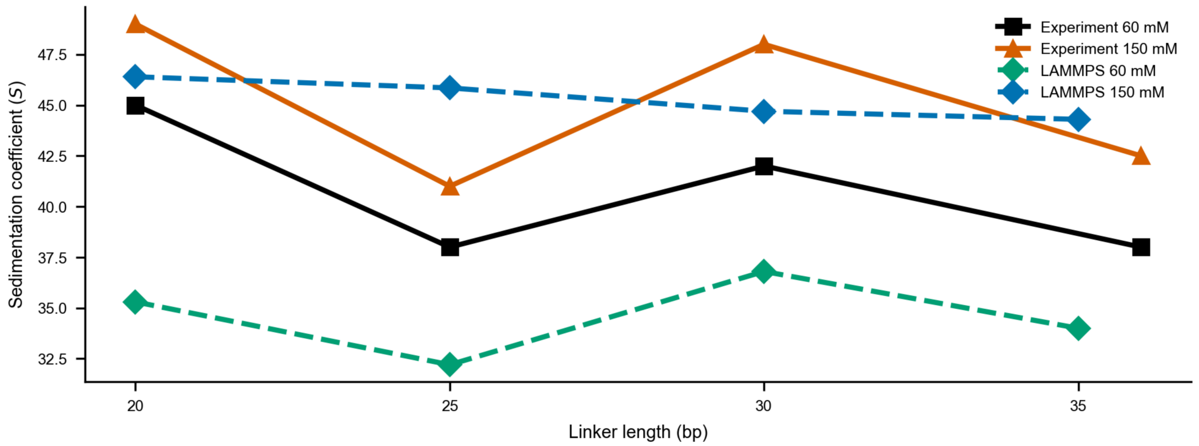


FIG. S3. **OpenCGChromatin resolves failure of previous parameterization to capture experimental compaction periodicity.** Sedimentation coefficients for 12-nucleosome arrays with regular linker lengths spaced at 5 bp increments at 150 mM [21] show that the previous model [1] fails to capture the characteristic $10N$ versus $10N{+}5$ bp oscillations observed experimentally [10, 11]. OpenCGChromatin (main text Fig. 2a) accurately reproduces these oscillations through improved protein–DNA parameterization.

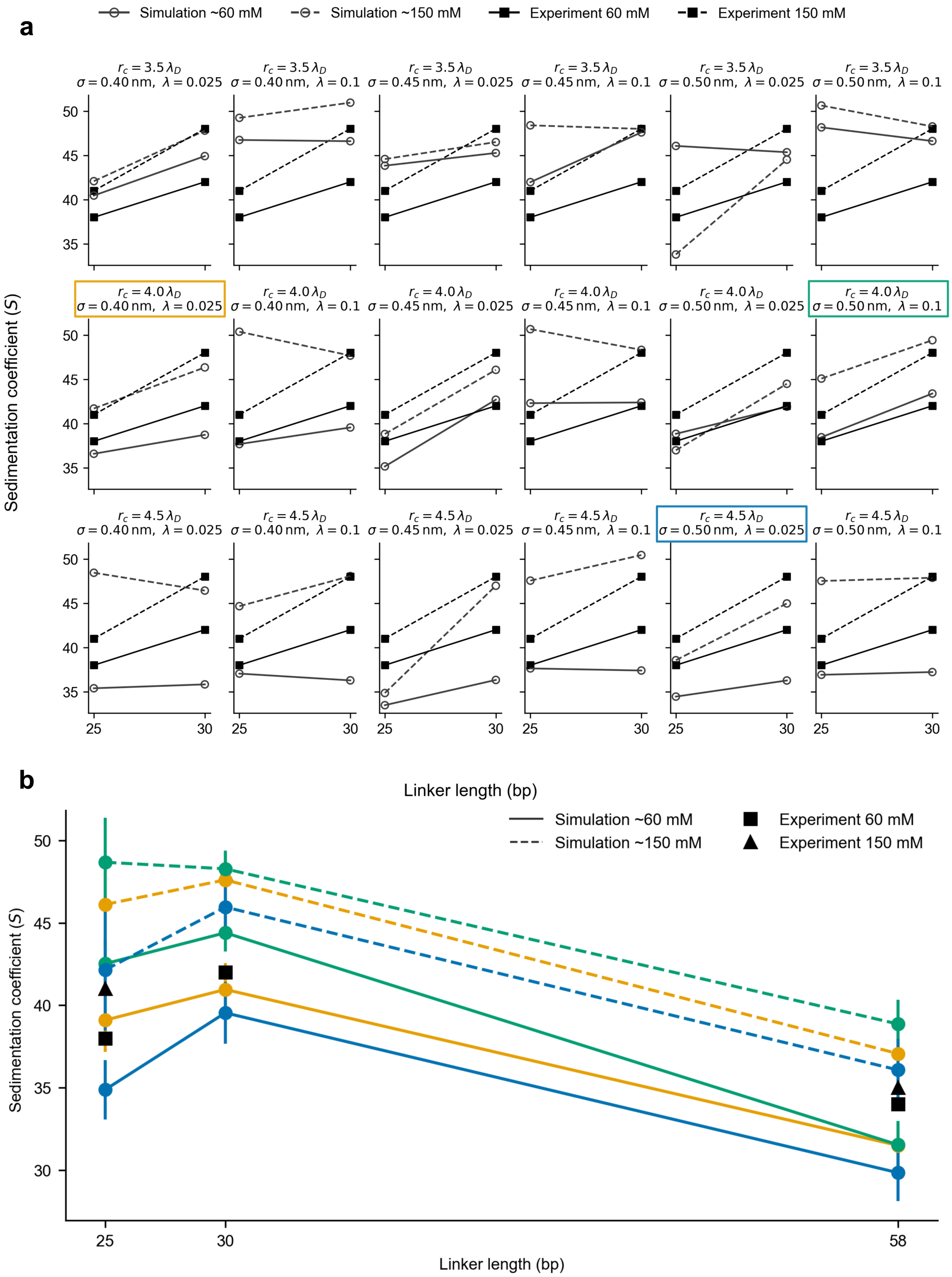


FIG. S4. **Protein–DNA parameterization grid and validation.** (a) Initial parameter screening using regular MD simulations of 12-nucleosome arrays with 25 and 30 bp linkers at 150 and 60 mM salt. The grid explores combinations of protein–phosphate $\sigma$ (0.40, 0.45, 0.50 nm), hydrophobicity $\lambda$ (0.10, 0.025), and electrostatic cutoff (3.5, 4.0, 4.5$\times\lambda_D$). (b) Refined validation using Hamiltonian replica exchange (HREX) simulations for selected parameter sets across 25, 30, and 58 bp linkers. Final parameters were chosen based on agreement with experimental sedimentation coefficients and reproduction of the $10N$ versus $10N$+5 periodicity.

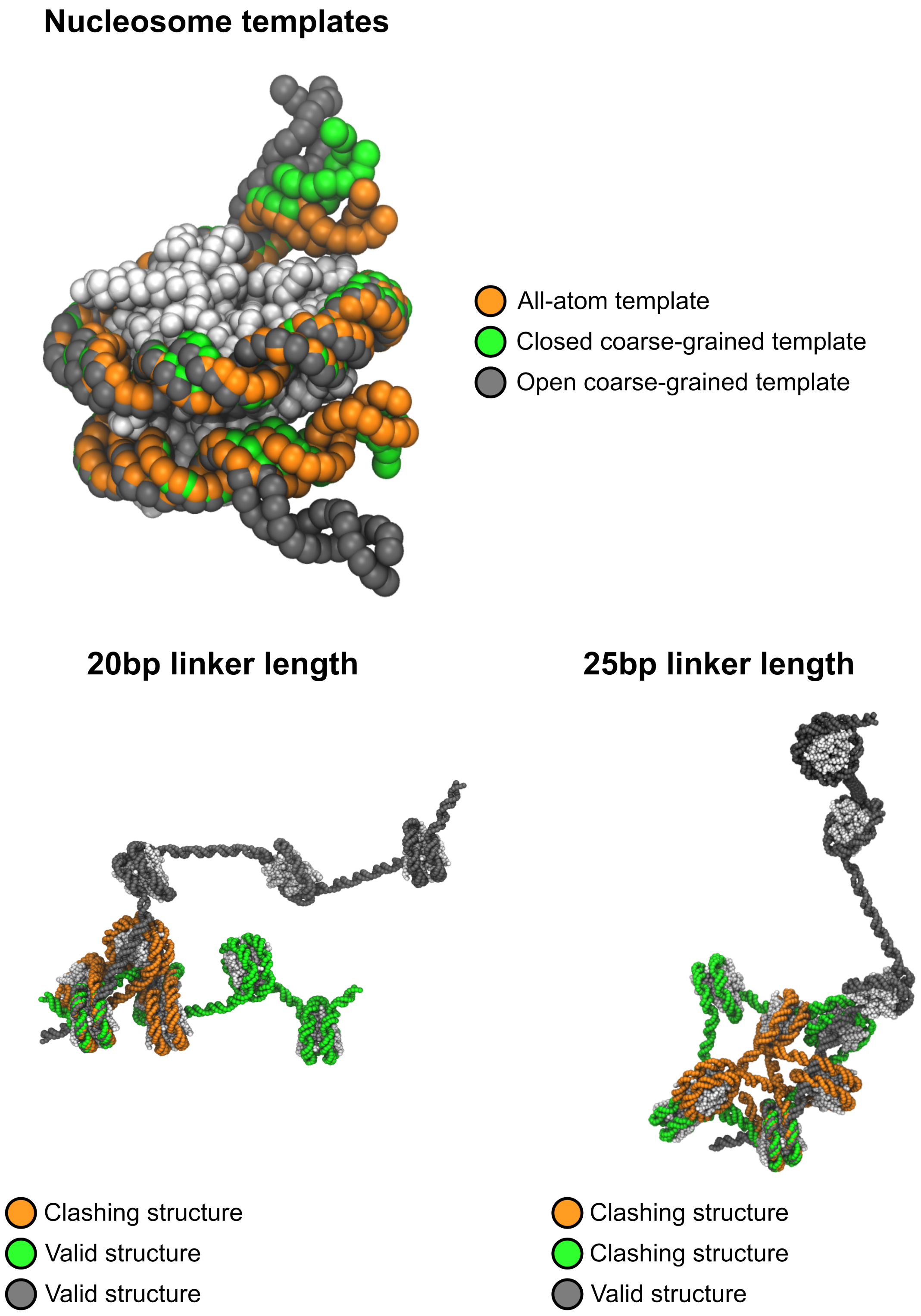


FIG. S5. **Schematic of chromatin array construction.** Illustration showing the three nucleosome templates used in system building: (i) atomistic template with ∼40° entry/exit angle (derived from PDB 1KX5, suitable only for mononucleosomes due to steric clashes), (ii) closed coarse-grained template with ∼90° entry/exit angle (suitable for most linker lengths), and (iii) open coarse-grained template (required for $10N+5$ linkers such as 25 and 36 bp to avoid out-of-phase steric clashes). The geometric construction procedure accounts for B-form DNA helical twist to position consecutive nucleosomes with appropriate relative orientations.

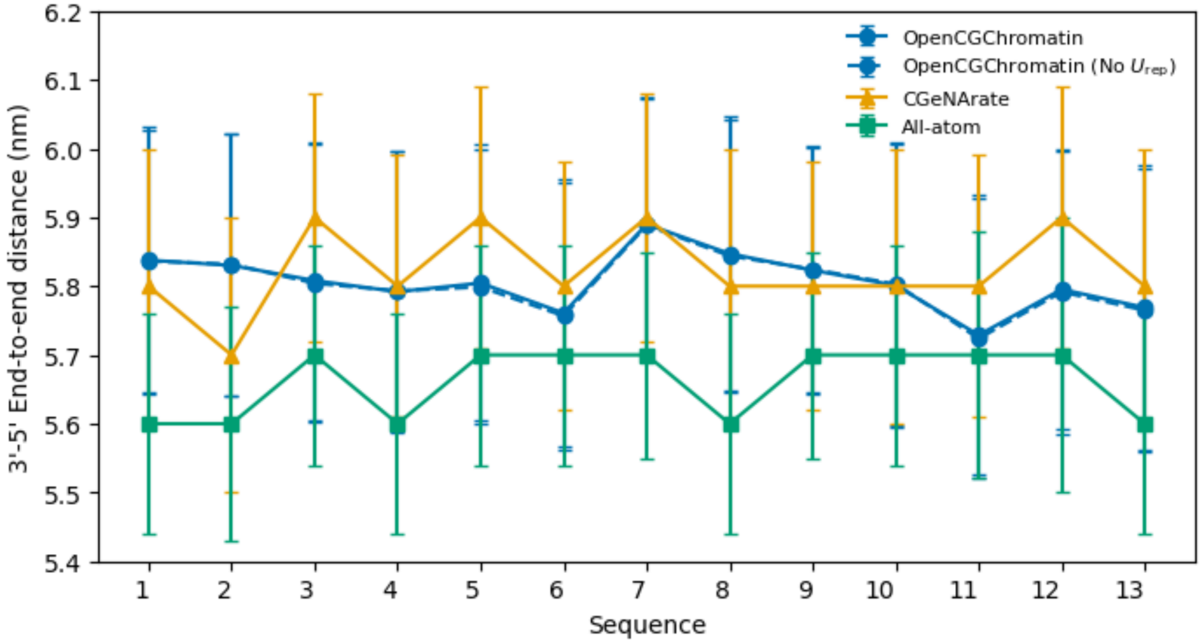


FIG. S6. **DNA model validation using the miniABC dataset.** Comparison of end-to-end distance distributions for 13 DNA sequences (18 nucleotides each) covering all possible base pair tetramer combinations. Four models are compared: (i) the original CGeNArate coarse-grained dsDNA model, (ii) all-atom MD simulations that CGeNArate was parameterized to reproduce, (iii) OpenCGChromatin DNA model with repulsive potential $U_{rep}$ between consecutive excluded volume sites, and (iv) OpenCGChromatin DNA model without $U_{rep}$. The close agreement across all models confirms that our modifications (virtual sites, protein–DNA interactions, and $U_{rep}$) preserve sequence-dependent DNA mechanical properties.

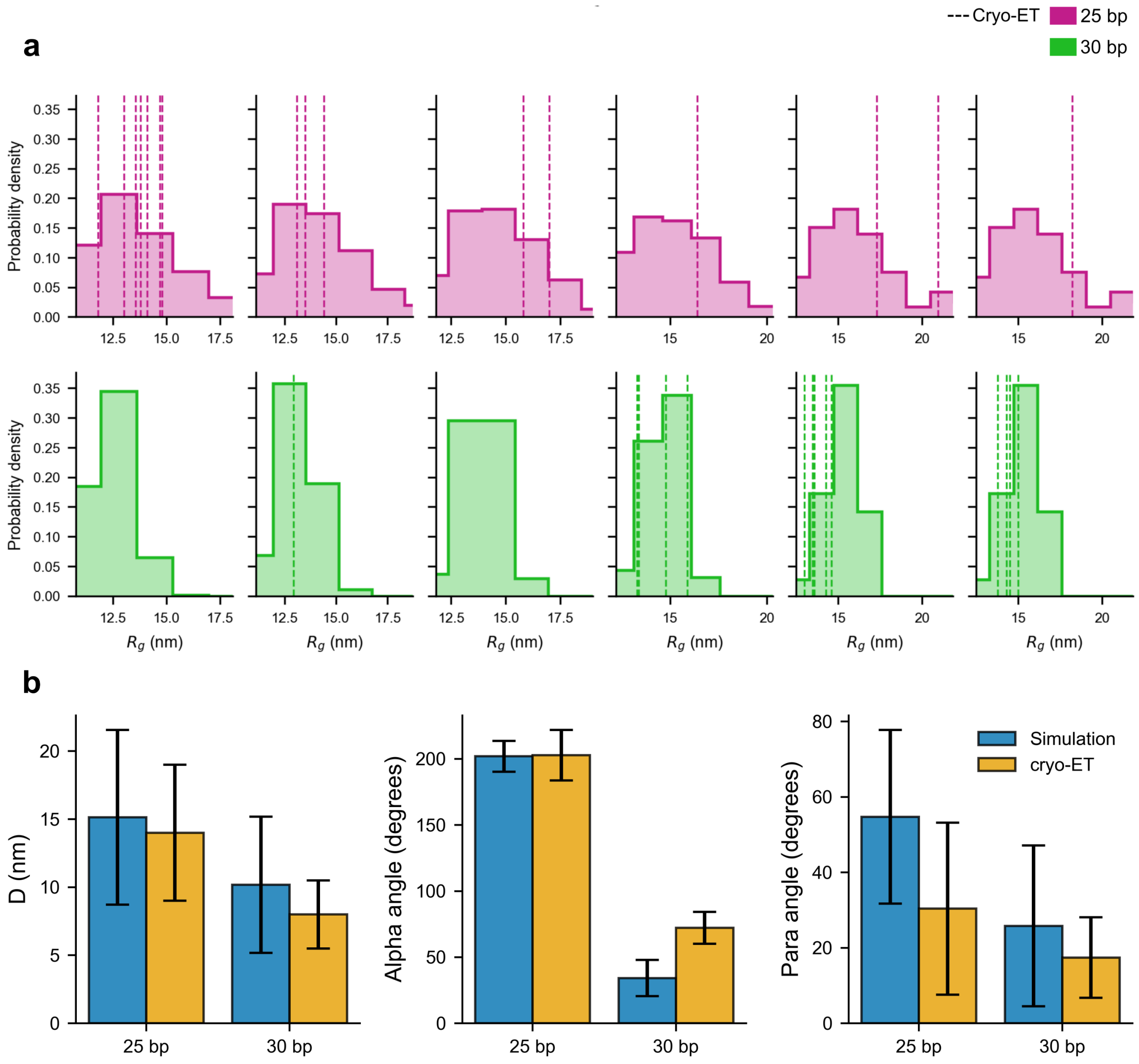


FIG. S7. **Cryo-ET fiber validation demonstrates accurate reproduction of chromatin compaction and structural metrics.** (a) Fragment-wise radius of gyration ($R_g$) analysis reproduces the 25 bp > 30 bp compaction ordering observed experimentally. For each fragment length $k \in \{7, \ldots, 12\}$ nucleosomes, histograms show simulated $R_g$ distributions calculated from contiguous $k$-nucleosome windows within 12-nucleosome arrays for 25 bp (magenta) and 30 bp (green) linkers at 150 mM salt; vertical lines mark individual cryo-ET fragment measurements [9]. Across all fragment lengths, 25 bp chromatin consistently exhibits larger $R_g$ values than 30 bp chromatin, and experimental measurements fall within or close to the simulated distribution ranges. (b) Comparison of simulated structural metrics with those calculated for cryo-ET fibers [9]. Metrics shown are: $D$, the center-of-mass distance between second-nearest-neighbor nucleosomes; $\alpha_{i,i+1}$, the dihedral angle between the planes of nearest-neighbor nucleosomes; and $\text{para}_{i,i+2}$, the dihedral angle between the planes of second-nearest-neighbor nucleosomes. Mean values and standard deviations are shown for both simulations and experiments at 150 mM salt.

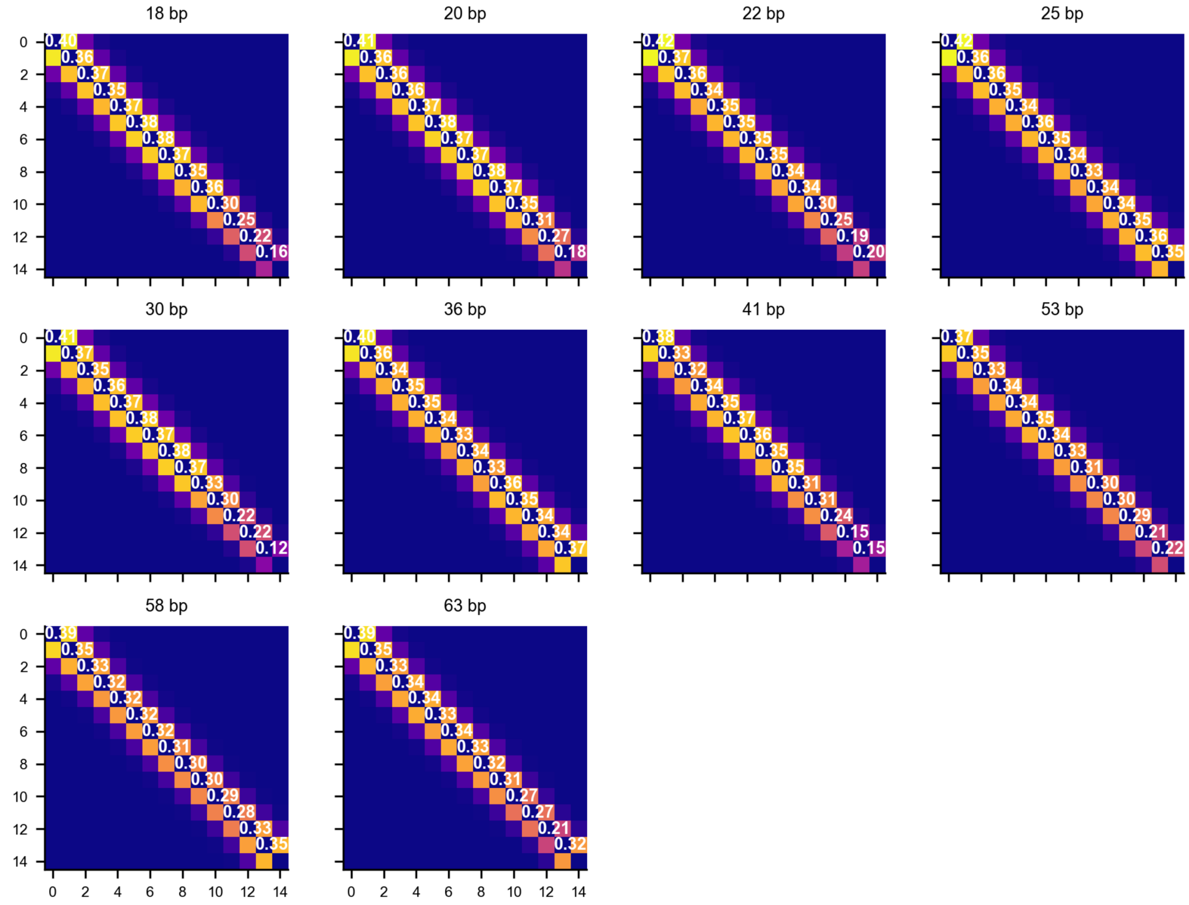


FIG. S8. **Hamiltonian replica exchange acceptance probability matrices demonstrate efficient sampling.** Exchange acceptance probabilities between all replica pairs for representative HREX simulations of 12-nucleosome arrays with different linker lengths. Fifteen replicas span Debye lengths from 0.8 nm (150 mM salt) to 1.3 nm (60 mM salt) using an exponential spacing distribution. The matrices show efficient exchange between neighboring replicas, with typical acceptance rates of 20–40% for adjacent pairs, confirming that the chosen Debye length ladder enables adequate sampling across configurational and salt-concentration space. Higher acceptance rates at lower replica indices reflect the weaker salt-dependence of compact states at high salt concentrations.

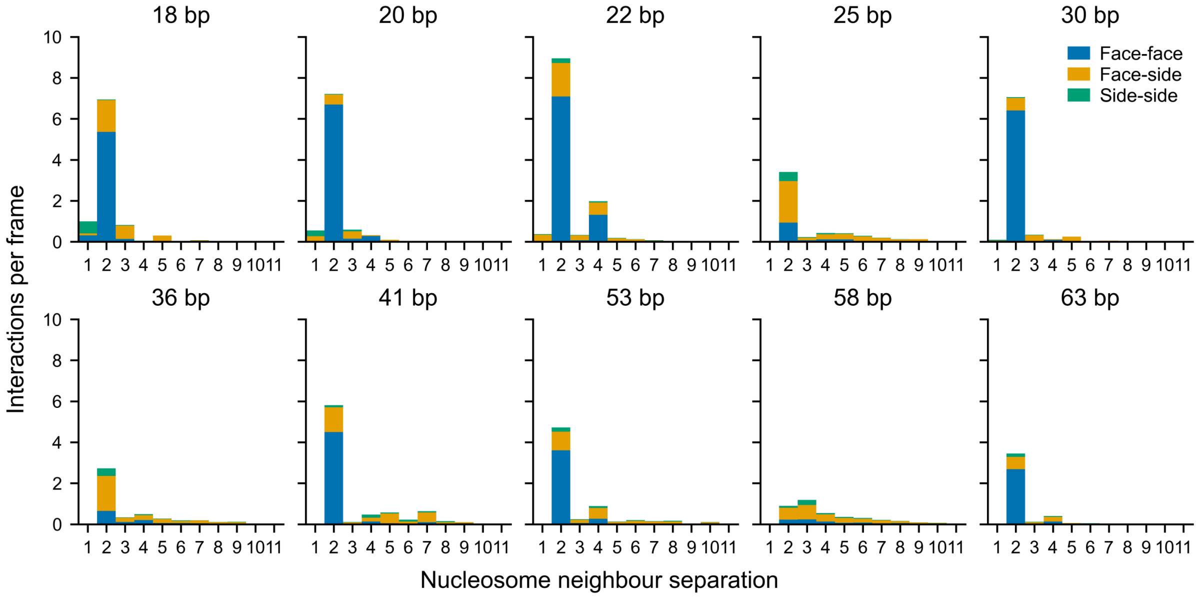


FIG. S9. **Per-frame nucleosome–nucleosome interaction counts across linker lengths.** Equivalent to main text Fig. 2c but showing absolute interaction counts per frame (not normalized per linker) to highlight the progressive reduction in inter-nucleosome contacts as linker length increases. Bars are decomposed by interaction type (face–face, face–side, side–side) and neighbor order ($i \pm 2$, $i \pm 3$, $i \pm 4$, etc.). The absolute counts reveal that while the relative proportions of interaction types change with the $10N$ versus $10N{+}5$ periodicity (main text Fig. 2c), the total number of contacts decreases with linker length due to increased DNA–DNA electrostatic repulsion and reduced fiber compaction.

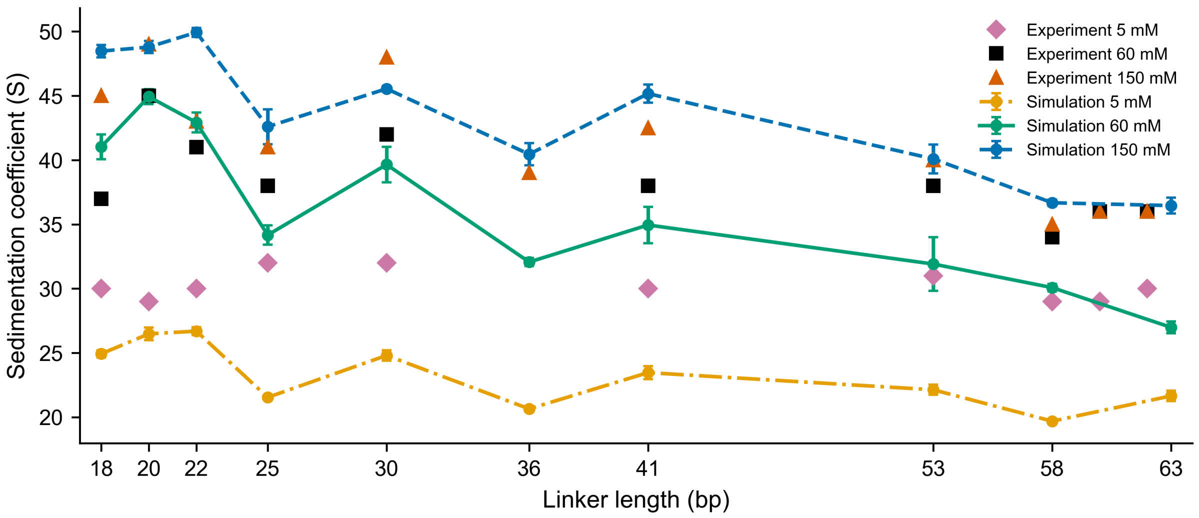


FIG. S10. **Linker-length dependence of chromatin-fibre compaction extended to 5 mM salt.** Sedimentation coefficients for 12-nucleosome arrays are shown as a function of linker length at 5, 60, and 150 mM monovalent salt. Experimental values at 60 and 150 mM are shown for comparison (black squares and orange triangles, respectively) [10, 11]. The 5 mM simulations used the same 12-nucleosome array setup and linker-length sweep as the 60 and 150 mM simulations, but with $\lambda_D \approx 4.36$ nm, corresponding to an implicit ionic strength of approximately 5.06 mM. Sedimentation coefficients were calculated from the final 2.5 $\mu$s of each 5 $\mu$s regular MD simulation, using 10,000 frames. Uncertainties were estimated as the standard deviation across five regular, non-overlapping trajectory blocks. The larger deviation from experiment at low salt and longer linker lengths is consistent with the limitations of the Debye–Hückel treatment discussed in the main text.

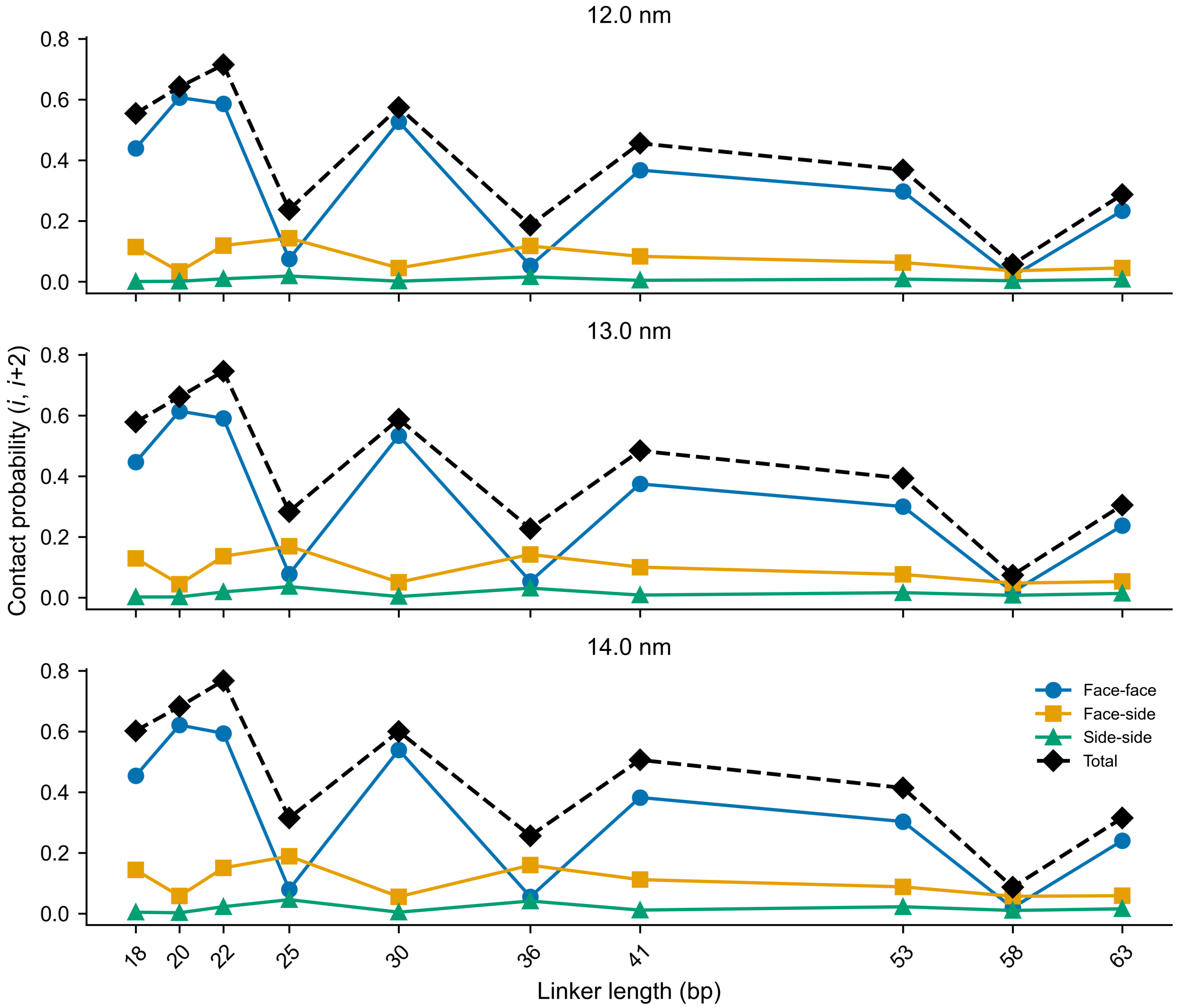


FIG. S11. **Cutoff-robustness of the nucleosome-level contact decomposition.** Linker-length dependence of inter-nucleosome contact probabilities for second-nearest-neighbour $(i, i{+}2)$ pairs, decomposed into face–face, face–side, side–side and total contributions (the same quantities plotted in main text Fig. 2b), evaluated at three centre-to-centre distance cutoffs of 12.0, 13.0 and 14.0 nm (top, middle, and bottom panels, respectively). The 13.0 nm cutoff is the one used in the main text. The $10N$ versus $10N{+}5$ oscillation pattern, the dominance of face–face contacts at $10N$ linkers, and the rank ordering of contact types are all preserved across the cutoff variation, demonstrating that the contact-decomposition trends reported in main text Fig. 2b are not artifacts of a single cutoff choice.

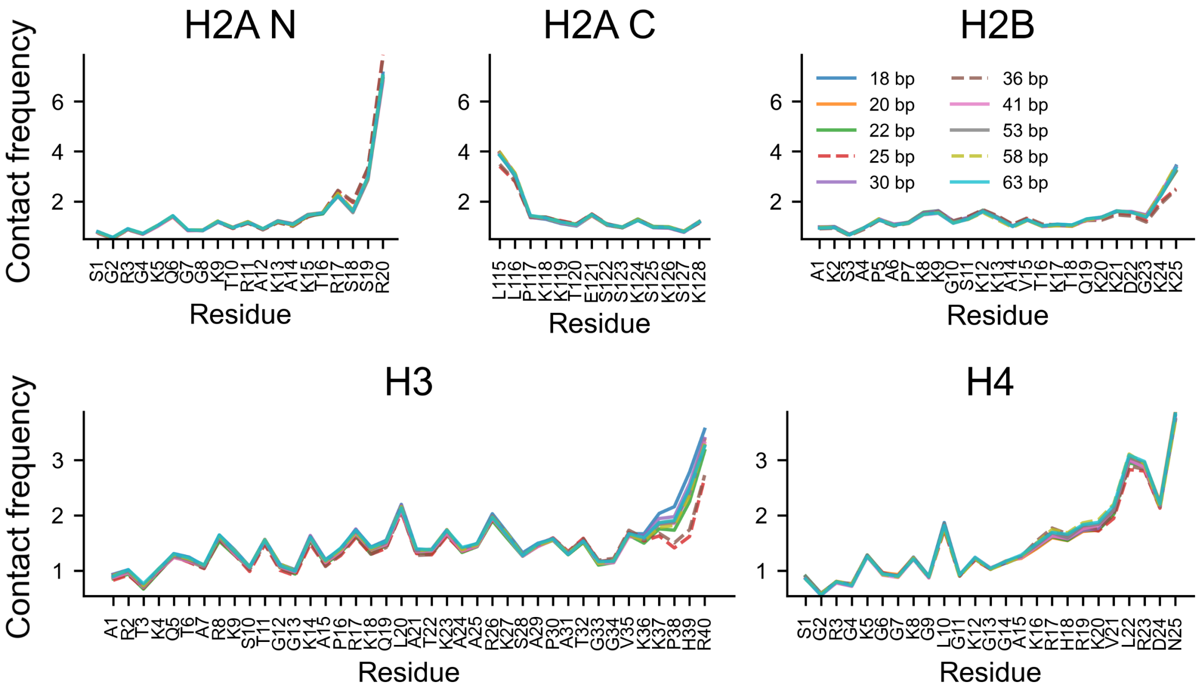


FIG. S12. **Intra-nucleosomal histone-tail contacts show little linker-length dependence.** Residue-level contact frequencies between histone-tail residues and their parental nucleosome for the 12-nucleosome fibres analysed in main text Fig. 3. Profiles overlap closely across the 18–63 bp linker-length sweep, unlike the inter-nucleosomal contacts shown in main text Fig. 3a.

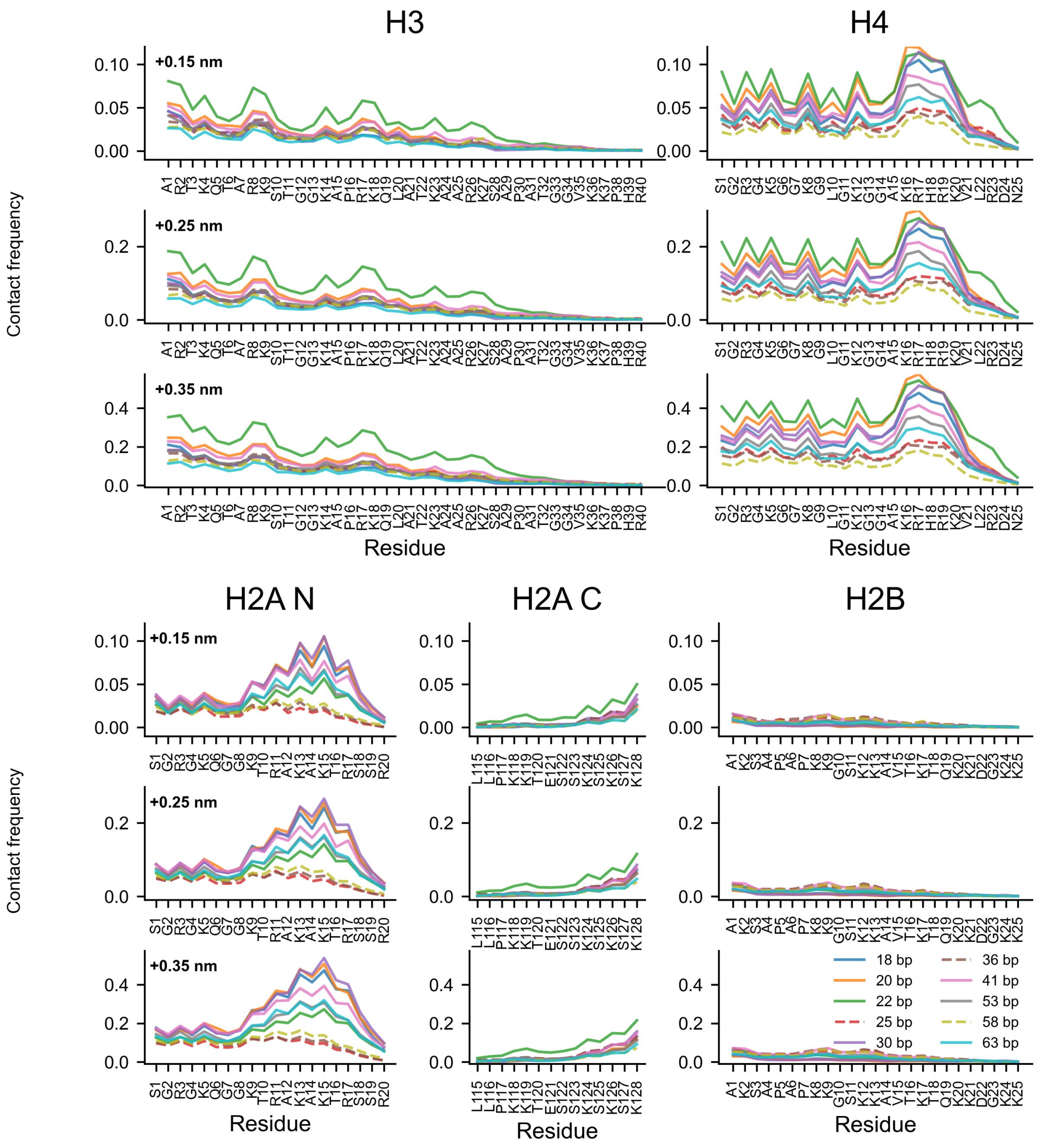


FIG. S13. **Residue-level histone-tail contact trends are robust to the contact cutoff.** Inter-nucleosomal residue-level contact frequencies, excluding the parental nucleosome, are shown for each histone tail across linker lengths of 18–63 bp. Contact cutoffs were defined using distance offsets of +0.15, +0.25, and +0.35 nm relative to $\frac{1}{2}(\sigma_i + \sigma_j)$. The qualitative contact profiles and linker-length-dependent trends are preserved across the three cutoff choices.

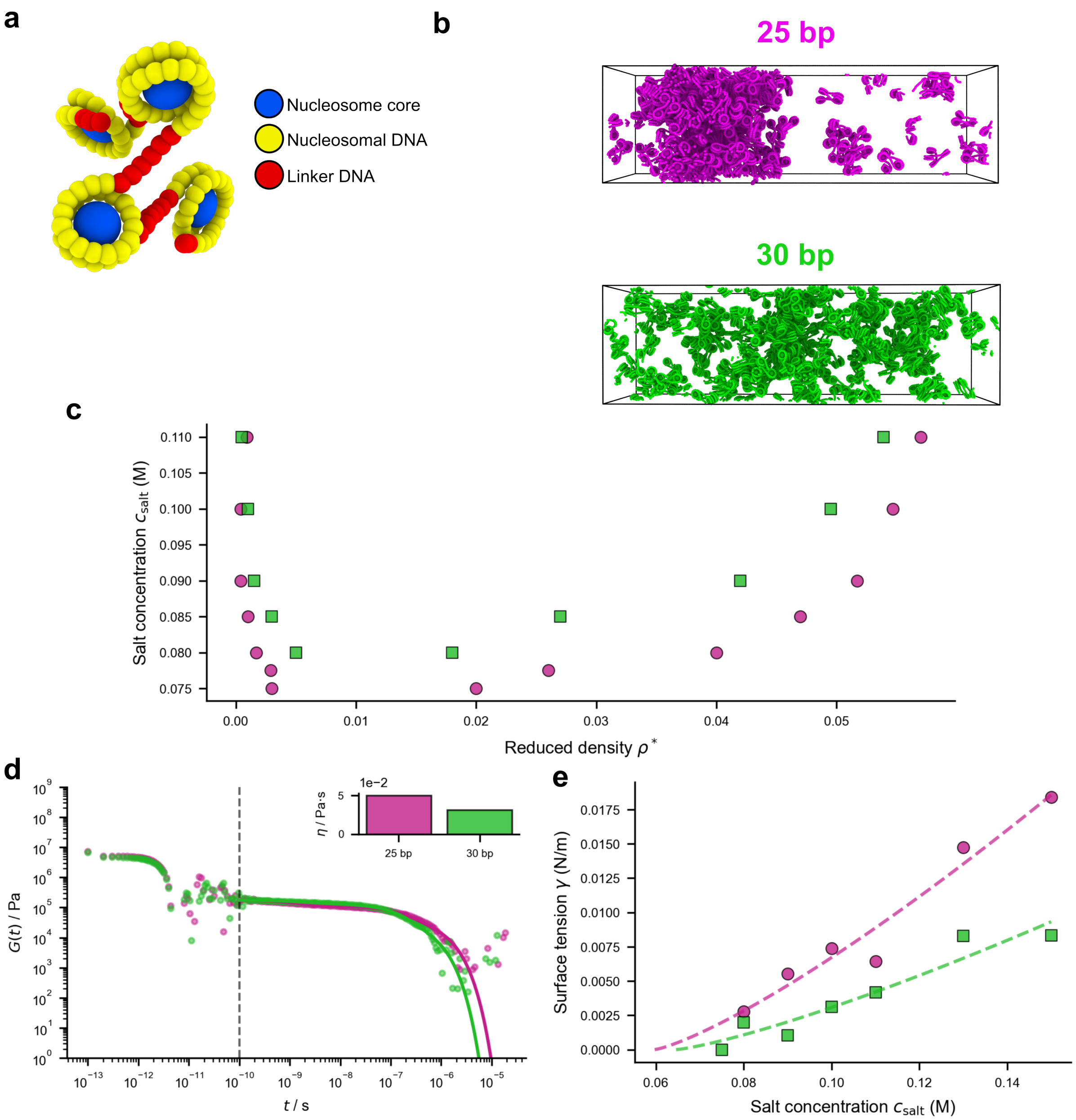


FIG. S14. **Minimal coarse-grained simulations reveal rheological features of chromatin condensates.** (a) Schematic illustrating the minimal coarse-grained chromatin model introduced by Farr and colleagues [1], in which each nucleosome is represented as a single anisotropic particle (1 bead per histone core, 1 ellipsoid per 5 bp DNA) with orientation-dependent interactions. (b) Representative simulation snapshots at 80 mM implicit salt concentration from direct coexistence simulations of tetranucleosome arrays with 25 and 30 bp linkers, showing distinct phase separation behavior. (c) Binodal phase diagrams for 25 and 30 bp chromatin as a function of salt concentration, demonstrating that 25 bp chromatin forms more stable condensates (phase separates at higher salt concentrations) than 30 bp chromatin. (d) Shear stress relaxation modulus, $G(t)$, as a function of time for 25 and 30 bp chromatin condensates at 150 mM salt, revealing similar short-time behavior but higher long-time relaxation modulus for 25 bp condensates. Inset bar chart indicates zero-shear viscosity obtained by integrating $G(t)$. The vertical line separates the time regime where direct numerical integration was used from the regime evaluated by fitting to the first three Maxwell modes. (e) Surface tension plotted against salt concentration for 25 and 30 bp chromatin condensates, with fits to the scaling form $\gamma = A(c_s - c_{s,c})^{1.26}$ [22], where $c_{s,c}$ is the critical salt concentration. The 25 bp condensates exhibit higher surface tension, consistent with stronger intermolecular interactions.

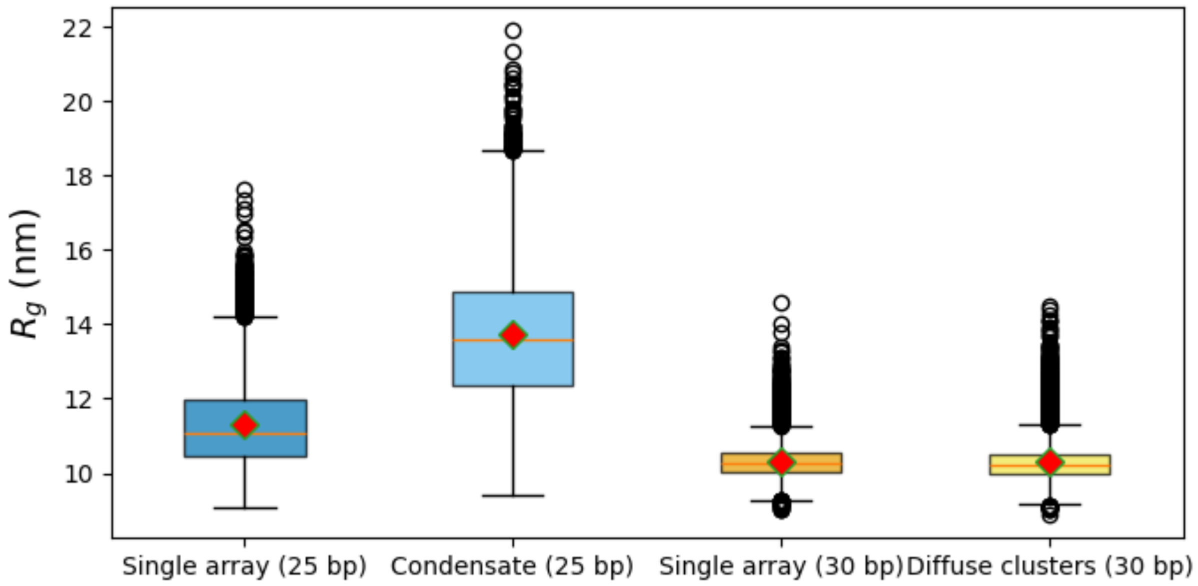


FIG. S15. **Tetranucleosome radius of gyration inside versus outside condensates.** Radius of gyration ($R_g$) distributions for (a) 25 bp and (b) 30 bp tetranucleosome arrays measured from the same conformational ensembles analyzed in main text Fig. 5: dilute phase (HREX simulations at 150 mM salt) and condensed phase (direct coexistence simulations at 100 mM salt). The 25 bp arrays (a) show a modest increase in mean $R_g$ upon phase separation (condensed > dilute), consistent with conformational expansion driven by increased inter-array interactions. In contrast, 30 bp arrays (b) maintain similar $R_g$ distributions in both conditions, reflecting their rigid zigzag conformations. The $R_g$ metric is highly correlated with the first principal component (PC1) from the structural clustering analysis in main text Fig. 5, confirming that PC1 primarily captures the compaction state of the arrays.

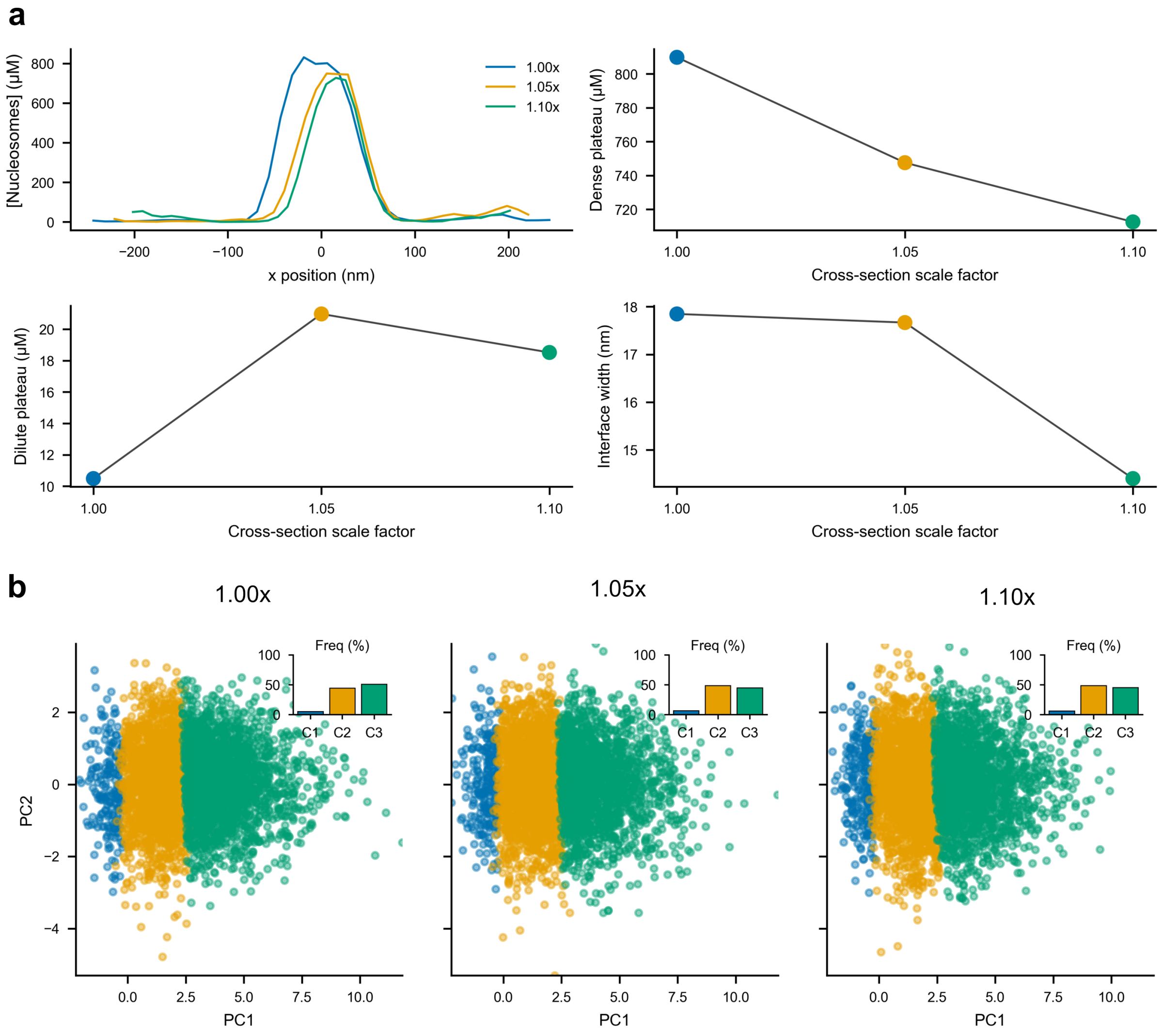


FIG. S16. **Direct-coexistence density profiles and structural clustering are insensitive to modest changes in box cross section.** The dense phase forms a slab-like domain within an elongated periodic simulation box, producing two dense–dilute interfaces along the long box axis. Starting from an equilibrated 25 bp direct-coexistence configuration in the default box (1.00×; Table S2), the transverse box dimensions were increased by scale factors of 1.05× and 1.10×, while the long-axis length was adjusted to keep the total volume constant. (a) Nucleosome density profiles along the long box axis for the three cross-section scale factors, with fitted dense-plateau density, dilute-plateau density, and interface width. Profiles were calculated from 9 μs trajectories after discarding the first 4 μs as burn-in, using one frame every 0.1 μs from the remaining 5 μs ($n = 50$ frames). The fitted quantities vary only modestly across the tested cross sections, indicating that the coexistence profile is not strongly biased by the default transverse box dimensions. (b) Structural-clustering projections for the same 50 frames from each trajectory onto the principal-component axes derived from the default-box ensemble, with cluster-population insets. Cluster definitions are the same as in main text Fig. 5. Similar cluster populations are obtained across the tested cross sections, indicating that the structural analysis is also insensitive to these modest changes in box geometry.

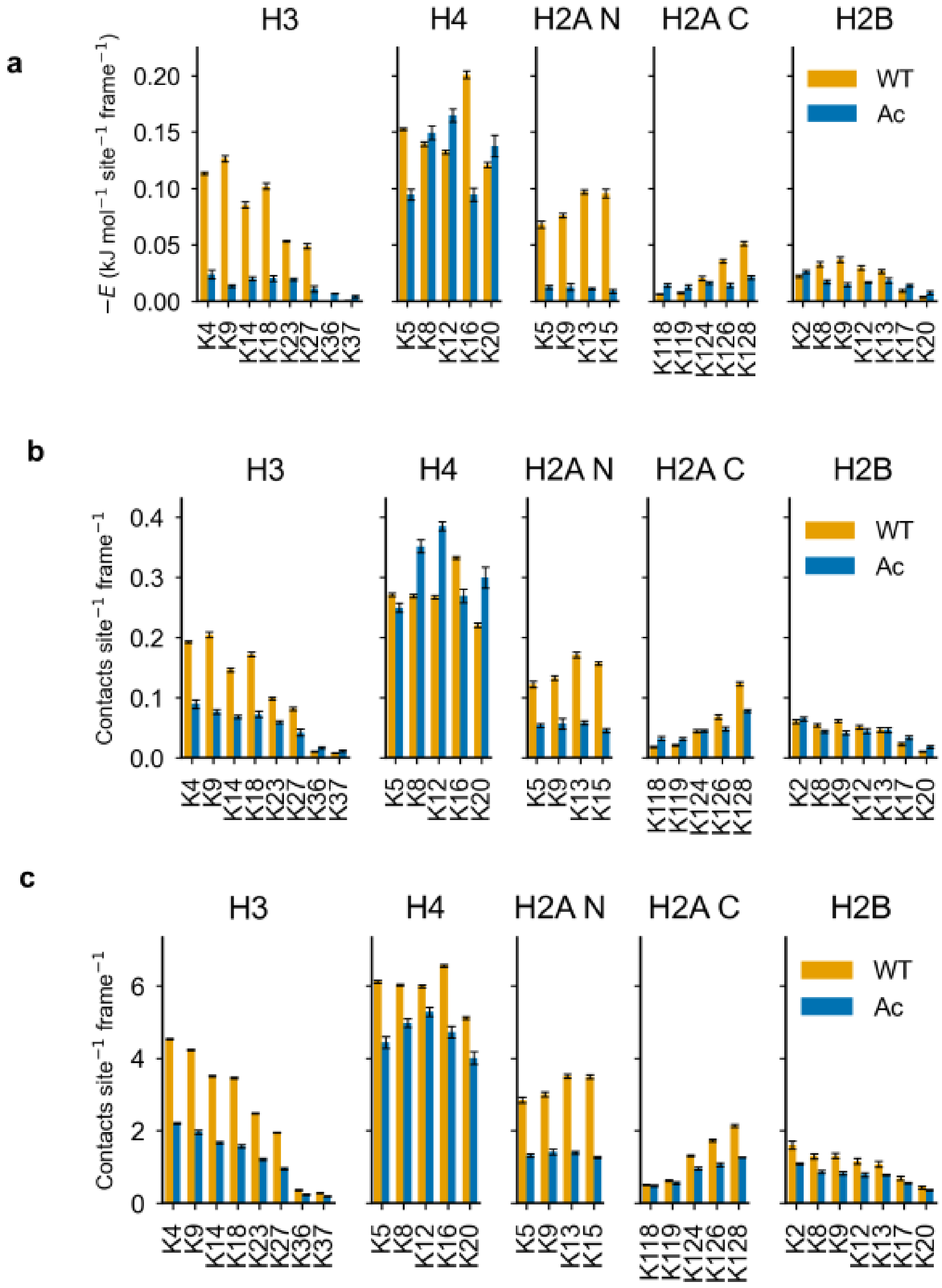


FIG. S17. **Close-contact and raw-frequency controls for the acetylation energy analysis.** Per-site lysine contact contributions for the 108-nucleosome acetylation simulations, aggregated across the 0 Ac, 5×12 Ac and 54 Ac systems and separated by lysine acetylation state (WT or Ac). (a) Full-energy lysine interaction contributions evaluated using the standard close-contact definition, $r < r_c = \frac{1}{2}(\sigma_i + \sigma_j) + 0.25$ nm. (b) Unweighted close-contact frequencies using the same $r < r_c = \frac{1}{2}(\sigma_i + \sigma_j) + 0.25$ nm cutoff. (c) Unweighted Debye-range contact frequencies using the $r < 1.6$ nm cutoff used for the main acetylation full-energy analysis. Together, these controls show that the main-text Debye-range full-energy result is not simply a consequence of changing the contact cutoff or of raw contact abundance alone. Bars show means over five analysis blocks; error bars show block standard deviations.

[1] S. E. Farr, E. J. Woods, J. A. Joseph, A. Garaizar, and R. Collepardo-Guevara, Nat Commun **12**, 2883 (2021).
[2] A. Sridhar, S. E. Farr, G. Portella, T. Schlick, M. Orozco, and R. Collepardo-Guevara, Proceedings of the National Academy of Sciences **117**, 7216 (2020).
[3] D. Farré-Gil, J. P. Arcon, C. A. Laughton, and M. Orozco, Nucleic Acids Research **52**, 6791 (2024).
[4] C. A. Davey, D. F. Sargent, K. Luger, A. W. Maeder, and T. J. Richmond, Journal of Molecular Biology **319**, 1097 (2002).
[5] K. Luger, A. W. Mäder, R. K. Richmond, D. F. Sargent, and T. J. Richmond, Nature **389**, 251 (1997).
[6] J. Anderson and J. Widom, Journal of Molecular Biology **296**, 979 (2000).
[7] Y. C. Kim and G. Hummer, Journal of Molecular Biology **375**, 1416 (2008).
[8] L. Chen, M. J. Maristany, S. E. Farr, J. Luo, B. A. Gibson, L. K. Doolittle, J. R. Espinosa, J. Huertas, S. Redding, R. Collepardo-Guevara, and M. K. Rosen, Nat Commun **16**, 6315 (2025).
[9] H. Zhou, J. Huertas, M. J. Maristany, K. Russell, J. H. Hwang, R.-w. Yao, J. Hutchings, M. Shiozaki, X. Zhao, L. K. Doolittle, B. A. Gibson, M. Riggi, J. R. Espinosa, Z. Yu, E. Villa, R. Collepardo-Guevara, and M. K. Rosen, Multi-scale structure of chromatin condensates rationalizes phase separation and material properties (2025).
[10] S. J. Correll, M. H. Schubert, and S. A. Grigoryev, The EMBO Journal **31**, 2416 (2012).
[11] M. V. Bass, T. Nikitina, D. Norouzi, V. B. Zhurkin, and S. A. Grigoryev, Journal of Biological Chemistry **294**, 4233 (2019).
[12] P. D. Dans, A. Balaceanu, M. Pasi, A. S. Patelli, D. Petkevičiūtė, J. Walther, A. Hospital, G. Bayarri, R. Lavery, J. H. Maddocks, and M. Orozco, Nucleic Acids Research **47**, 11090 (2019).
[13] Z. Zhang, X. Liu, K. Yan, M. E. Tuckerman, and J. Liu, J. Phys. Chem. A **123**, 6056 (2019).
[14] J. D. Chodera and M. R. Shirts, The Journal of Chemical Physics **135**, 194110 (2011).
[15] X. Ding, X. Lin, and B. Zhang, Nature Communications **12**, 1091 (2021).
[16] W. Lu, C. Bueno, N. P. Schafer, J. Moller, S. Jin, X. Chen, M. Chen, X. Gu, A. Davtyan, J. J. De Pablo, and P. G. Wolynes, PLoS Comput Biol **17**, e1008308 (2021).
[17] P. Eastman, J. Swails, J. D. Chodera, R. T. McGibbon, Y. Zhao, K. A. Beauchamp, L.-P. Wang, A. C. Simmonett, M. P. Harrigan, C. D. Stern, R. P. Wiewiora, B. R. Brooks, and V. S. Pande, PLoS Comput Biol **13**, e1005659 (2017).
[18] P. Eastman, R. Galvelis, R. P. Peláez, C. R. A. Abreu, S. E. Farr, E. Gallicchio, A. Gorenko, M. M. Henry, F. Hu, J. Huang, A. Krämer, J. Michel, J. A. Mitchell, V. S. Pande, J. P. Rodrigues, J. Rodriguez-Guerra, A. C. Simmonett, S. Singh, J. Swails, P. Turner, Y. Wang, I. Zhang, J. D. Chodera, G. De Fabritiis, and T. E. Markland, J. Phys. Chem. B **128**, 109 (2024).
[19] W. Humphrey, A. Dalke, and K. Schulten, Journal of Molecular Graphics **14**, 33 (1996).
[20] A. R. Tejedor, R. Collepardo-Guevara, J. Ramírez, and J. R. Espinosa, J. Phys. Chem. B **127**, 4441 (2023).
[21] C. Phillips, *Dissecting the Architectural Properties of Chromatin and the Influence of the H1 Linker Histone on Chromatin Regulation*, Ph.D. thesis, Apollo - University of Cambridge Repository (2024).
[22] J. S. Rowlinson and B. Widom, *Molecular theory of capillarity* (Courier Corporation, 2013).
